# Faecal microbiota transplantation from Alzheimer’s participants induces impairments in neurogenesis and cognitive behaviours in rats

**DOI:** 10.1101/2022.11.04.515189

**Authors:** Stefanie Grabrucker, Moira Marizzoni, Edina Silajdžić, Nicola Lopizzo, Elisa Mombelli, Sarah Nicolas, Sebastian Dohm-Hansen, Catia Scassellati, Davide Vito Moretti, Melissa Rosa, Karina Hoffmann, Jane A English, Aonghus Lavelle, Cora O’Neill, Sandrine Thuret, Annamaria Cattaneo, Yvonne M Nolan

## Abstract

The gut microbiome is emerging as an important susceptibility factor in Alzheimer’s disease (AD) possibly due to the increased prevalence of pro-inflammatory genera in gut microbiota of AD participants. Microbiota-mediated changes in cognition and adult hippocampal neurogenesis (AHN), an important process for memory which is altered in AD, position the microbiota-gut-brain axis as a key regulator of AD. However, it is unknown whether gut microbiota alterations are the cause or consequence of AD symptoms. We transplanted faecal microbiota from AD participants and age-matched controls into microbiota-depleted naïve adult rats and found impairments in AHN and associated memory tasks, which correlated with clinical cognitive scores. Discrete changes in the rat caecal and hippocampal metabolome were evident. Serum from AD participants also decreased neurogenesis in vitro and correlated with cognitive scores and pro-inflammatory genera. Our results reveal that the cognitive symptoms in AD may be due to alterations in gut microbiota, and that impaired neurogenesis may be a mechanistic link between altered gut microbiota and cognitive impairment in AD.

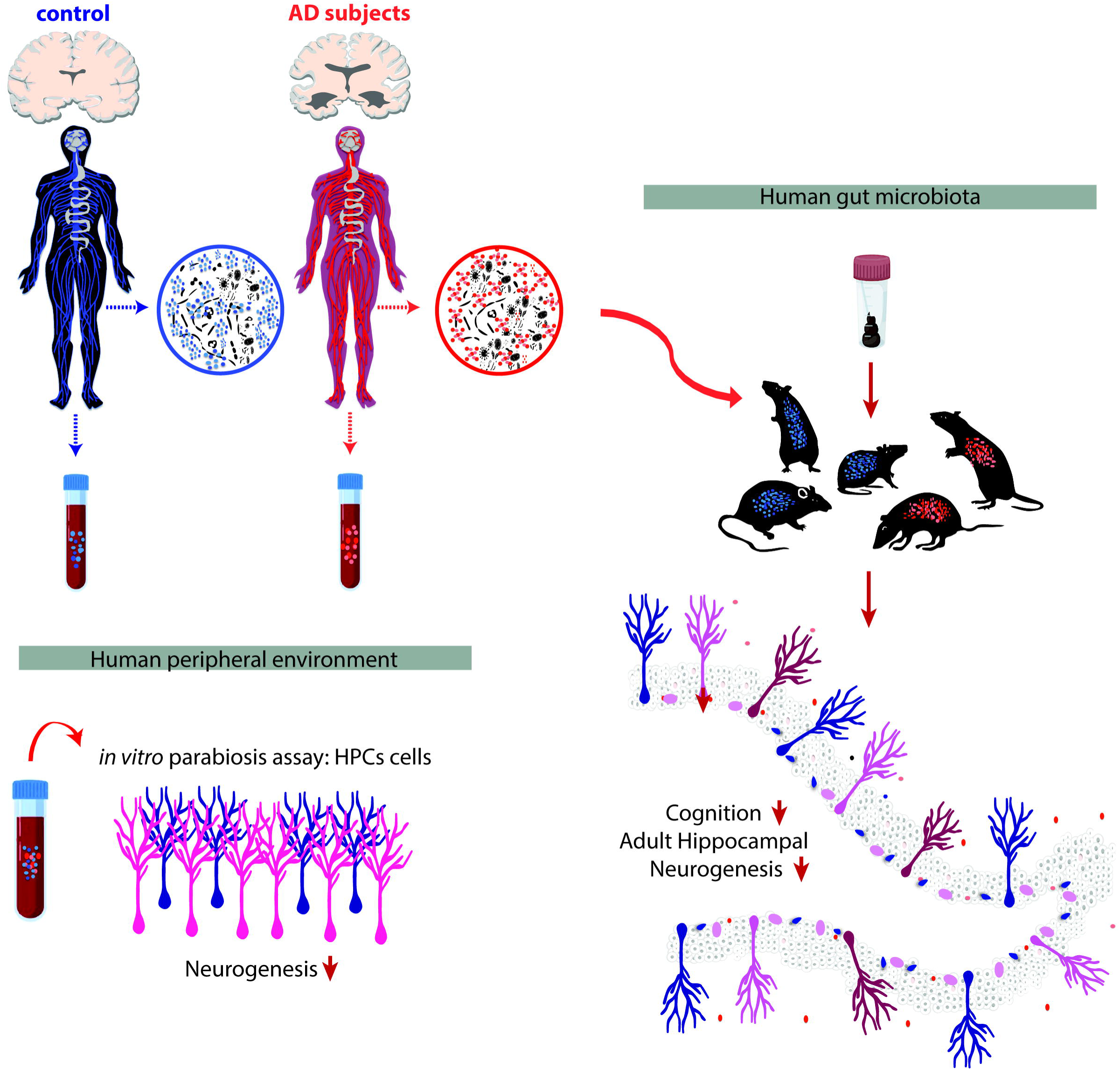

---

Alzheimer’s disease (AD), the leading cause of dementia, is a complex neurodegenerative disorder characterised by progressive impairments in memory, cognitive function, and behaviour (“2016 Alzheimer’s disease facts and figures.,” 2016; Blennow et al., 2006). Neuropathological hallmarks of AD include extracellular deposition of amyloid-β (Aβ) plaques, gradual intraneuronal accumulation of neurofibrillary tangles made up of hyperphosphorylated cytoskeletal tau protein, neuroinflammation and neuronal death in specific brain regions (Heneka et al., 2015; Iqbal et al., 2005; Long and Holtzman, 2019; Selkoe and Hardy, 2016). The hippocampus plays a critical role in learning and memory (Squire, 1992) and is particularly vulnerable to AD pathology, being one of the earliest brain areas to be affected (Braak et al., 1993). It is becoming increasingly recognised that AD is a multifactorial disease, substantially influenced by genetic, lifestyle and environmental factors (Bellenguez et al., 2022; Bertram and Tanzi, 2008; Jansen et al., 2019; Kivipelto et al., 2018; Lambert et al., 2013; Robinson et al., 2017; Tanzi and Bertram, 2005), that may result in common brain pathology (Gong et al., 2018; Veitch et al., 2019).

The gut microbiome is emerging as a key susceptibility factor and a target for investigation in AD due to research demonstrating altered microbial composition in AD (Cattaneo et al., 2017; Guo et al., 2021; Haran et al., 2019; Li et al., 2019; Ling et al., 2020b, 2020a; Liu et al., 2019; Vogt et al., 2017; Zhuang et al., 2018) and the presence of bacterial endotoxins, which contribute to inflammatory processes (Asti and Gioglio, 2014; Zhan et al., 2018, 2016). Indeed, it is hypothesised that Aβ has anti-microbial functions and that infectious or sterile inflammatory stimuli may drive amyloidosis. Preclinical studies showing the contribution of gut microbiota to age-related cognitive changes have bolstered the concept of altered microbial composition contributing to the symptomology of AD (Li et al., 2020; Minter et al., 2016; Rei et al., 2022; Varesi et al., 2022). Moreover, a recent study using statistical genetic approaches demonstrated a positive significant genetic overlap and correlation between AD and gastrointestinal tract disorders (Adewuyi et al., 2022).

Alterations in gut microbiota have also been shown to modulate adult hippocampal neurogenesis (AHN), an important process for learning and memory (Cruz-Pereira et al., 2020; Deng et al., 2010; Gonçalves et al., 2016), which declines with age (Heine et al., 2004) and is altered early in AD (Choi and Tanzi, 2019; Moreno-Jimenez et al., 2019; Tobin et al., 2019). AHN is a key process for cognitive functions such as spatial learning and pattern separation, an ability to distinguish between highly similar events or environments (Clelland et al., 2009; Garthe and Kempermann, 2013; Sahay et al., 2011), both of which are impaired during ageing (Holden and Gilbert, 2012; Yassa and Stark, 2011) and in AD (Ally et al., 2013; Parizkova et al., 2020). Crucially, AHN is altered in early Braak stages, preceding neurofibrillary tangles and amyloid β plaque formation in the hippocampus (Moreno-Jimenez et al., 2019), suggesting that AHN dysfunction is an early feature of AD pathogenesis. Transplantation of gut microbiota derived from a 5xFAD mouse model of AD has been shown to impair memory function and AHN in C57BL/6 mice (Kim et al., 2021). It has also been shown that a decrease in newly-generated neurons in the dentate gyrus of people with advancing Braak stages is accompanied by changes in inflammatory cell load (Ekonomou et al., 2015). In addition, AHN is significantly reduced by inflammation, which is central to neuropathological developments in AD (Ryan et al., 2013).

While studies to date implicate the microbiome-gut-brain axis in AD, it is yet unknown whether alterations in the intestinal microbiota composition reported in AD participants are a cause or consequence of symptoms, metabolic, neuroplasticity and inflammatory changes in these individuals. We addressed this using a novel approach combining characterisation of a gut microbiota signature in a cohort of control and AD subjects, back translation using faecal microbiota transplantation from AD participants to rat to determine behavioural, neurogenic and inflammatory features mediated through gut microbiota, and an *in vitro* neurogenesis assay using participant serum to assess how the systemic milieu of AD participants contributes to changes in hippocampal neurogenesis.

## RESULTS

### Individuals with AD show systemic inflammation and gut microbiota dysbiosis

Recent studies have described a potential link between the development of AD pathologies and alterations in the intestinal microbial community (Liu et al., 2020). Dysregulation in peripheral inflammatory responses has been associated with the aetiology and progression of the disease (Bettcher et al., 2021). However, there is little evidence showing that alterations in gut microbiota composition are associated with local and systemic peripheral inflammation in AD participants. Therefore, we first characterised the peripheral inflammatory status and gut microbial composition of control subjects and AD participants. To that end, RNA was extracted from the whole blood of 69 healthy control subjects and 64 participants diagnosed with AD (Fig. 1A) and the expression of pro-inflammatory cytokines and inflammatory marker genes was analysed. Our results show that IL-1β, the inflammasome marker NLRP3 and MIF were significantly upregulated in AD participants, indicating the presence of systemic inflammation in AD participants (Fig. 1B). To investigate the intestinal inflammatory status, we characterised the inflammatory protein calprotectin in the faeces of control and AD participants, which has been shown to correlate with the presence and severity of intestinal inflammation (Bjarnason, 2017). Notably, faecal calprotectin levels were significantly increased in AD participants compared to control subjects (Fig. 1C). To validate whether increased inflammation in the intestine was associated with an alteration of gut microbiota composition, bacterial 16S rRNA gene sequencing was carried out in a subset of AD participants and controls to identify their faecal microbiota composition (Extended Data Fig. 1A-E). No significant change in alpha and beta diversities was detected between control subjects and AD participants (Extended Data Fig. 1B-D). However, at phylum level, AD participants were characterised by a higher abundance of Bacteroidetes (Fig. 1D) reported to comprise many pro-inflammatory species (Nomura et al., 2021), and a lower abundance of the phyla Firmicutes and Verruocomicrobiota, shown to contain species that can act as anti-inflammatory (Stojanov et al., 2020). At the genus level, AD participants demonstrated on average a significant reduction in the relative abundance of the anti-inflammatory short chain fatty acid butyrate-producing genera *Coprococcus* (Slingerland et al., 2017) and *Clostridium sensu stricto 1* (Fig. 1D). In addition, similar to a previous report using a transgenic mouse model of AD (Chen et al., 2020), the relative abundance of the inflammation-promoting genera *Desulfovibrio* was significantly increased in AD participants compared to cognitively healthy control subjects (Fig. 1D). These results confirmed that the observed alteration in phylum and genus levels of gut microbiota composition may be linked to intestinal inflammation in AD (Goyal et al., 2021).

**Figure 1:**
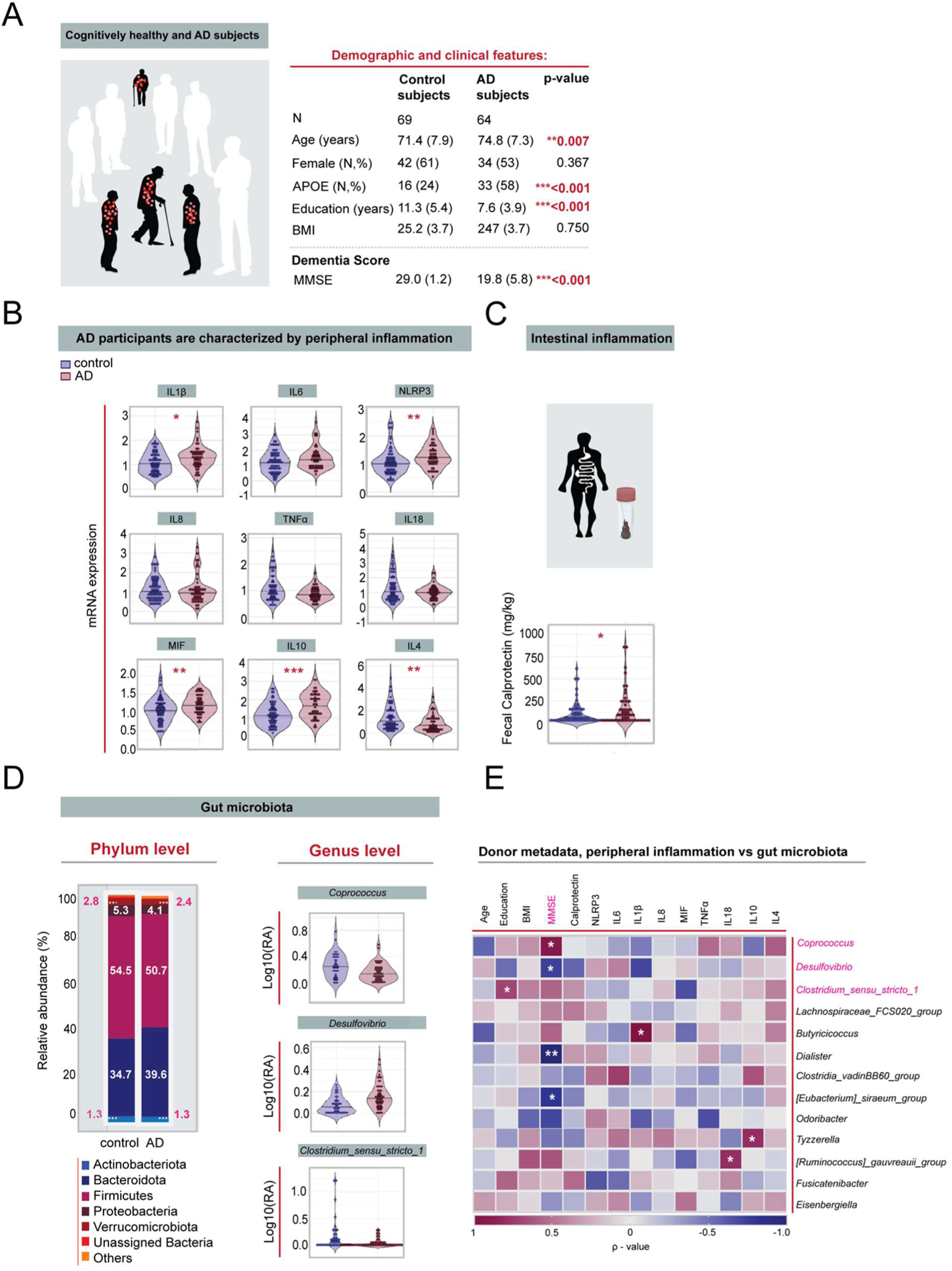
Characterization of peripheral inflammation and gut microbiota dysbiosis in Alzheimer’s disease participants. **(A)** Demographic and clinical features of control subjects (n=69) and AD subjects (n=64), p-values were calculated using unpaired, two-tailed Student’s t-test for continuous Gaussian variables (or Mann-Whitney test for non-Gaussian variables) and Chi-square test for categorical data. There were 8 missing values for APOE genotype, 6 missing values for MMSE and BMI. AD subjects were significantly older (\*\**p*= 0.007), showed a significant increase in the prevalence of APOE carriers (\*\*\**p*< 0.001), were less educated (\*\*\**p*<0.001) and had lower MMSE score (\*\*\**p*<0.001) than cognitively healthy controls. **(B)** There was increased expression of the cytokines IL-1β (\**p*=0.018), NLRP3 (\*\**p*=0.003), MIF (\*\**p*=0.008), IL-10 (\*\*\**p*<0.001), and decreased expression of IL-4 (\*\**p*=0.006) in AD subjects (n=52) compared to control subjects (n=68), unpaired, two-tailed Student’s t-test for continuous Gaussian variables (or Mann-Whitney test for non-Gaussian variables). **(C)** Faecal calprotectin was significantly increased in AD subjects (n=64) compared to control subjects (n=69), Mann-Whitney test, \**p*=0.022. **(D)** Gut microbiota composition at phylum level from control (n=41) and AD subjects (n=54). AD subjects are characterized by a higher abundance of pro-inflammatory phyla Bacteroides, and lower abundance of anti-inflammatory phyla Firmicutes and Verrucomicrobiota. Relative abundance of genera that differed significantly between controls and AD subjects after batch correction using percentile-normalization. Mann-Whitney tests, multiple testing corrections using the Benjamini–Hochberg method, and FDR ≤ 0.1 was considered significant (*Coprococcus*, *p*=0.099; *Clostridium in sensu stricto 1*, *p*=0.099; *Desulfovibrio*, *p* =0.006). **(E)** Correlation between human gut microbiota, human donor metadata, faecal calprotectin and serum inflammatory markers. Heatmap shows Spearman rank coefficients, with red indicating strong positive correlation, and blue indicating strong negative correlation. *P* values for significant correlations (α<0.05) are noted (\**p*<0.05, \*\**p*<0.01, \*\*\**p*<0.001). Black horizontal lines in violin plots indicate medians. All data are presented as mean ± SEM, \**p*<0.05, \*\**p*<0.01, \*\*\**p*<0.001, NS, not significant. Abbreviations: (BMI) Body Mass Index calculated as weight/height^2^ and measured in kg/cm^2^, (MMSE) Mini Mental State Examination, (AD) Alzheimer’s Disease, (APOE) Apolipoprotein E.

To further understand whether the detected peripheral alterations are associated with the clinical phenotype of AD participants, we correlated Mini-Mental State Exam (MMSE) scores with the peripheral immune profile and with the gut microbiota alterations in our cohort. Notably, we observed significant associations between the MMSE score and the gut microbiota signature (Fig. 1E). Importantly, a positive correlation was observed between the abundance of *Coprococcus* and the MMSE score, indicating a relationship between the reduction in an anti-inflammatory bacteria and cognitive impairment. In line, inverse correlations were detected between the abundance of the inflammatory-related genera *Desulfovibrio, Dialister* and the MMSE score, further supporting a direct association between the prevalence of inflammatory associated gut microbiota and the cognitive performance of AD participants. In order to understand whether the microbial alterations are responsible for decreased cognitive performance, we next performed studies aimed at revealing the mechanisms underlying the correlation between altered gut microbiota composition and cognitive impairment.

### Transplantation of gut microbiota from individuals with AD caused intestinal dysfunction in rats

To elucidate the functional contribution of human gut microbiota to AD aetiology, we transplanted faecal samples from cognitively healthy subjects and AD participants into microbiota-depleted rats (Fig. 2A). For this purpose, human cognitively healthy and AD donors were selected based on clinical features, using the MMSE and Clinical Dementia Rating (CDR) (Fig. 2B). To validate donor engraftment and the temporal colonisation dynamics of gut microbiota transplantation throughout the study, we analysed baseline faecal samples using 16S rRNA gene sequencing, reassessed composition and diversity ten days after human donor colonization (D10) and again at the end of the study (D59; timeline, Fig. 2A).

**Figure 2:**
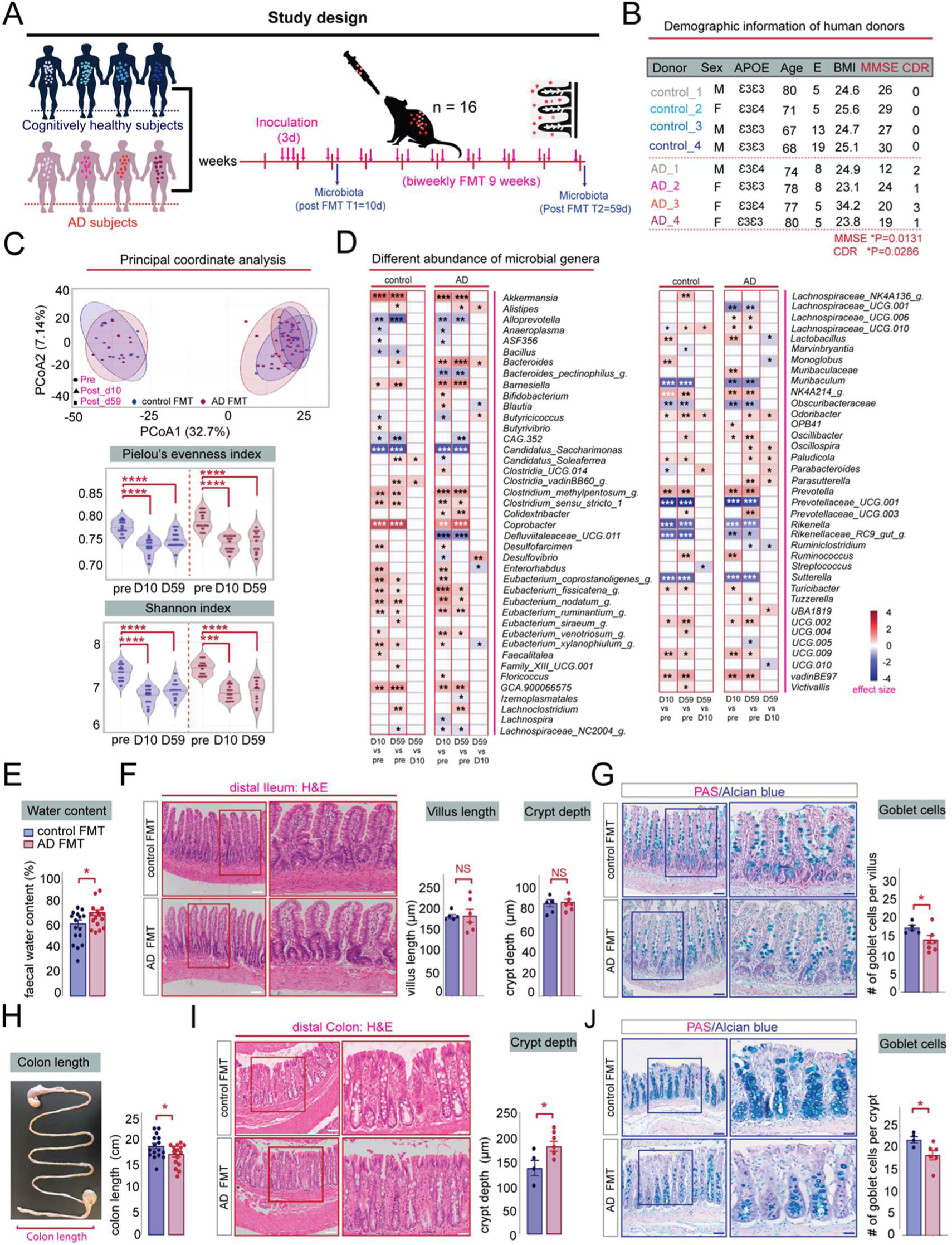
Rats colonized with faecal material from AD participants harbour different bacterial genera and display alterations in intestinal morphology and function. **(A)** Schematic representation of the experimental procedure: microbiota-depleted rats were colonized with human faecal samples from cognitively healthy control or AD donors (n=4). One faecal sample from each human donor was used to colonize 4 microbiota-depleted rats (n=16). Faecal pellets were collected at day 0, 10 days after human donor colonization (T1) and at the end of the study (T2= 59 days). Vertical bars represent weekly intervals. **(B)** Donor metadata: Demographic and clinical characteristics of human donors (n=4) used for rat colonization. MMSE and CDR scores were used to determine cognitive function applicable to dementia of human donors. Cognitive function was significantly impaired in human AD donors compared to control donors as measured by MMSE, unpaired, two-tailed Student’s t-test, \**p*=0.0131 and CDR scores, Mann-Whitney test, \**p*=0.0286. Human donors in both groups display no significant difference in age, unpaired, two-tailed Student’s t-test, *p*=0.8326; education level, Mann-Whitney test, *p*=0.6571 and BMI, Mann-Whitney test, *p*=0.6857. **(C)** Upper panel: Principal coordinate analysis showing the effects of FMT on faecal microbiome in rats in terms of β-diversity as measured by Bray-curtis distance. Ellipses indicate 95% confidence intervals per group. Linear mixed-effects model, significant time × FMT interaction effect (\**p*=0.047). Lower panel: Violin plots displaying the effects of FMT on rats in terms of α-diversity. Black horizontal lines in violin plots indicate medians. The Pielou’s evenness and Shannon index decreased following FMT, regardless of donor group, (ANOVA with repeated measures \**p* < 0.05, \*\*\**p* < 0.001, \*\*\*\**p* < 0.0001). **(D)** Left panel: Heatmap showing genera differentially altered by FMT. Colour depicts effect size, with blue (negative) indicating higher abundances pre-treatment and red (positive) indicating higher abundances post-treatment, Wilcoxon signed-rank test followed by Benjamini–Hochberg correction as per the ALDEx2 library, \**q* < 0.1, \*\**q* < 0.01, \*\*\**q* < 0.001. **(E)** AD FMT rats (n=16) display a significant increase in faecal water content compared to control FMT rats (n=16), unpaired, two-tailed Student’s t-test, \**p*=0.0309. **(F)** Left panel: Representative images of H&E-stained sections of the distal ileum from control and AD FMT rats. Scale bars: 200 µm left magnification, 100 µm right magnification. Right panel: AD colonized rats show no significant change in ileal villus length, unpaired, two-tailed Student’s t-test, *p*=0.7749 and crypt depth, unpaired, two-tailed Student’s t-test, *p*=0.9060. **(G)** Left panel: Representative PAS/Alcian blue staining of ileal sections. Scale bars: 200 µm left magnification, 100 µm right magnification. Right panel: AD FMT rats display significant goblet cell loss in the proximal colon, unpaired, two-tailed Student’s t-test, \**p*=0.0403. **(H)** Average colon length was significantly reduced in AD FMT rats compared to control FMT rats, two-tailed Student’s t-test, \**p*=0.0500. **(I)** Left panel: Representative images of H&E-stained sections of the proximal colon from control and AD FMT rats. Scale bars: 200 µm left magnification, 100 µm right magnification Right panel: AD FMT rats displayed a significant increase in colonic crypt depth, two-tailed Student’s t-test, \**p*=0.0356. **(J)** Left panel: Representative PAS/Alcian blue staining of colon sections. Right panel: AD FMT rats display a significant goblet cell loss in the proximal colon, unpaired, two-tailed Student’s t-test, \**p*=0.0403. All data are presented as mean ± SEM, \**p*<0.05, \*\**p*<0.01, \*\*\**p*<0.001, **** p<0.0001, NS, not significant. Abbreviations: (APOE) Apolipoprotein E, (E) Education, (BMI) Body Mass Index, (MMSE) Mini Mental State Examination, (CDR) Clinical Dementia Rating, (AD) Alzheimer’s Disease.

Principal coordinate analysis (PCoA) of the Bray-curtis distance matrix revealed that the temporal microbial changes were not influenced by the clinical status of the donors (Fig. 2C, upper panel). Additionally, a decrease in alpha diversity after human donor colonisation was observed in rats of both groups at both timepoints, indicating a loss of bacterial species (Fig. 2C, lower panels). These differences were independent from the clinical status of the donors. Fifty-nine days after FMT, a decrease in gained taxa was observed in rats that received faecal material from AD participants, suggesting that their microbiota tended to return to its original state (Extended Data Fig. 2C). On average, 40 percent of the taxa from human donors engrafted into recipient rats, without differences between rats that received FMT from cognitively healthy controls or AD (Extended Data Fig. 2D). Transplantation efficiency was lowest for taxa belonging to the phyla *Bacteroidota* and *Proteobacteria* (Extended Data Fig. 2F). Post-FMT, 78 genera were significantly altered in recipient rats (Fig. 2D). Compared with the original microbiota, human AD colonised rats displayed greater alterations in microbial genera than rats receiving control FMT at D10 (51 genera vs 41) but not at D59 (44 vs 45) (Fig. 2D).

Intestinal dysfunction, along with enteric inflammation and accumulation of AD-related proteins in intestinal tissue, has been described in genetic mouse models of AD (Chen et al., 2020; Honarpisheh et al., 2020; Pellegrini et al., 2020). In order to elucidate whether human AD gut microbiota actively participate in pathophysiological processes of the GI tract, we evaluated the general health and overall appearance of the rat intestine after FMT treatment. Our results show that human AD FMT had no significant impact on faecal pellet output or caecum weight of recipient rats (Extended Data Fig. 3A, B). Likewise, no general adverse effects on body weight composition or food intake were detected after FMT treatment in both groups (Extended Data Fig. 3C, D). Additionally, no significant differences were detected in the expression of genes encoding pro-inflammatory cytokines in colonic tissue of rats after FMT (Extended Data Fig. 4A-I). In line with this, the presence of cytokines in the systemic circulation was unaltered (Extended Data Fig. 5A, B), suggesting that no local inflammatory event occurred in the GI tract due to humanization with AD FMT. However, human AD colonised rats displayed a significant increase in faecal water content (Fig. 2E) and water intake (Extended Data Fig. 3E) along with a reduction in colon length (Fig. 2H), suggesting specific effects of the human AD gut microbiota on colonic function. These findings were further supported by structural changes in the depth of the colonic but not ileal crypts as determined by H&E staining (Fig. 2F, I). Compared to control colonised rats, AD recipient rats displayed crypt hyperplasia in the proximal colon, with the lengths of the crypt significantly elongated (Fig. 2I). Additionally, we found a reduction in the number of goblet cells in the colon and ileum, suggesting altered mucin production after transplantation with human AD gut microbiota (Fig. 2G, J). Collectively, our data demonstrate that the presence of human AD gut microbiota alters intestinal morphology and promotes intestinal dysfunction, which may increase the susceptibility to intestinal inflammation that we observed in AD participants.

### Human AD FMT induced cognitive deficits in hippocampal-neurogenesis dependent behaviours

To investigate whether alterations of the human gut microbiota contribute to the cause of the AD behavioral phenotype, we evaluated next the cognitive and non-cognitive performance of rats colonised with faecal material from human control subjects and AD participants. Rats were tested two weeks after initial human donor colonisation in a behavioural test battery to assess task-specific memory performance and other AD-related co-morbidities (Fig. 3A). Rats colonized with faecal material from AD donors exhibited no change in locomotor parameters in the open field test compared to rats colonised with material from control participants (Extended Data Fig. 6A). Additionally, we did not detect significant changes in anxiety-related behaviors in the Elevated Plus Maze (EPM) test (Extended Data Fig. 6B), nor were there significant alterations in antidepressant-like behavior in the Forced Swim test (FST) (Extended Data Fig. 6C), indicating no specific effects of the human gut microbiota on co-morbid features of AD in rats.

**Figure 3:**
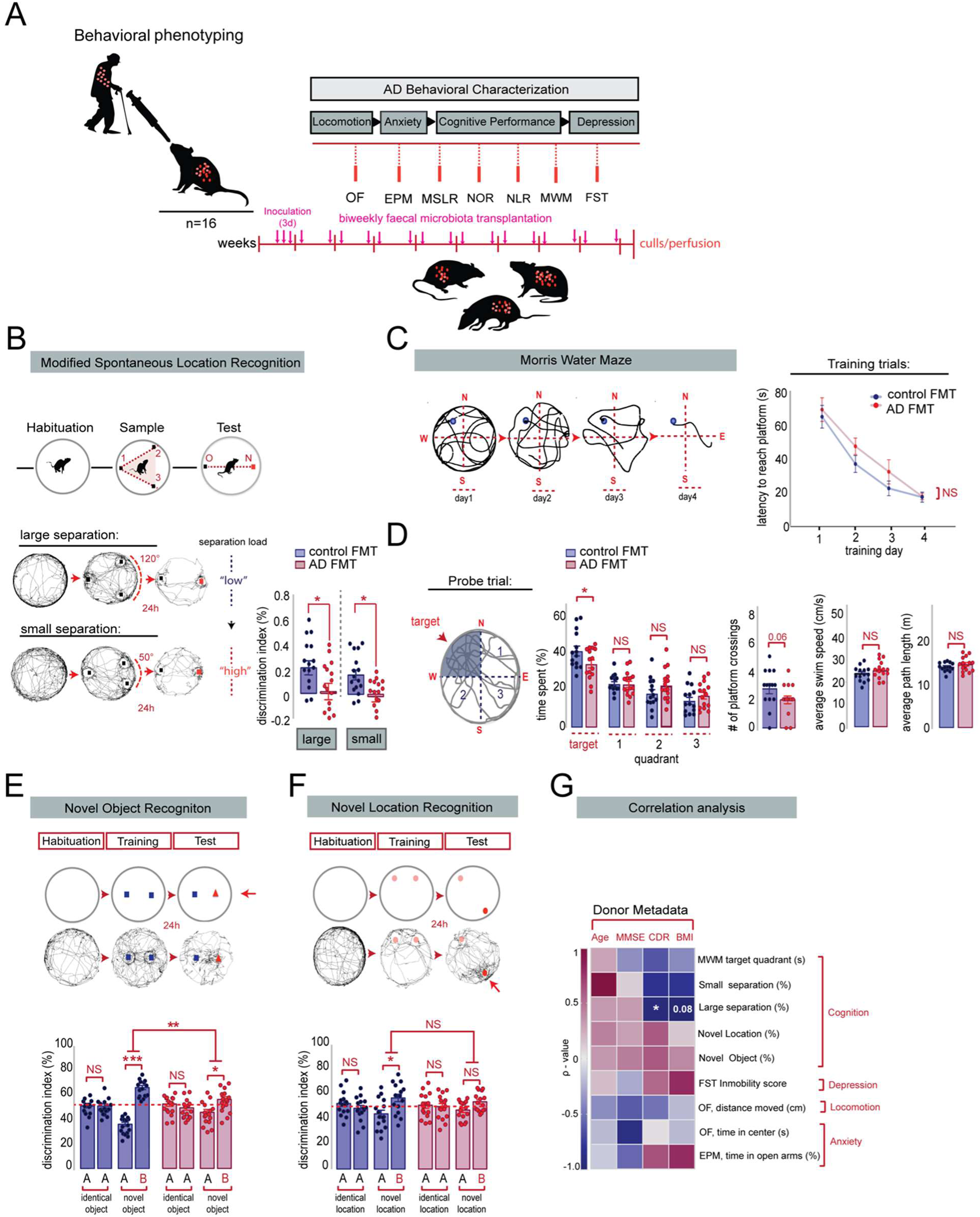
AD FMT induced cognitive deficits in hippocampal-neurogenesis dependent behaviours. **(A)** Overview of the AD-related behaviors. Vertical bars represent weekly intervals. Behaviors performed include: Open field (OF), Elevated Plus Maze (EPM), Modified Spontaneous Location Recognition test (MSLR), Novel Object Recognition (NOR), Novel Location Recognition (NLR), Morris Water Maze (MWM) and Forced Swim test (FST). **(B) Upper panel:** Schematic illustration of the MSLR task. **Lower panel:** AD FMT rats displayed a significant reduction in the discrimination index in the large separation task, two-tailed Student’s t-test, \**p*=0.0353 and small separation task, Mann-Whitney test, \**p*=0.0108. **(C) Left panel:** Representative tracing graphs in platform trials in the MWM **Right panel:** Escape latency during platform trials in MWM. No significant differences in learning was detected during the acquisition training days in the MWM test, Two-way mixed ANOVA, effect of FMT: *p*=0.2148, effect of time: \*\*\**p*<0.0001, FMT x time interaction: *p*=0.6672. **(D)** When challenged in the probe trial, AD FMT rats spent significantly less time in the target quadrant, unpaired, two-tailed Student’s t-test, \**p*=0.0498. Average number of platform crossings was reduced in AD FMT rats compared to control FMT rats, Mann-Whitney test, *p*=0.0670. AD FMT colonized rats displayed no significant change in the locomotory parameters of the probe trial, as determined by average swim speed, unpaired, two-tailed Student’s t-test, *p*=0.1582 and path length, unpaired, two-tailed Student’s t-test, *p*=0.2246. **(E) Upper panel:** Schematic illustration of the novel object recognition test. **Lower panel:** In the training session, no significant difference was detected between the differentiation index of the two identical objects in the control FMT rats, two tailed paired Student’s t-test, *p*=0.9859 and AD FMT rats, *p*=0.4149. In the test session, AD FMT rats show a significant reduction in the discrimination index of the novel objects compared to control FMT rats, unpaired, two-tailed Student’s t-test, \*\**p*=0.0019. **(F) Upper panel:** Schematic illustration of the novel object location test. **Lower panel:** In the training sessions, no significant difference was detected between the differentiation index of the two identical locations in the control FMT rats, two tailed paired Student’s t-test, *p*=0.4244 and AD FMT rats, *p*=0.8267. In the test session, AD FMT rats show no significant reduction in the discrimination index of the novel location compared to control FMT rats, unpaired, two-tailed Student’s t-test, *p*=0.4032. **(G)** Spearman’s rank and Pearson correlation between rat behavior and human donor metadata. Heatmap showing Spearman p (rho) or Pearsons R correlation coefficients, with red indicating strong positive correlation, and blue indicating strong negative correlation. P values for significant correlations (α<0.05) are noted. All data are presented as mean ± SEM, \**P*<0.05, \*\**P*<0.01, \*\*\**P*<0.001, NS, not significant.

Given previous reports in both human AD participants and transgenic animal models of AD that impairments of AHN contribute to memory deficits (Moreno-Jimenez et al., 2019), we employed the Modified Spontaneous Location Recognition test (MSLR) task to assesses pattern separation (Fig. 3B), which has been shown to be reliant on functional AHN (Bekinschtein et al., 2013). We found that rats harboring human AD microbiota were significantly impaired in discriminating between the familiar and novel locations in the “large separation condition” (low load on pattern separation) and “small separation condition” (high load on pattern separation) (Fig. 3B). Memory performance was consistently affected in a number of other cognitive tasks (Fig. 3C-F)., which have also shown to be dependent on AHN (Drapeau et al., 2003; Goodman et al., 2010; Jessberger et al., 2009). Whereas spatial learning in the Morris Water Maze (MWM) remained unaffected (Fig. 3C), humanized AD colonised rats spent significantly less time in the target quadrant during the probe trial than control FMT treated rats, indicating impairments in long-term spatial memory (Fig. 3D). Consistent with AD-relevant cognitive deficits, we found that rats colonised with material from AD donors performed significantly worse in a recognition memory task (Fig. 3E) and failed to discriminate the novel location in the novel location recognition test (Fig. 3F).

These observations were further corroborated by correlating the clinical human donor profile to the AD behavioral readouts of the recipient rats. We found an inverse correlation between the discrimination index of the MSLR (large separation) and the clinical dementia rating score (CDR) of the human donors (Fig. 3G), supporting the hypothesis that dementia donor-specific microbiota changes impact on cognitive function. Overall, our results demonstrate that the presence of human gut microbiota from AD participants promotes cognitive deficits related to the clinical symptoms of AD and impairs in particular AHN-dependent memory performance in recipient rats.

### Transplantation of gut microbiota from individuals with AD decreased adult hippocampal neurogenesis and impaired dendritogenesis of adult-born neurons

Based on these results and given the well-documented role of the dentate gyrus (DG) in supporting pattern separation and the potential susceptibility of this region to AD-related inflammatory changes, we assessed AHN and microglial activation in the rat DG after human FMT (Fig. 4A). Interestingly, rats harboring gut microbiota from AD participants did not show significant differences in microglia density (Extended Data Fig. 7A) and showed only minor differences in Iba1 somal size (Fig. 4b, Extended Data Fig. 7B) in the DG compared to control FMT colonised rats, suggesting that neuroinflammatory processes in this brain region may not mediate the cognitive impairments observed in AD FMT recipient rats.

**Figure 4:**
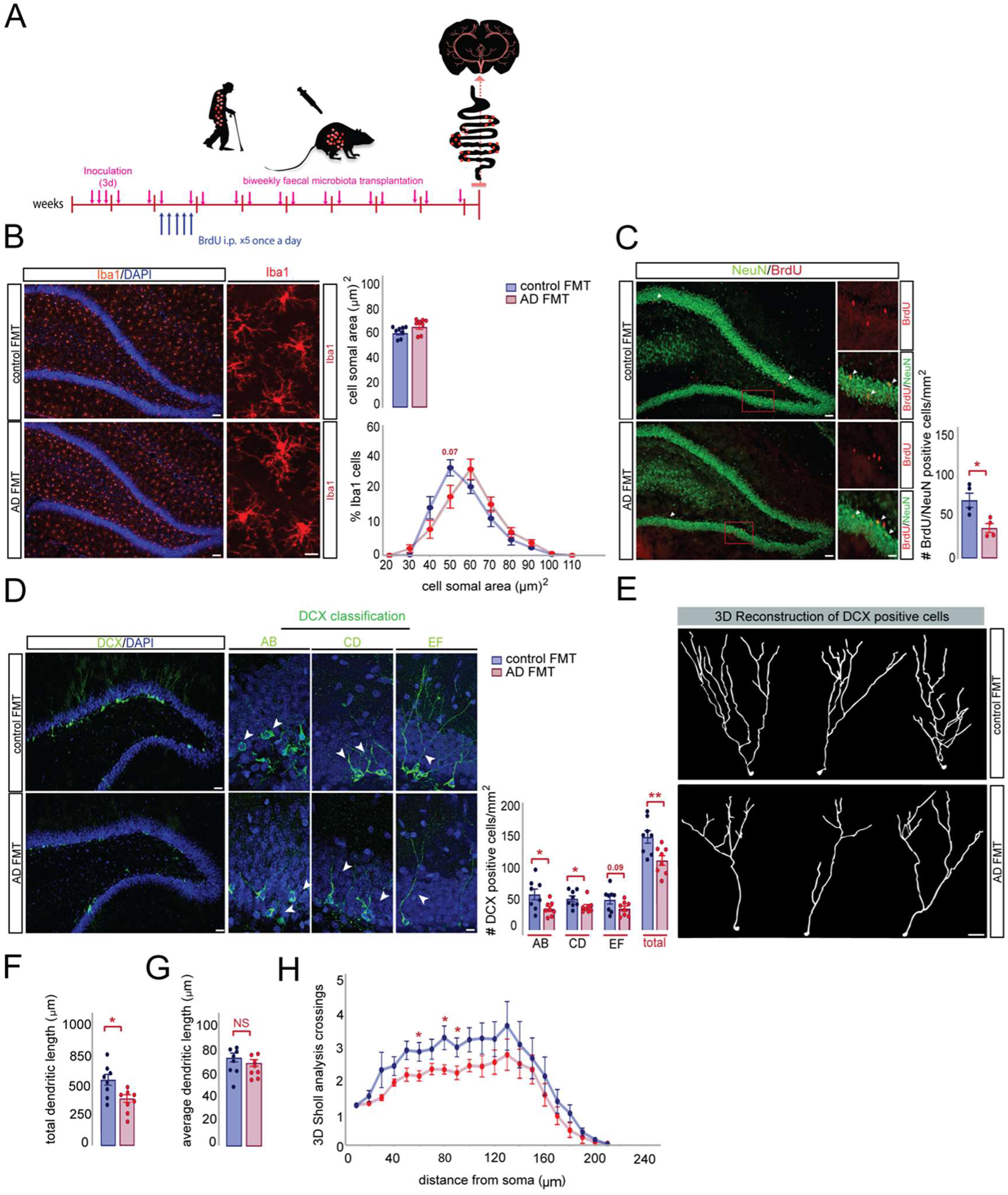
Transplantation of gut microbiota from AD participants decreased adult hippocampal neurogenesis and impaired dendritogenesis of adult-born neurons in rats. **(A)** Experimental timeline showing the effect of FMT on the survival of newborn neurons and microglia activation: Rats were injected with BrdU (150 mg/kg) once per day for 5 days, 10 days post donor colonization and sacrificed at day 59. **(B)** Left panel: Representative images of Iba1 positive cells in the DG of control and AD FMT rats. Scale bars: 200 µm left magnification, 40 µm right magnification. Right panel: Quantification of cell soma area of Iba1-positive cells in the DG of control and AD FMT rats, Mann-Whitney test, *p*=0.1605. Distribution analysis of soma area shows a shift from small to larger cell body sizes in AD FMT colonized rats in comparison to control FMT colonized rats. Two-way mixed ANOVA, effect of FMT: *p*=0.3154, effect of soma area: \*\*\**p*<0.0001, FMT x soma area interaction: *\*p*=0.0240. **(C)** Left panel: Representative images of BrdU/NeuN positive cells in DG of control and AD FMT rats. Scale bars: 200 µm left magnification, 50 µm right magnification. Newborn neurons were analyzed using double labeling for BrdU (red) and the neuronal marker NeuN (green). Arrowheads indicate double-positive cells (orange). Right panel: AD FMT rats show a significant reduction in the number of BrdU/NeuN cells, two-tailed Student’s t-test, \**p*=0.0159. **(D)** Left panel: Representative images of AB, CD and EF types of DCX-positive cells classified based on their dendritic tree morphology. Scale bars: 200 µm left magnification, 40 µm right magnification. Right panel: AD FMT rats show a significant reduction in the number of AB type, two-tailed Student’s t-test, \**p*=0.0290; CD type, Mann-Whitney test, \**p*=0.0281 and total number of neuroblasts in the DG, two-tailed Student’s t-test, \*\**p*=0.0099. **(E)** Representative 3D reconstructions of control and AD FMT DCX neurons. Scale bar represents 20 μm. **(F)** Morphological analysis revealed a significant decrease in the total dendritic length in AD FMT treated rats, two-tailed Student’s t-test, \**p*=0.0124 **(G)** The average dendritic length was unaltered between control and AD FMT rats, two-tailed Student’s t-test, *p*=0.3533. (H) Sholl analysis revealed a reduction in dendritic complexity in AD FMT rats compared to control FMT rats, Two-way mixed ANOVA, effect of FMT: \**p*=0.0452, effect of distance from soma: \*\*\**p*<0.0001, FMT x distance interaction: *p*=0.9358. All data are presented as mean ± SEM, \**p*<0.05, \*\**p*<0.01, NS, not significant.

To examine whether the impaired cognitive function in AD FMT colonised rats might be related to plaque formation, we performed Thioflavin-S fluorescent microscopy of coronal brain sections, which revealed the absence of plaque deposition in the hippocampus and cortex region in both treatment groups (Extended Data Fig. 7 C-D). We next measured the survival of new neurons in the DG by performing *in vivo* BrdU labeling and co-localization with the neuronal marker NeuN (Fig. 4C). A significant decrease in the number of BrdU/NeuN positive cells in the DG of rats colonised with AD FMT compared to control recipient rats was evident, indicating reduced survival of new neurons in the DG (Fig. 4C). In line with these findings, we observed that the number of DCX-positive neuroblasts was significantly lower in AD FMT colonised rats (Fig. 4D). This density reduction was specific to AB- and CD- but not EF-type neuroblasts (classified based on their dendritic tree morphology).

New neurons produced in the hippocampus of the DG have been shown to functionally integrate into mature circuits and contribute to the ability to recall memories. One requirement, however, for the successful synaptic integration and subsequent functionality of these neurons is the dendritic sprouting and proper arborisation into the molecular layer (ML) of the DG (Rosenzweig and Wojtowicz, 2011). To gain insight into the mechanism that may contribute to the observed memory deficits in AD colonised rats, we carried out 3D-reconstruction of DCX-positive cells using Sholl analysis to evaluate the complexity of their dendritic trees (Fig. 4E). Compared to control FMT colonised rats, DCX labeled cells in AD FMT treated rats displayed a significant reduction in the total dendritic length, whereas the average dendritic length was unaltered (Fig. 4F-G). Furthermore, when we quantified the number of dendritic intersections, we observed a significant reduction of dendritic complexity in AD colonised rats compared to control recipient rats (Fig. 4H).These data indicate that AD gut microbiota negatively affect the survival and dendritic arborization of adult-born neurons into the ML of the DG. Furthermore, our data reinforce previous findings in postmortem human brains that cognition and AHN is already altered before extracellular amyloid deposition (Moreno-Jimenez et al., 2019).

### Serum from individuals with AD decreased neurogenesis in human hippocampal progenitor cells *in vitro*

Since the hippocampal neurogenic niche is highly vascularised (Licht and Keshet, 2015; Pluvinage and Wyss-Coray, 2020; Smith et al., 2018) and gut microbiota composition explains up to 58% of the variance of individual plasma metabolites (Dekkers et al., 2022), AD gut microbiota-induced changes in the systemic environment, mediated by blood, have the potential to modulate AHN. Given the inaccessibility of the DG and the absence of validated biomarkers and neuroimaging tools to quantify hippocampal neurogenesis in living humans, we used our *in vitro* neurogenesis assay to study how the AD systemic environment modulates hippocampal neurogenesis (de Lucia et al., 2021; Du Preez et al., 2022; Villeda et al., 2014, 2011) (Fig. 5). To that end, human hippocampal progenitor cells (HPCs) were exposed to serum from control subjects and AD participants (Fig. 5A) and the percentage of cells expressing markers for neural stem cell proliferation (Ki67), differentiation (MAP2, DCX), and programmed cell death (CC3) was determined (de Lucia et al., 2020). Our results show that, while the average cell density of control and AD serum exposed HPCs cells remained unaffected (Fig. 5B), a reduction in the expression of Ki67 positive cells (Fig. 5B) in response to AD serum occurred, indicating that the serum of AD participants decreases the proliferative capacity of differentiating HPCs. Furthermore, following seven days of neuronal differentiation, we observed a decrease in the percentage of MAP2 and DCX positive neuroblasts in AD serum-treated cells (Fig. 5C), suggesting that the serum of AD subjects impaired neuronal differentiation.

**Figure 5:**
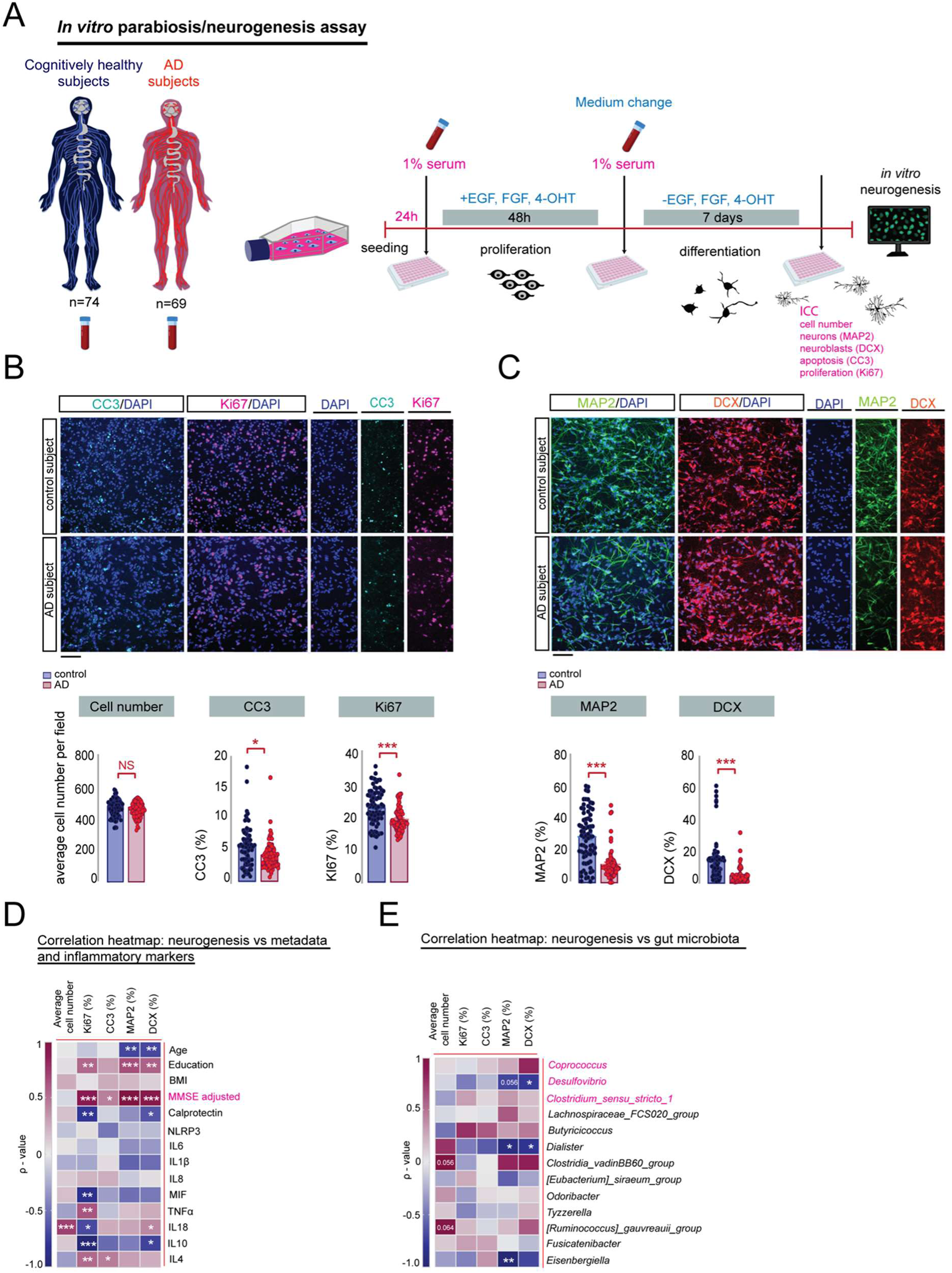
Serum from AD subjects decreased neurogenesis in human hippocampal progenitor cells. **(A)** Experimental timeline of the *in vitro* parabiosis/neurogenesis assay: human hippocampal progenitor cells were cultured with 1% human serum from age-matched cognitively healthy and AD participants and the following read-outs during the differentiation phase were obtained: cell number, apoptotic cell death (CC3), proliferating cells (Ki67), neurons (MAP2), neuroblasts (DCX). **(B) Upper panel:** Representative confocal micrographs showing CC3-positive HPCs cells (green: CC3-positive cells, blue: DAPI) and Ki67-positive HPCs cells (red: DCX-positive cells, blue: DAPI) during the differentiation phase of the assay. Scale bar: 100 μm. **Lower panel:** No significant change was detected in the average number of hippocampal progenitor cells per field between control and AD participants during the differentiation phase of the human serum assay, unpaired, two-tailed Student’s t-test, *p*=0.1855. Staining for the apoptotic marker CC3 revealed a significant decrease in cell death in HPCs cells receiving serum from AD participants compared to controls, Mann-Whitney test, \**p*=0.0100. Serum from AD participants induced a significant reduction in the expression level of the proliferation cell marker Ki67, Mann-Whitney test, \*\*\**p*<0.0001. **(C) Upper panel:** Immunofluorescence staining of MAP2-positive HPCs cells (green: MAP2-positive cells, blue: DAPI) and DCX-positive HPCs cells (red: DCX-positive cells, blue: DAPI) during the differentiation phase of the assay. Scale bar: 100 μm. **Lower panel:** Serum from AD participants caused a significant reduction in the expression levels of neurons (MAP2), Mann-Whitney test, \*\*\**p*<0.0001 and neuroblasts (DCX), Mann-Whitney test, \*\*\**p*<0.0001. Cellular readouts are expressed as percentage relative to neural cell number. 15 fields were analysed per well, n=3 technical replicates. **(D)** Spearman correlation between *in vitro* cellular neurogenesis readouts and human donor gut microbiota. **(E)** Spearman’s rank correlation between *in vitro* cellular neurogenesis readouts, human donor metadata and serum inflammatory markers. Correlation heatmaps showing Spearman’s rank coefficients, with red indicating strong positive correlation, and blue indicating strong negative correlation. *P* values for significant correlations (α<0.05) are noted. Control serum n=74, AD serum n=69. All data are presented as mean ± SEM, \**p*<0.05, \*\**p*<0.01, \*\*\**p*<0.001, NS, not significant.

To further determine whether the circulatory environment of AD participants can alter the morphological maturation of neuronal cells, we carried out a detailed morphometric analysis of MAP2 and DCX positive HPCs (Extended Data Fig. 8). Similar to previous reports using an AD transgenic mouse model (Ferreiro et al., 2020), we found that neuronal HPCs cells treated with AD serum displayed a substantially different morphology, characterised by an increase in the total neurite length and highly increased branching points in both MAP2 and DCX-positive cells (Extended Data Fig. 8B, C) compared to HPCs cells exposed to control serum. Overall, these data highlight that the systemic, circulatory environment of AD participants is capable of affecting neuronal proliferation, differentiation, and dendritic morphology.

Notably, we could again link these observations to the clinical phenotype of AD participants. We found a significant positive association between the MMSE score (Fig. 5E) and the prevalence of DCX, KI67, and MAP2 cells. Additionally, in line with our previous findings (Fig. 1E), the abundance of the pro-inflammatory genera *Desulfovibrio* and *Dialiste*r inversely correlated with the *in vitro* expression of the neuronal marker DCX (Fig. 5D). The association of specific bacterial genera with markers for neurogenesis, which are also highly correlated with the MMSE score (Fig. 1E), further support the hypothesis that impaired neurogenesis may be the mechanistic link between the observed altered gut microbiota composition and cognitive impairment in AD.

### Recipient rats transplanted with human AD gut microbiota show discrete metabolomic profiles

To better understand which factors may trigger the previously observed effects of AD FMT on AHN and associated behaviours, we performed untargeted metabolomic analyses of caecal content and hippocampal tissue from control and AD FMT colonised rats. Partial least squares discriminant analyses suggested an effect of donor status on the caecal (Figure 6A) and hippocampal (Figure 6B) metabolome. Indeed, differential expression analyses revealed that 13 (out of 184) metabolites in the caecal content (Figure 6C and 6E) and 3 (out of 123) metabolites in the hippocampus (Figure 6D and 6F) were nominally (*p* < 0.05) differentially abundant in AD FMT-compared to control FMT-colonised rats (for a full list of quantified features, see Supplementary Table 4). It should be noted however that metabolic features in either dataset did not pass the 5% FDR-based correction for multiple testing.

**Figure 6:**
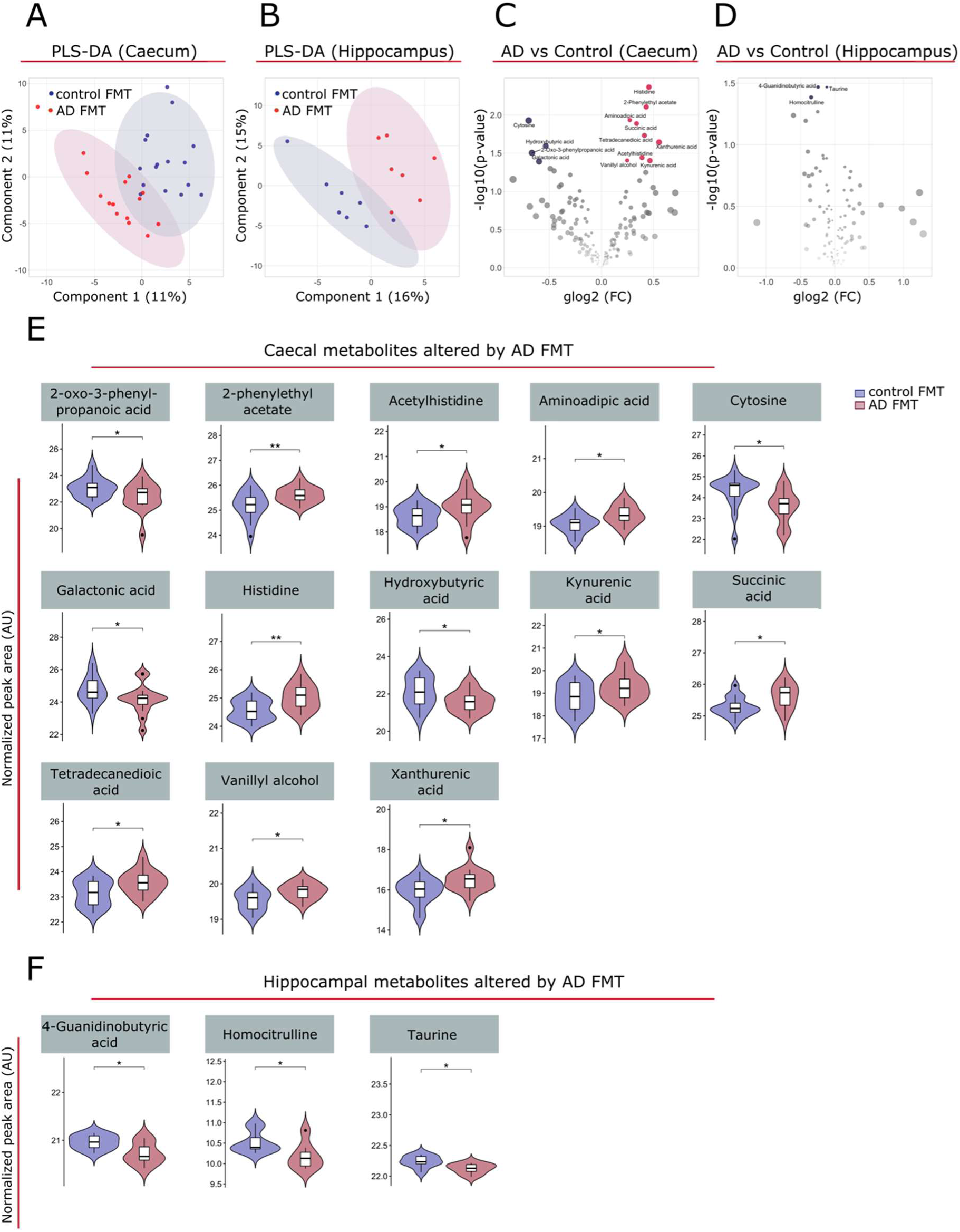
FMT from AD participants induced alterations in the caecal and hippocampal metabolomes. **(A-B)** Partial least squares discriminant analysis (PLS-DA) plot depicting the effect of control and AD FMT on the caecal content (**A**) and hippocampal (**B**) metabolomes. (with 95% concentration ellipses). Both models were non-significant (pQ^2^ > 0.05) by n=1,000 permutation tests. **(C-D)** Volcano plots of metabolites quantified in caecal content (n=184 features) (**C**) and hippocampal tissue (n=123 features) (**D**) of rats colonized with control and AD FMT. Red dots represent metabolites nominally (*p* < 0.05) upregulated in AD FMT rats, and blue dots represent metabolites nominally (*p* < 0.05) upregulated in control FMT rats. **(E-F)** Violin and box- and whisker plot (median represented by horizontal line) of normalized peak area of the caecal (n=15-16) and hippocampal (n=7-8) metabolites differentially regulated between rats receiving human FMT from control subjects and AD participants, as depicted in **C-D**. \**p*<0.05, \*\**p*<0.01 (unadjusted *p*-values from moderated *t*-tests, Limma). Histidine, *p* = 0.0043; 2-Phenylethyl acetate, *p* = 0.0079; Aminoadipic acid, *p* = 0.012; Cytosine, *p* = 0.012; Succinic acid, *p* = 0.013; Tetradecanedioic acid, *p* = 0.019; Xanthurenic acid, *p* = 0.023; Hydroxybutyric acid, p = 0.026; 2-Oxo-3-phenylpropanoic acid, *p* = 0.031; Acetylhistidine, *p* = 0.036; Vanillyl alcohol, *p* = 0.039; Kynurenic acid, *p* = 0.040; Galactonic acid, *p* = 0.041. 4-Guanidinobutyric acid, *p* = 0.034; Taurine, *p* = 0.034; Homocitrulline, *p* = 0.041. For a full list of quantified metabolites, refer to Supplementary Table 4. Abbreviations: FC, fold change; glog_2_, generalized logarithm base 2.

In caecal content, the amino acid histidine and its derivative acetylhistidine were nominally altered, in line with previous reports on peripheral metabolic profiles of both AD participants and transgenic AD mouse models (González-Domínguez et al., 2015; Hunsberger et al., 2020; Mazurkiewicz-Kwilecki and Nsonwah, 1989; Nielsen et al., 2021; Xi et al., 2021). Increased levels of histidine may indicate perturbed homeostasis of histamine (González-Domínguez et al., 2015), which is a critical regulator of the immune response and has been shown to influence AHN as well blood-brain barrier (BBB) integrity (Song et al., 2018). Additionally, histidine plays a critical role in the membrane binding and neurotoxicity of amyloid β fibrils (Smith et al., 2010) (Brännström et al., 2017). Aminoadipic acid was found to be nominally upregulated, and this has also been found in the plasma metabolome of people with AD and mild cognitive impairment compared to age- and sex-matched controls (Wang et al., 2014). Galactonic acid and succinic acid (nominally decreased and increased, respectively, in AD FMT rats) were found in one study to be increased in the urine metabolome of presenilin double knock-out mice, and this was correlated with gut microbiota composition changes (Gao et al., 2021). The tryptophan metabolites kynurenic acid and xanthurenic acid were nominally upregulated in AD recipient mice, which have been found to be altered in AD serum and urine and cerebrospinal fluid (Whiley et al., 2021). Increases in faecal concentration of the carboxylate tetradecanedioic acid, found nominally upregulated here, have been closely linked to aging and the prediction of neurocognitive disorders (Han et al., 2022; Xie et al., 2021). We also detected nominally decreased levels of the metabolite hydroxybutyrate. Beta-hydroxybutyrate mitigates against the pathogenesis of AD through various mechanisms, including through mitochondrial metabolism, metabolism of Aβ and tau proteins, inhibition of inflammation, promotion of AHN and through the regulation of cognitive function (Benjamin et al., 2017; Wang et al., 2022).

While many of the above metabolites can cross the BBB, they were not differentially abundant in the hippocampus. In the hippocampus, 4-guanidinobutyric acid and taurine were nominally downregulated. These metabolites were similarly found to be decreased in an age-dependent manner in the hippocampi of AD (3xTg) model mice (Zhao et al., 2021) and are known to cross the BBB (Tachikawa and Hosoya, 2011). Critically, taurine has been found to regulate adult (Gebara et al., 2015) and developmental (Hernández-Benítez et al., 2012) AHN, as evidenced both *in vivo* and *in vitro*. Homocitrulline was also found to be nominally downregulated. Interestingly, (homo)citrullination of AD-related and inflammatory proteins in the human medial temporal lobe proteome has been found to be increased in early AD compared to healthy age-matched controls, and these changes are associated with neuroinflammation (Gallart-Palau et al., 2017). Additionally, citrulline is increased in the plasma of people with AD (Corso et al., 2017) and the hippocampus of AD model (3xTg) mice (Zhao et al., 2017).

## DISCUSSION

To study the link between altered gut microbiota, inflammation, AHN and cognition, we transplanted faecal microbiota from AD and age-matched cognitively healthy subjects into microbiota-depleted naïve adult male rats. The results of the present study confirm that AD is characterised by peripheral and gut immune system activation. Gut inflammation, as determined by elevated levels of faecal calprotectin, has been reported in AD (Leblhuber et al., 2015), but studies also report no significant difference compared to cognitively healthy subjects (Kowalski and Mulak, 2022; Stadlbauer et al., 2020) reflecting the biological variability of calprotectin (D’Angelo et al., 2017) which can be affected by parameters such as medication use (Lundgren et al., 2019). Concerning peripheral immune activation, we found an increase in levels of pro-inflammatory cytokines IL-1β, NLRP3, MIF, an increase in IL-10 and a decrease in IL-4 at the transcription level in the blood of AD participants compared to controls. The levels of cytokines are possible indicators of neuroinflammation but many reports on their levels in AD are controversial or inconclusive (Brosseron et al., 2014). The reason for this variance arises from several factors, such as age range of the participants, proportion of each sex, the clinical phase, presence and extent of medical comorbidities, study design, sample preparation and methodological issues. In general, our data demonstrated an imbalance in the inflammatory profile of AD participants compared with controls and confirmed the involvement of NLRP3 (Heneka et al., 2013; Ising et al., 2019; Saresella et al., 2016) and MIF (Bacher et al., 2010; Craig-Schapiro et al., 2011; Lee et al., 2008; Popp et al., 2009) in AD pathology.

The decrease in the abundance of the phyla Firmicutes and the increase in Bacteroidetes observed in AD participants compared with age-matched cognitively healthy subjects was in line with several findings in the US (Vogt et al., 2017) and Chinese populations (Liu et al., 2019; Zhuang et al., 2018). At genera level and in line with our findings, reduced abundance of *Clostridium sensu stricto 1* is associated with adverse outcomes in AD (e.g., worse cognitive performance, neurodegeneration, and increased white matter lesion load) (Ling et al., 2020b; Vogt et al., 2017) and was more abundant in participants with better cognitive function and larger global brain volume (Karamujić-Čomić et al., 2020). *Desulfovibrio* has also been found to be enriched in other AD cohorts (Haran et al., 2019; Ling et al., 2020b) and is associated with reduced caecal levels of SCFAs and inflammation in mice (Sawin et al., 2015). As previously reported, we found a lower proportion of bacteria with the potential to synthetise butyrate (Haran et al., 2019; Ling et al., 2020b), a microbial metabolite negatively associated with cortical amyloid accumulation (Marizzoni et al., 2020a). Furthermore, we confirmed the reduced abundance of the anti-inflammatory genus *Coprococcus* (Ling et al., 2020b) in AD, which is associated with higher odds of amyloid positivity (Verhaar et al., 2021). However, when interpreting microbiota data, it is important to keep in mind that results may be influenced by geography (He et al., 2018), the inclusion criteria of study participants and methodological difference (i.e. sample collection, storage methods, different bioinformatics pipelines) (Choo et al., 2015; Marizzoni et al., 2020b). Not surprisingly, human feaces from different donors engrafted in rats at different rates. After transfer of faecal microbiota from cognitively healthy subjects to rats however, the taxa diversity remained relatively stable over time, whereas after transfer from AD donors, there was a greater alteration in taxa between 10- and 59-days post FMT. Notably, one of the genera that was increased at day 59 post AD-FMT compared to day 10 was *Desulfovibrio*, a genus that was also significantly enriched in AD participants.

To determine whether the changes in the gut microbiota composition in participants with AD precipitate symptoms associated with AD, we performed a series of behavioural tests on rats colonised with faecal material from cognitively healthy or AD subjects. Importantly, we found that an impairment in pattern separation, which is reported in AD (Ally et al., 2013; Parizkova et al., 2020) and is associated with indicators of Alzheimer’s pathology such as increased cortical beta-amyloid burden (Webb et al., 2020), medial temporal lobe tau (Leal et al., 2019), cerebrospinal fluid levels of phosphorylated tau (Berron et al., 2019), and apolipoprotein E genotype (Sinha et al., 2018), was transferred via the gut microbiota to manifest as a behavioural impairment in rats. While alterations in the gut microbiome have been associated with hippocampal-dependent cognition during ageing (Connell et al., 2022), there is no published evidence to date of the contribution of gut microbiota to this AHN-reliant behaviour. We found that AD-FMT also induced an impairment in long-term spatial memory, which is also associated with AHN levels (Drapeau et al., 2003), and supports a recent intra-species report using a transfer of faecal sample from the 5xFAD mouse model of AD into a WT recipient (Kim et al., 2021). We also observed that recognition memory was impaired in AD-FMT rats which is in line with previous work showing similar impairment in mice that received FMT from AD participants (Fujii et al., 2019).

Interestingly, the memory impairments observed in rats after FMT from AD donors are reliant on AHN and are supported by a recent study suggesting that defective neurogenesis contributes to memory failure in AD (Mishra et al., 2022). Accumulating evidence suggests that the gut microbiome can impact AHN; germ-free mice display increased AHN (Ogbonnaya et al., 2015), and antibiotic treatment reduced AHN, which was reversed by probiotic treatment in mice, possibly through brain monocytes (Möhle et al., 2016) and voluntary exercise in rats (unpublished observation). While we did not find any significant change in the hippocampal microglial status in rats harbouring AD gut microbiota, we found that the survival of new neurons and the number of newly-born neurons was decreased, which supports our behavioural observations. We also found that AD-FMT rats displayed an impairment in the morphological development of these newborn neurons, rendering a delay or a defect in this critical process of enabling functional integration into hippocampal circuits (Zhao et al 2008, Deng et al, 2010). In line with these findings, intra-species FMT (5xFAD to WT mice) reduced the survival and proliferation of neurons (Kim et al., 2021), and the recruitment and dendritic complexity of immature neurons was deficient in a familial AD mouse model (Mishra et al., 2022). Interestingly, recent studies have shown that that transfer of a WT mouse microbiome into APPswe/PSEN1dE9 mice improved spatial memory and synaptic plasticity (Sun et al., 2019), that reconstitution with normal gut flora in newborn mice protected against decreased AHN and cognitive deficits in antibiotic-treated mice (Liu et al., 2022), and that augmentation of AHN restored the dendritic development of new neurons, coupled with an enhancement of spatial recognition memory (Mishra et al., 2022). Thus, future studies should address whether the changes we observed in adult rats that received FMT from AD human donors are reversible, as well as mechanistic pathways that connect the gut microbiome to AHN.

In line with our findings in rats *in vivo*, using the *in vitro* neurogenesis assay, we observed decreased neurogenesis in HPCs treated with blood serum from participants with AD; we found a lower level of Dcx-positive neuroblasts and Map2-positive immature neurons. These findings are in agreement with studies that report fewer immature neurons in with AD compared to cognitively healthy controls (Zhou et al, 2022; Moreno-Jiménez et al. 2019, Li et al, 2008). We also found a decrease in the percentage of proliferating HPCs after treatment with AD serum which replicates our previous findings in an independent cohort of AD and cognitively healthy subjects (Walgrave et al., 2021). Intriguingly, we previously found in cognitively healthy subjects that an increased baseline cell death during differentiation predicts cognitive decline over the subsequent 12 years (Du Preez et al., 2022), whereas, here, we observe a decrease in cell death after treatment with AD serum, suggesting that there is a differential effect of AD serum on cell death during the disease course. Neurite outgrowth is a critical step in the formation of new neurons and integration into the hippocampal circuitry (Rodríguez-Iglesias et al., 2019). When we examined how the AD systemic environment influences neurite outgrowth *in vitro*, complexity measures were increased after AD serum treatment. Our findings are in line with a study that showed enhanced neurite outgrowth and arborisation in a primary neuronal culture from transgenic Drosophila melanogaster larvae expressing human Aβ42 compared to control (Saad et al., 2015), which occurred prior to the AD pathology. Surprisingly, our findings of longer and more complex neurites under AD serum conditions *in vitro* contrasts with our observations in the rat DG. One of the reasons for the opposing finding in the AD-FMT adult rat HPCs and the AD-serum treated embryonic human HPCs *in vitro* could be the difference in maturation rates of HPCs. García-Corzo et al. (García-Corzo et al., 2022) demonstrated that adult HPCs exhibit various molecular and cellular properties that differ from those present in developmental HPCs including more restricted multipotency, acquisition of a quiescence state and different transcriptomic programs underlying those cellular processes. Thus, the differences in the neurite outgrowth we observe may be due to maturation differences between immortalised foetal human HPCs and adult rat HPCs.

When we investigated the association between in vitro neurogenic readouts and the gut microbiota composition of the serum donors, we found that the percentage of DCX-positive neuroblasts inversely correlated with pro-inflammatory genera Desulfovibrio and Dialister and the percentage of Map2-positive cells inversely correlated with genera Desulfovibrio, Dialister and Eisenbergiella. This is interesting considering a previously published observations that genus Eisenbergiella increases with severity of cognitive impairment (Stadlbauer et al., 2020) and that genus Desulfovibrio was negatively correlated with MMSE scores (Palmas et al., 2022; Ren et al., 2020).

The human gut microbiota produces a myriad of bioactive molecules, and its composition has been found to account for a significant amount of variation in individual circulating metabolite levels (Dekkers et al., 2022). We found alterations in the caecal and hippocampal metabolomes of rats colonised with AD FMT compared to control FMT. The majority occurred in the caecum, but these were not reflected in the hippocampus, suggesting they may not directly influence AHN. The limited amount of differential expression and low effect sizes may be due to the variable levels of engraftment (i.e. by donor and genera) and reversion of microbial signatures with time, and thus did not allow for pathway analyses. Notably, however, the free amino acid taurine was nominally decreased in AD FMT rats. Taurine administration has repeatedly been shown to increase hippocampal neural stem and progenitor cell proliferation, survival, and net neurogenesis both *in vivo* and *in vitro* (Gebara et al., 2015), likely by acting on the ERK pathway (Shivaraj et al., 2012; Wu et al., 2022). Given its BBB permeability (Tachikawa and Hosoya, 2011), it is conceivable that some of the observed effects of AD FMT on AHN could partly be mediated via taurine. The current data suggest that microbial metabolites could impact both cognitive function and AD pathology, possibly linking specific microbial signatures to the observed changes in hippocampal neurogenesis and associated cognitive behaviours.

In conclusion, our results demonstrate that colonisation of rats with the faecal samples from individuals with AD, but not from cognitively healthy controls induced behavioral and neurogenic alterations in rats typical of AD. We further show that the expression of caecal metabolites involved in the neurogenic process and cognitive function are altered after faecal transplant from individuals with AD. As well as the gut microbiota and microbial metabolites having a causal role in the behavioural and AHN alterations in AD, we found a direct and negative impact of serum from individuals with AD on neurogenesis *in vitro*. Future studies could help elucidate the link between the gut, circulatory system and brain in the manifestation of cognitive changes in AD by aligning metabolomic analyses of caecal content, blood, and hippocampal tissue. In addition, future studies should include participants earlier in the disease course (e.g. individuals with mild cognitive impairment) to examine at which stage the gut microbiome diverges and how this impacts on AHN and cognition. Overall, our data reveal that impaired neurogenesis may be a mechanistic link between altered gut microbiota and cognitive impairment in AD.

## Acknowledgements

We thank Tara Foley, Gerry Moloney, Patrick Fitzgerald Suzanne Crotty and Anna Golubeva for technical assistance. This project was funded by the Network of Centres of Excellence in Neurodegeneration (COEN) via by Science Foundation Ireland for SG, SN, SD-H, JE, AL, CO’N and YN (SFI 17/COEN/3475 & SFI 19/FFP/6820), and the Italian Ministry of Health for MM, NL, EM, CS, DVM, MR and AC, and the Medical Research Council (MRC) UK for ES, KH and ST (MR/S00484X/1). YMN, AL and CO’N are funded Investigators of the APC Microbiome Ireland, which is a research centre funded by SFI, through the Irish Government’s National Development Plan (Grant number 12/RC/2273).

## Author information

These authors contributed equally: Stefanie Grabrucker, Moira Marizzoni, Edina Silajdžić

These authors contributed equally: Sandrine Thuret, Annamaria Cattaneo, Yvonne M Nolan

## Contributions

S.G., M.M., E.S., S.T., A.M.C., C.O’N and Y.N. designed and planned the study. S.G., E.S., S.N., N.L., E.M. were involved in data collection. S.G., M.M., E.S., S.N., S.D-H., K.H. performed data analysis. S.G., M.M., E.S., S.N., S.D-H., A.L. J.E. C.O’N, S.T., A.M.C. and Y.N. were involved in data interpretation. S.G., M.M., E.S., and Y.N. drafted the article with critical revision from all authors.

## Ethics declarations

### Competing interests

SG, SN, SD-H, AL, JE, CO’N, YN, ES, KH, MM, NL, EM, CS, DVM, MR, AC and ST declare no conflict of interest in relation to this study.

---

**Extended Data Figure 1:**
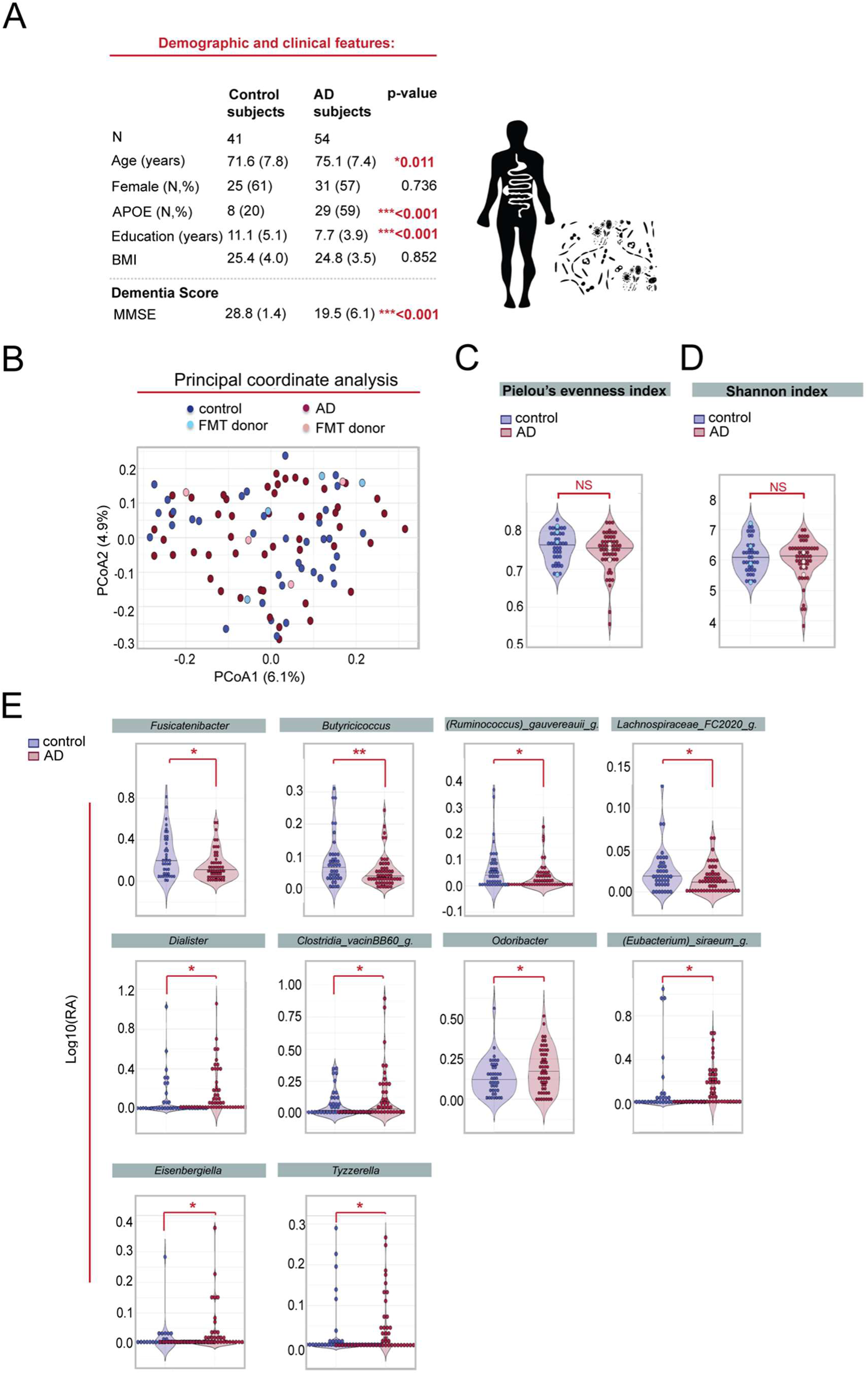
Taxonomic characterization of gut microbiota composition in AD participants and cognitively healthy control subjects. **(A)** Demographic and clinical features of AD participants (n=54) and cognitively healthy control subjects (n=41) used for stool sample analysis. *P*-values were calculated using unpaired, two-tailed Student’s t-test for continuous Gaussian variables (or Mann-Whitney test for non-Gaussian variables) and Chi-square test for categorical data. There were 6 missing values for APOE, MMSE and BMI. AD participants were older (\**p*=0.011), showed a significant increase in the prevalence of APOE carriers (\*\**p*<0.001), were less educated (\*\**p*<0.001) and had lower MMSE score (\*\**p*<0.001) than cognitively healthy controls. **(B)** Principal coordinate analysis (PCoA) of Bray Curtis distances from 16S rRNA sequencing of AD participants and cognitively healthy controls. The eight samples used for FMT inoculation experiments are highlighted in light blue for controls and light pink for AD subjects. No significant differences were reported between groups (PERMANOVA adjusted for age and batch effect, *p*=0.137). **(C-D)** Alpha diversity as measured by Pielou’s evenness and Shannon indexes from 16S rRNA sequencing of AD participants and cognitively healthy controls. Light blue and light pink symbols denote samples used for FMT inoculation. No significant differences were reported between groups as tested by one-way ANOVA adjusted for age and batch effect (Pielou’s evenness index, *p*=0.226; Shannon index, *p*=0.373). **(E)** Violin plots representing the relative abundance of 10 genera that were significantly altered between controls and AD participants using Mann-Whitney test, without FDR correction. (*Fusicantenibacter*, *p*=0.014; *Butyricicoccus*, *p*=0.006; [Ruminococcus]_gauvereauii_group, *p*=0.028; Lachnospiraceae_FC2020_group, *p*=0.015; Dialister, *p*=0.019; Clostridia_vacinBB60_group, *p*=0.046, Odoribacter, *p*=0.041; [Eubacterium]_siraeum_group, *p*=0.033, Eisenbergiella, *p*=0.050; Tyzzerella, *p*=0.040). All data are presented as mean ± SEM, \**p*<0.05, \*\**p*<0.01, NS, not significant.

**Extended Data Figure 2:**
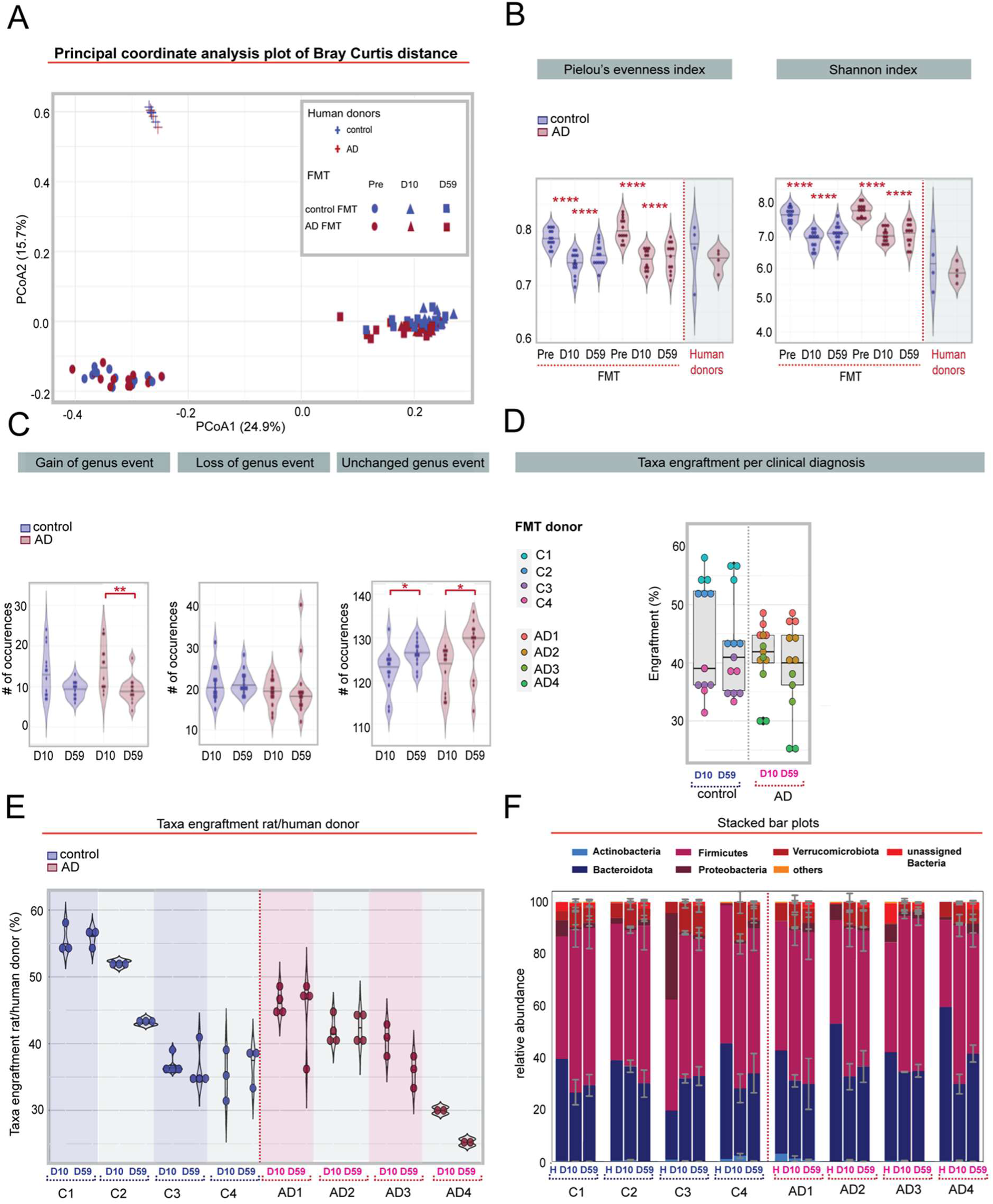
16S rRNA analysis of the faecal microbiome in human inoculated rats. **(A)** Principal coordinate analysis of the compositions of the faecal microbiome as measured by Bray-curtis distance of rats and human donors. **(B)** Violin plots showing alpha diversity as measured by Pielou’s evenness and Shannon indexes of rats and human donors. Black horizontal lines in violin plots represent the medians. **(C)** Violin plots showing the number of genera gained, lost and unchanged per rat 10 and 59 days after FMT. The fraction of genera in rats before antibiotic treatment and at D10 or D59 is shown. The abundance of genera gained decreased from D10 to D59 in rats that received AD FMT (\*\**p*=0.003), and the number of unchanged genera increased for both FMT treatment (control FMT group, \**p*=0.050; AD FMT group, \**p*=0.023). **(D)** Taxa engraftment in rats at the genera level by diagnosis. **(E)** Taxa engraftment in rats at the genera level by donor. **(D-E)** The percentage of genera in rats **(D)** and respective human donors is shown **(E)**. No significant difference was reported. **(F)** Stacked bar plots showing the taxonomic profile in donors and rats after 10 days (D10) and 59 days (D59) from the first FMT at the phylum level by diagnosis **(D)** and by donor **(E)**. Difference in means in **(C-E)** were tested by two-way ANOVA with repeated measures with donor diagnosis as between-subject factor and time from the first FMT as within-subject factor. All data are presented as mean ± SEM, \**p*<0.05, \*\**p*<0.01, \*\*\**p*<0.001, \*\*\*\**p* < 0.0001, NS, not significant.

**Extended Data Figure 3:**
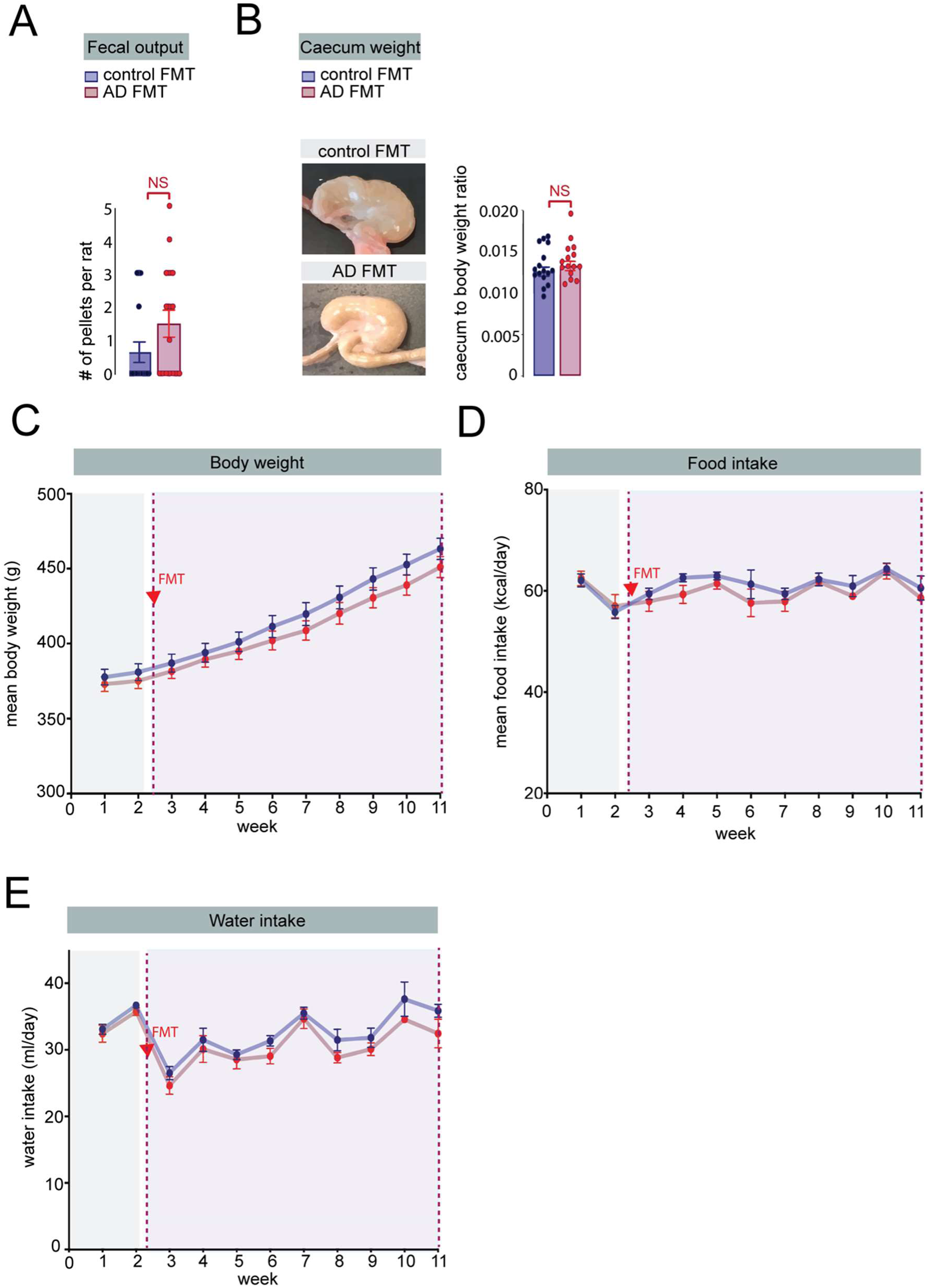
Effect of FMT transplantation on physiological parameters. **(A)** Faecal pellets produced in a 15 min period by control and AD FMT rats. AD FMT had no effect on faecal pellet output, Mann-Whitney test, *p*=0.116. **(B)** Caecum weight of control and AD FMT rats. Caecum weight did not differ in AD FMT rats compared to controls, unpaired, two-tailed Student’s t-test, *p*=0.3731. **(C)** Body weight composition of control and AD FMT rats over time. Mean body weight did not differ in rats colonized with AD FMT compared to rats colonized with control FMT, Two-way mixed ANOVA, effect of FMT: *p*=0.3356, effect of time: \*\*\**p*<0.001, FMT x time interaction: *p*=1.605, control FMT n=16, AD FMT n=16. **(D)** Food intake of control and AD FMT rats over time. AD FMT had no effect on food intake (per donor cage), Two-way mixed ANOVA, effect of FMT: *p*=0.0992, effect of time: \**p*=0.0380, FMT x time interaction: *p*=0.9447, control FMT n=4, AD FMT n=4. **(E)** Water intake measured over the course of the study from control and AD FMT rats. AD FMT had a significant effect on water intake (per donor cage), Two-way mixed ANOVA, effect of FMT: \**p*=0.02, effect of time: \*\*\**p*<0.001, FMT x time interaction: *p*=0.9855, control FMT n=4 cages, AD FMT n=4 cages. All data are presented as mean ± SEM, NS, not significant.

**Extended Data Figure 4:**
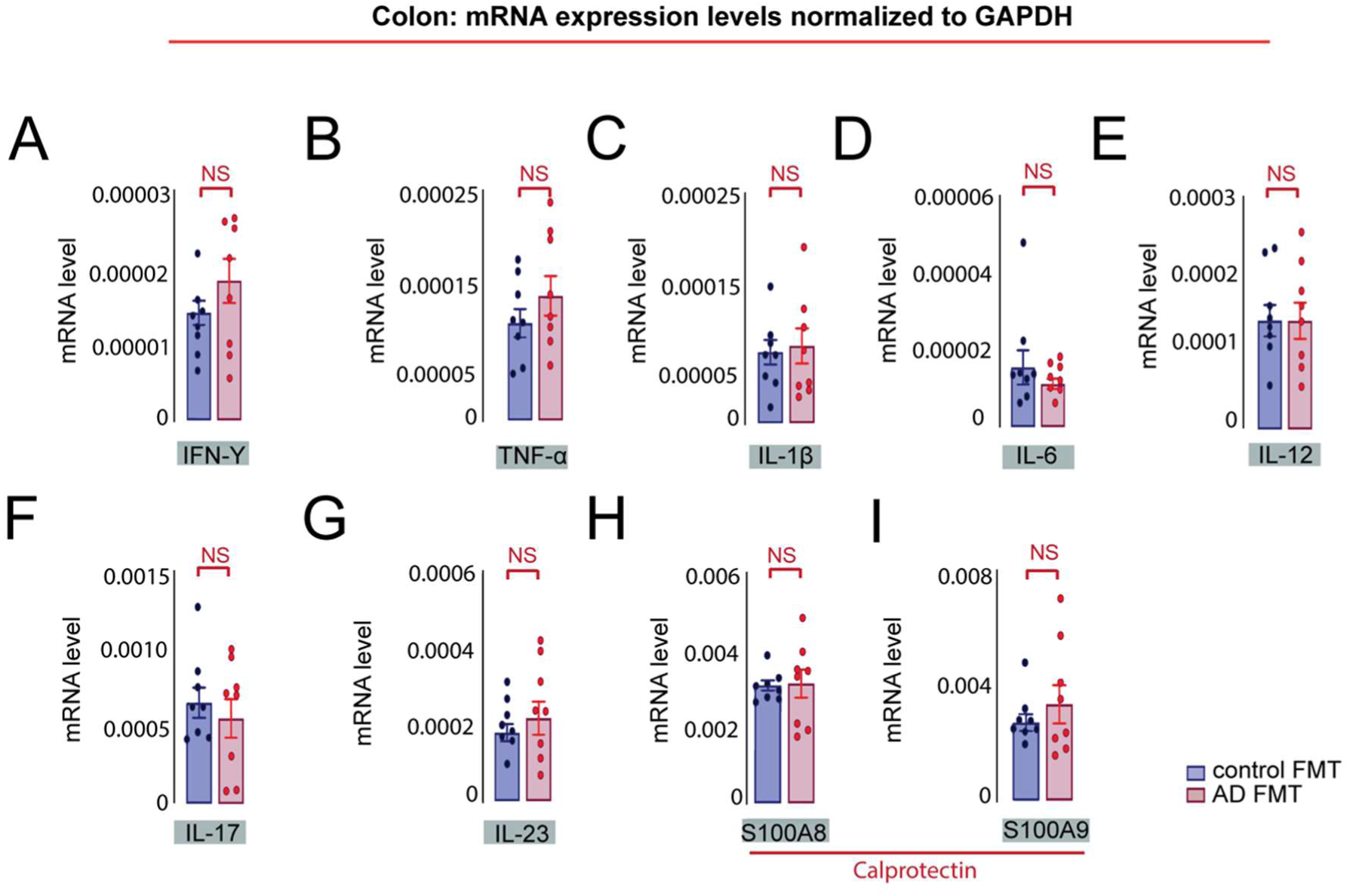
FMT from AD participants had no effect on pro-inflammatory cytokine expression in the rat colon. Relative cytokine mRNA expression in the proximal colon of rats colonized with control and AD FMT. No significant difference was detected in the mRNA expression of **(A)** IFN-α, unpaired, two-tailed Student’s t-test, *p*=0.2248; **(B)** TNF-α, unpaired, two-tailed Student’s t-test, *p*=0.2794; **(C)** IL-1β, unpaired, two-tailed Student’s t-test, *p*=0.775; **(D)** IL-6, Mann-Whitney test, *p*=0.6456; **(E)** IL-12, unpaired, two-tailed Student’s t-test, *p*=0.9947; **(F)** IL-17, unpaired, two-tailed Student’s t-test, *p*=0.5321; **(G)** IL-23, unpaired, two-tailed Student’s t-test, *p*=0.4496; **(H)** S100A8, unpaired, two-tailed Student’s t-test, *p*=0.9039; **(I)** S100A9, Mann-Whitney test, *p*=0.999, control FMT n=8, AD FMT n=8. All data are presented as mean ± SEM, NS, not significant.

**Extended Data Figure 5:**
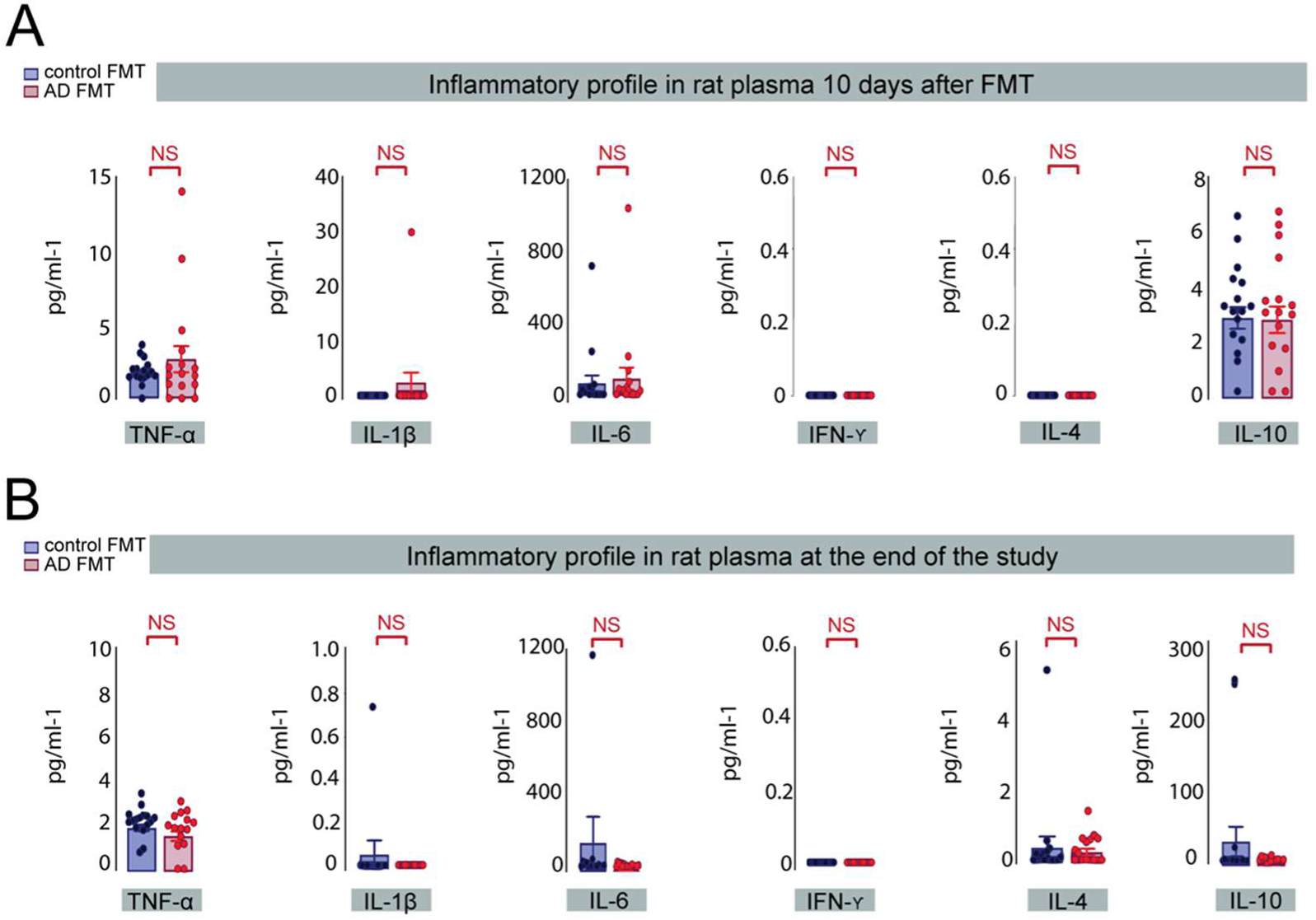
Peripheral inflammation is similar in rats colonized with AD microbiota and control microbiota. Pro-inflammatory cytokine protein expression in the plasma of rats colonised with control and AD FMT as measured by Meso Scale V-PLEX assay. **(A)** AD FMT rats display no significant change in TNF-α, Mann-Whitney test, *p*=0.9625; IL-1β, Mann-Whitney test, *p*=0.9999; IL-6, Mann-Whitney test, *p*=0.7141 IFN-α, Mann-Whitney test, *p*=0.9999; IL-4, Mann-Whitney test, *p*=0.9999 and IL-10, two-tailed Student’s t-test, p=0.9142 72 h after initial FMT inoculation, control FMT n= 16, AD FMT n=16. **(B)** AD FMT rats displayed no significant change in TNF-α, two-tailed Student’s t-test, p=0.1605; IL-1β, Mann-Whitney test, *p*=0.9999; IL-6, Mann-Whitney test, *p*=0.5199; IFN-α, Mann-Whitney test, *p*=0.9999; IL-4, Mann-Whitney test, *p*=0.5836 and IL-10, Mann-Whitney test, *p*=0.7701 at the end of the study, control FMT n=16, AD FMT n=15. All data are presented as mean ± SEM, NS, not significant.

**Extended Data Figure 6:**
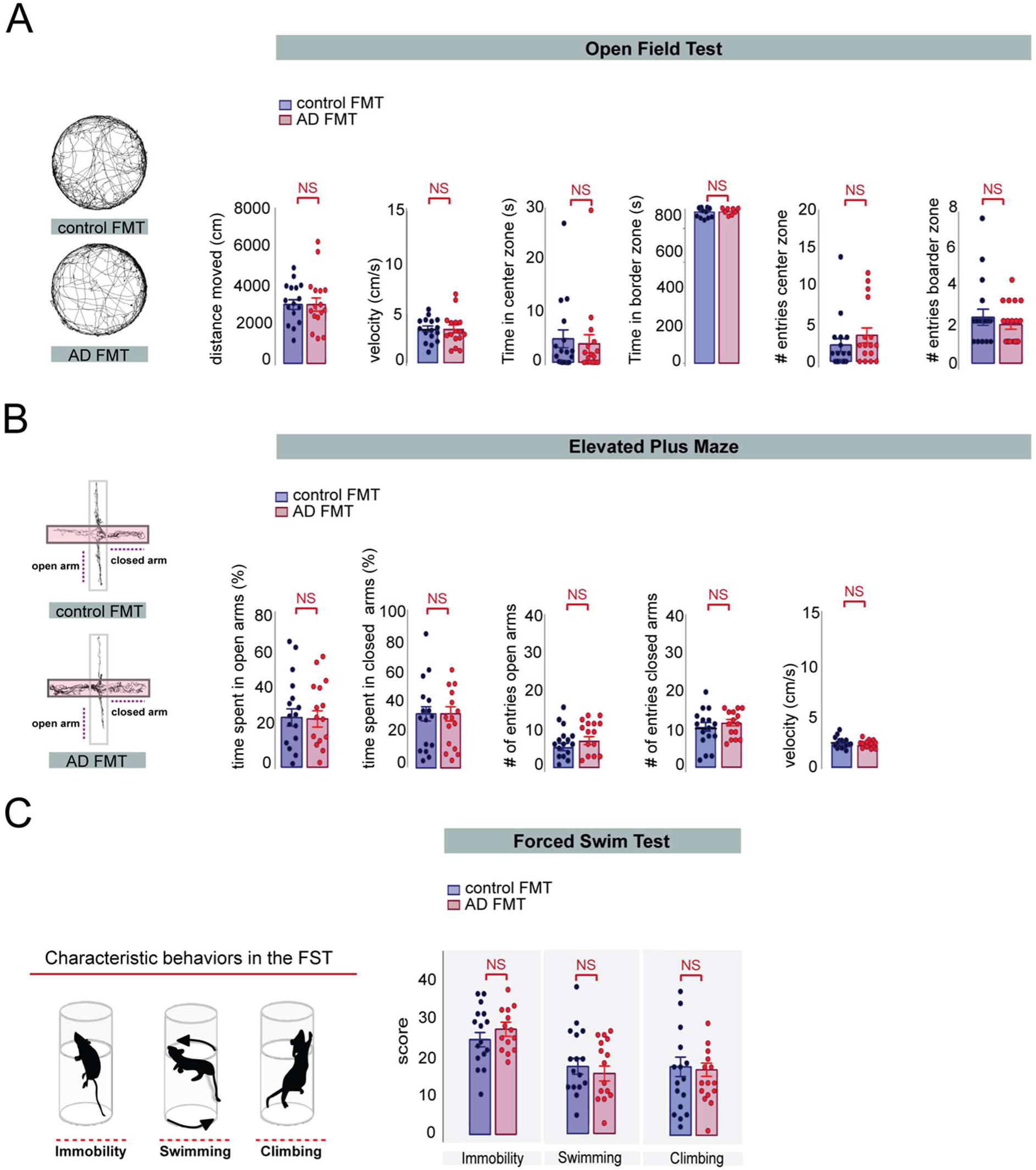
Colonisation with faecal material from AD participants had no impact on locomotion, anxiety and depressive like behavior in rats. **(A) Left panel:** Examples of trajectories of rats colonised with human control and AD FMT in the open field test. **Right panels:** AD FMT rats travelled a similar distance during a 30 min free exploration time in the circular open field arena, unpaired, two-tailed Student’s t-test, *p*=0.995. AD FMT colonised rats displayed no significant change in velocity in the open field test, unpaired, two-tailed Student’s t-test, *p*=0.9972. No significant change was detected in time spent in the center zone, Mann-Whitney test, *p*=0.4547 or border zone, Mann-Whitney test, *p*=0.5490. Additionally, AD colonised rats displayed no significant difference in the number of entries in the center zone, Mann-Whitney test, *p*=0.3056, or numbers of entries into the border zone, Mann-Whitney test, *p*=0.8062 compared to control FMT rats, control FMT n=16, AD FMT n=16. **(B) Left panel:** Example of tracking paths of rats colonised with human control and AD FMT in the Elevated Plus Maze test. **Right panels:** No significant change was detected in percent time spent in the open arms, unpaired, two-tailed Student’s t-test, *p*=0.9004 or closed arms, unpaired, two-tailed Student’s t-test, *p*=0.7994. No significant change was detected in the number of entries in the open arms, unpaired, two-tailed Student’s t-test, *p*=0.2474 or closed arms, unpaired, two-tailed Student’s t-test, *p*=0.4487. Additionally, AD FMT colonised rats displayed no significant change in velocity in the Elevated Plus Maze, unpaired, two-tailed Student’s t-test, *p*=0.555. **(C) Left panel:** Graphical illustration of the modified rat Forced Swim Test. **Right panel:** AD FMT had no effect on the mean score of immobility, unpaired, two-tailed Student’s t-test, *p*=0.3179; swimming, unpaired, two-tailed Student’s t-test, *p*=0.5200 and climbing behavior, two-tailed Student’s t-test, *p*=0.8015. All data are presented as mean ± SEM, NS, not significant.

**Extended Data Figure 7:**
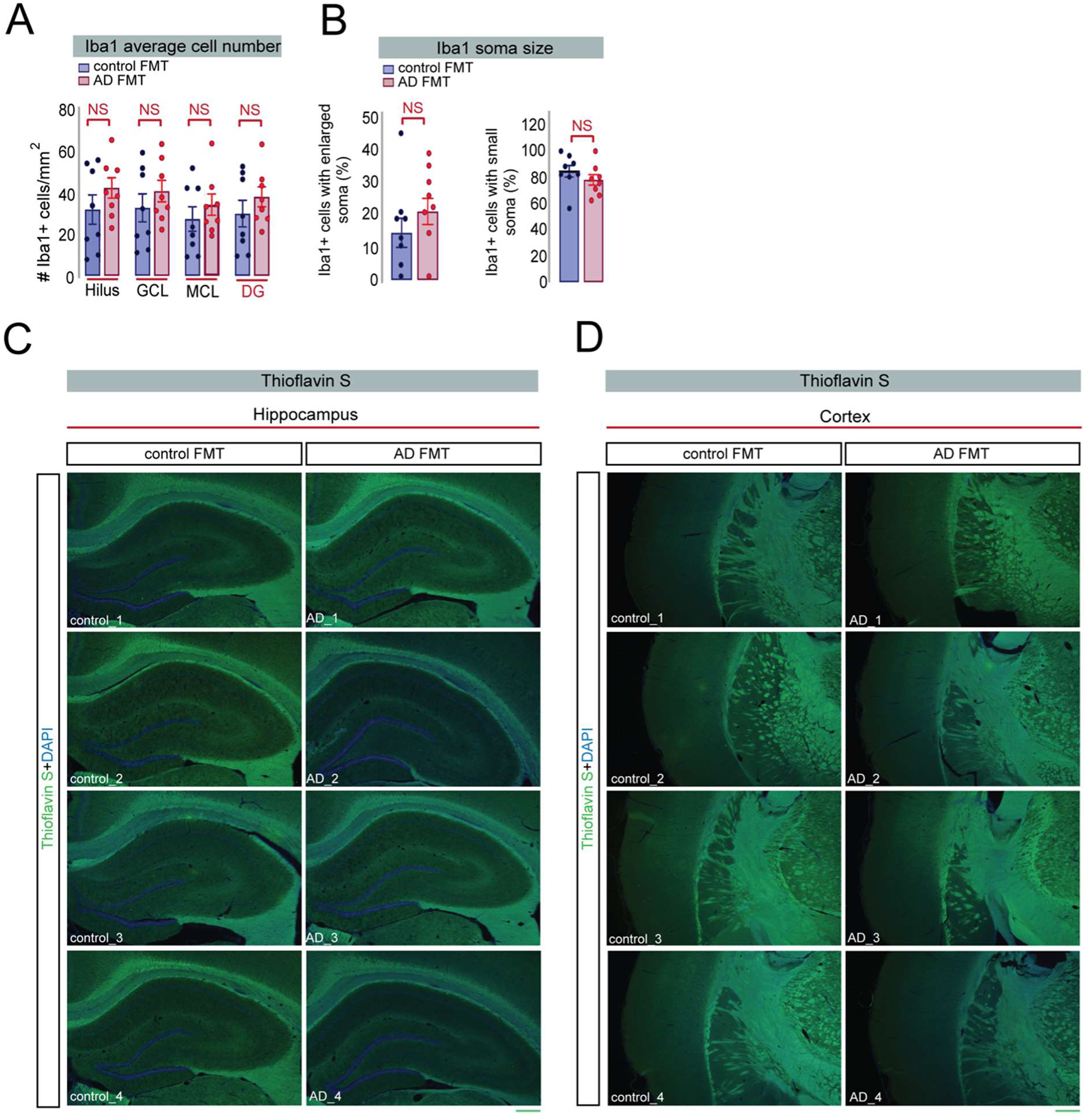
Colonization of faecal material from AD participants had no impact on Iba1 cell density and on amyloid plaque formation in the hippocampus and cortex of rats. **(A)** Quantification of microglia density in the dentate gyrus (DG) of human control FMT and AD colonised rats. No significant difference was detected in the average cell density of Iba1 positive cells in the Hilus, two-tailed Student’s t-test, *p*=0.2472; GCL, two-tailed Student’s t-test, *p*=0.3552; MCL, two-tailed Student’s t-test, *p*=0.2472 and whole DG, two-tailed Student’s t-test, *p*=0.3267 in AD colonised rats compared to control FMT inoculated rats. **(B)** No significant difference was detected in the percentage of Iba1 positive cells with enlarged soma area, two-tailed Student’s t-test, *p*=0.2932 and percentage of Iba1 cells with small soma area, two-tailed Student’s t-test, *p*=0.2932 **(C-D)** Representative Thioflavin-S staining (green: Thioflavin-S, blue: DAPI) of coronal brain sections from the hippocampus and cortex, (control and AD donor n=4). Absence of amyloid plaque formation in the hippocampus **(C)** and cortex **(D)** of control and AD FMT rats, Scale bar 500 µm. All data are presented as mean ± SEM, NS, not significant. Abbreviations: (GCL) Granule cell layer, (MCL) Molecular cell layer, (DG) Dentate Gyrus.

**Extended Data Figure 8:**
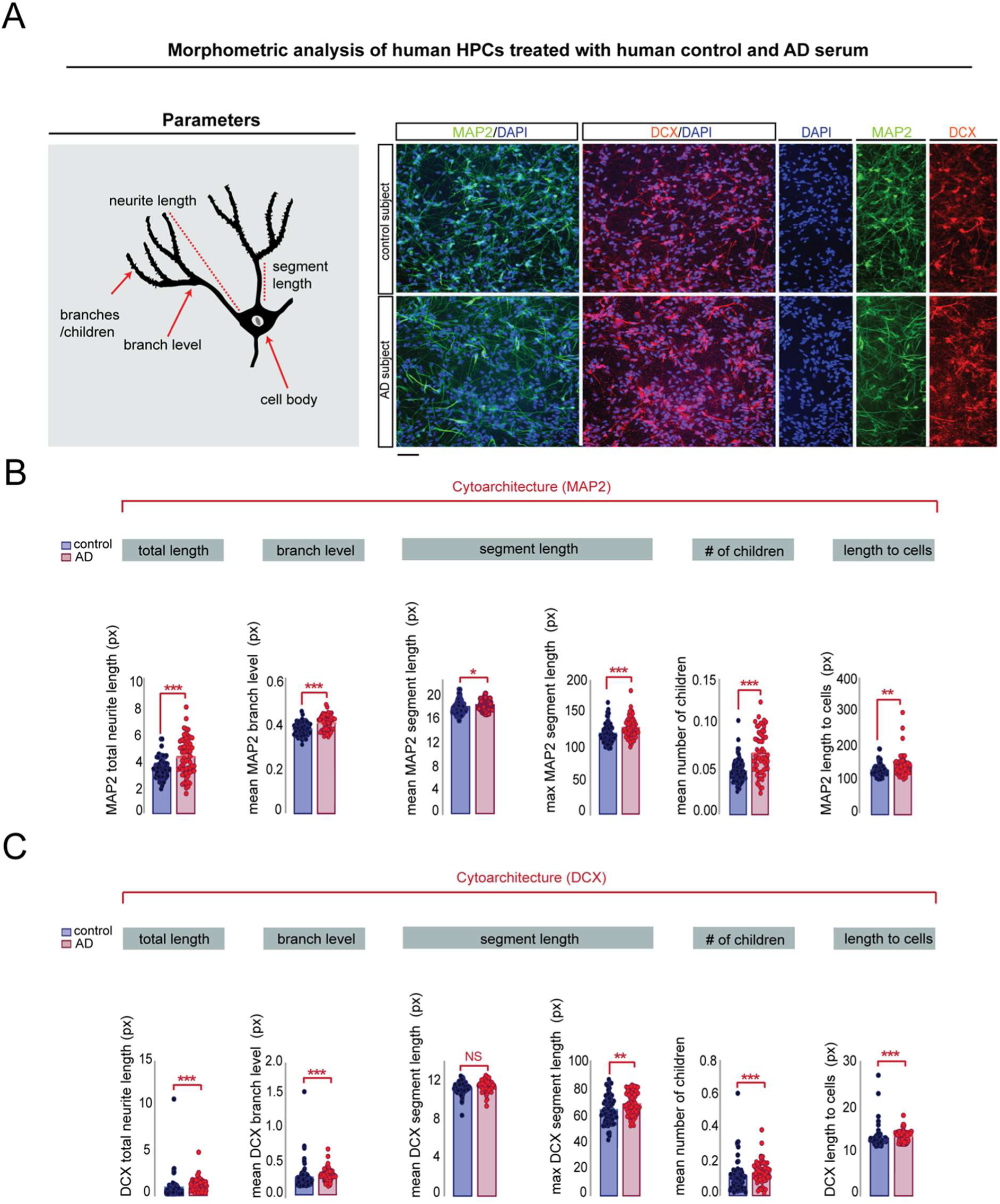
Serum from AD participants modulates dendritic arborization of immature and newly generated neurons. **(A) Upper left panel:** Graphical representation of morphometric parameters. **Upper right panel:** Representative confocal micrographs showing MAP2-positive human HPCs cells (green: MAP2-positive cells, blue: DAPI) and DCX-positive human HPCs cells (red: DCX-positive cells, blue: DAPI) during the differentiation phase of the human serum assay. Scale bar: 100 μm. **(B)** Serum from AD participants induced a significant increase of the total dendritic length, Mann-Whitney test, \*\*\**p*=0.0004 and mean branch levels of immature MAP2 neurons, \*\*\**p*<0.0001. Mean dendritic segment length, Mann-Whitney test, \**p*=0.0131 and max dendritic segment length, Mann-Whitney test, \*\*\**p*=0.0005 were significantly increased in HPCs cells treated with serum from AD participants compared to control subjects. There was a significant increase in the average number of children, Mann-Whitney test, \*\*\**p*<0.0001 and length to cells, Mann-Whitney test, \*\**p*=0.0054 of immature neurons in HPCs cells cultured with serum from AD participants. **(C)** Quantification of DCX positive neurons revealed a significant increase in total neurite length, Mann-Whitney test, \*\*\**p*<0.0001 and mean DCX branch levels, Mann-Whitney test, \*\*\**p*=0.0002. No significant difference was detected, in the mean DCX segment length, Mann-Whitney test, *P*=0.0626. Serum from AD participants induced a significant increase in the max DCX segment length, unpaired, two-tailed Student’s t-test, \*\**p*=0.0083. Serum from AD participants significantly increased the number of children, Mann-Whitney test, \*\*\**p*=0.0004 and decreased the length to cells in human HPCs cells, Mann-Whitney test, \*\*\**p*<0.0001. 15 fields were analysed per well, n=3 technical replicates. Control serum n=73, AD serum n=60. All data are presented as mean ± SEM, \**P*<0.05, \*\**P*<0.01, \*\*\**P*<0.001, Abbreviations: px=pixels.

## Methods

### Human participants

Participants with Alzheimer’s disease (AD) (n=69) and cognitively healthy controls (n=64) were recruited at the IRCCS Centro San Giovanni di Dio Fatebenefratelli, Brescia, Italy. All participants underwent clinical assessment for cognitive function and a physical examination. The Clinical Dementia Rating (CDR), and the Mini Mental State Examination (MMSE) (Folstein et al., 1975), assessed cognitive dysfunctions. Information was collected on diet, drug history, mood and behavior, Height and weight were measured, and body mass index (BMI) was recorded in kg/m^2^. The inclusion criteria for AD were: probable AD diagnosed according to NIA-AA criteria (McKhann et al., 2011), CDR Score ≥0.5, aged between 55 and 90 years, voluntary participation in the project and a signed informed consent. The inclusion criteria for the control participants were: absence of cognitive impairment, aged between 55 and 90 years, voluntary participation in the project and a signed informed consent. Exclusion criteria for both groups of participants were: pathologies of the gastrointestinal tract, chronic inflammatory diseases, tumor history, psychiatric disorders, substantial misuse or dependence on alcohol or substances in the 3 months prior the study, and chronic anti-inflammatory or antibiotic treatment in the 3 months prior to the study. The study was approved by the Ethics Committee of “Comitato Etico dell’IRCCS San Giovanni di Dio – Fatebenefratelli” (Brescia, Italy) under registration number 92/2017. Informed consent to participate in this study was given by all subjects or their caregivers.

### Human biological samples

Venous blood was sampled from an antecubital vein between 08:00–10:00 h and collected in EDTA tubes (BD 186 Vacutainer Systems, Franklin Lakes, NJ) for preparation of plasma. Stool samples were collected from a subset of participants (n=54 for AD and n=41 for controls) at their own home in a sterile plastic cup, stored at −20°C, and subsequently delivered to IRCCS Centro San Giovanni di Dio Fatebenefratelli, for storage at −20 °C until processing. For stool used in fecal microbial transplant (FMT) to rats, frozen samples were transferred into an anaerobic chamber, thawed and homogenized at a 100 mg/ml concentration in reduced sterile phosphate-buffered saline (PBS) containing 20% glycerol (w/v) and filtered through a stomacher bag. The filtered slurry was aliquoted, and frozen at −80°C. Samples were defrosted immediately before inoculating recipient rats.

### Human immune profile

For cytokine expression analysis in plasma, total RNA was isolated using the PAXgene blood miRNA kit according to the manufacturer’s protocol (PreAnalytiX, Hombrechtikon, CHE) and candidate gene expression analyses was performed using real-time PCR. Each target gene was normalized to the expression of three reference genes, glyceraldehyde 3-phosphate dehydrogenase, beta-actin, and beta-2-microglobulin, using Taqman Assays on a 384 wells Real Time PCR System (Biorad). The expression levels of each target gene were normalized to the geometric mean of all three reference genes, and the Pfaffl method was used to determine relative target gene expression of each gene in AD compared to controls.

For the quantitative detection of fecal calprotectin in stool samples, RIDASCREEN® Calprotectin (R-Biopharm Italia S.R.L.) immunoassay was performed on frozen stool samples, thawed immediately before the assay. Accordingly to the manufacterer’s protocol, 10 mg of stool sample was added to RIDA®TUBE Calprotectin pre-filled with 2.5 ml ready-to-use extraction buffer to standardize the starting material quantity. For quantification of calprotectin using a standard curve, positive controls reaching the concentration range specified in the assay kit information were included. Calprotectin quantification was performed according to the 4-parameter logistic log model.

### Animals

Adult male Sprague-Dawley rats (310-350 g) aged 11 weeks, (Envigo UK) were maintained in environmentally controlled conditions at the ambient temperature of 21°C± 2°C, 55±10% humidity, and under a 12-h light-dark cycle. Colonized animals were group-housed (4 per cage) in autoclaved cages with autoclaved bedding (Eco-Pure Aspen chips; Datesand, UK) and had access to autoclaved water and chow (Teklad Global Diet 2018S; Envigo, UK) *ad libitum*. All animal experiments were performed in compliance with the guidelines for the welfare of experimental animals issued by the Health Products Regulatory Authority (HPRA, Ireland), and the European Communities Council Directive (2010/63/EU) and approved by the Animal Experimentation Ethics Committee (#AE19130/P108) of University College Cork.

### Fecal microbiota transplantation

Following acclimatization, rats were administered an antibiotic cocktail of ampicillin (1g/l), vancomycin (500 mg/l), ciprofloxacin HCL (200 mg/l) and imipenem (250 mg/l) for 7 consecutive days in autoclaved drinking water, which has previously been shown to ensure sufficient microbiota depletion (Gheorghe et al., 2021; Staley et al., 2017). Rats were re-colonized 72 hours later via daily oral gavage of donor microbiota (300 μl of 100 mg/ml homogenized fecal slurry) for 3 days (Bruce-Keller et al., 2015). Briefly, each rat was gently restrained, and the fecal slurry was carefully administered through an autoclaved, curved stainless steel gavage needle (16G). During the antibiotic treatment and the initial 3-day primary colonization, rats were transferred to a new autoclaved cage to minimize the carryover of preexisting microbiota from the cage environment. FMT occurred once per day for the first three days to encourage microbiota engraftment, then twice per week throughout the study to offset potential founder/cage effects and reinforce donor microbiota genotype (Bruce-Keller et al., 2015).

### Study design and experimental timeline for animal experiments

After 7 days of antibiotics, rats were randomly allocated into one of the two groups, balanced by weight: rats receiving FMT from control subjects (control FMT, n=16) and rats receiving FMT from AD participants (AD FMT, n=16). To account for biological variation between individual donors, stool samples from 4 human control subjects and 4 human AD participants were gavaged into recipient rats (Fig. 2A). One fecal sample from each human donor was used to colonize 4 microbiota-depleted rats, which were then co-housed within the same cage until the end of the study. Baseline fecal pellets from each rat were collected at the start of the study (after acclimatization), 10 days following primary inoculation and at end of the study to assess the effect of human FMT on recipient fecal microbiota and ensure appropriate engraftment of microbiota throughout the study. Following 10 days of FMT primary inoculation, tail blood samples were taken to measure cytokine expression. Rats were intraperitoneally injected with BrdU (150 mg/kg of body weight, Sigma # B5002) once a day for five consecutive days. Forty-eight hours after behavioral testing to evaluate locomotory, cognitive performance, depression, and anxiety-like behavior (see section on behavioral studies) half of the rats from each treatment group were deeply anesthetized by intraperitoneal injection of pentobarbital (90mg/kg) and then transcardially perfused with 0.9% cooled phosphate-buffered saline (PBS) followed by 4% paraformaldehyde in PBS. Rats were decapitated, the brains were post-fixed overnight in 4% paraformaldehyde, submerged in 30% sucrose in 0.1 M PBS (pH 7.4), frozen in OCT compound and stored at −80°C. The second half of the rats per treatment group were sacrificed by decapitation. Trunk blood, caecum content, and brain samples were collected and frozen at −80°C until subsequent use. The small intestine and colon were opened longitudinally, 2 x 2 cm snips were taken from the distal ileum and colon, frozen in liquid nitrogen, and stored at −80°C for subsequent analysis. The remainder of the colon and ileum was prepared as swiss rolls and fixed in 10% neutral buffered formalin solution. To monitor general health and the effect of FMT, body weight, food consumption, and water intake were recorded weekly.

### Fecal sample collection and fecal water content in rats

Each rat was individually transferred in an autoclaved plastic cage, free of bedding material and sterilized with 70% ethanol. Fecal pellets for 16s microbiota analysis were collected and snap-frozen on dry ice before being stored at −80°C until analysis. For analysis of faecal water content, faecal pellets were collected immediately after expulsion and placed in sealed 1.5 ml tubes to avoid evaporation. Pellets were weighted to obtain wet weight and dried at 65°C for 24 h. Stool water content was calculated as faecal water content (%) = ((faeces wet weight – faeces dry weight)/ faeces wet weight) x100.

### 16S microbiota and bioinformatic analysis of human and rat samples

DNA was extracted from 180-200 mg of frozen stool using the QIAamp DNA Stool Mini Kit (Qiagen Retsch GmbH, Hannover, Germany) according to the manufacturer’s instructions. Bead-beating homogenization by TissueLyser II (Qiagen Retsch GmbH, Hannover, Germany) was performed to mechanically disrupt fecal samples before DNA extraction. The samples were homogenized for 10 minutes at 30 Hz. DNA was quantified using a NanoDrop ND-1000 spectrophotometer, and then stored at +4°C until subsequent analyses. The regions V3 and V4 of the bacterial ribosomal RNA 16S gene were amplified and purified according to 16S Metagenomic Sequencing Library Preparation protocol by Illumina. A paired-end sequencing of 300 cycles per read was performed. Animal samples were analyzed in a single sequencing run and human samples in three independent sequencing runs. The raw 16S data were processed using QIIME2 (Bolyen et al., 2019) (64-bit version 2021.4) run on MacBook Pro with Intel CPU 12 x 2.6 GHz processors and 16 GB of RAM. Sequencing Illumina MiSeq data were already demultiplexed. Forward and reverse primers, reads containing ambiguous bases or homopolymers greater than eight base pairs in length were removed. A maximum number of expected errors equal to 2 and reads truncation if the quality score was less than 2 was set up. The DADA2 denoising process was applied with the default parameters for primer removal, quality control and amplicon sequence data correction. Taxonomy was assigned with DADA2 against the SILVA reference database release (v.138) (Quast et al., 2013). ASVs were aggregated at genus level. Genera with incomplete taxonomy, assigned to the Archea domain or found in less than 10% of the subjects across all samples were removed. After filtering, 105 different genera were left in total over all microbiome samples in rats and 99 in humans. Alpha and beta diversities were calculated using the q2-diversity plugin and included Bray-Curtis dissimilarity, Shannon and Pielou’s evenness indexes. The feature table was rarefied to the sample depth corresponding to the sample with the lowest read count.

Statistical analyses were performed in R (v.3.6.3) with the Rstudio GUI (v.1.2.5033) except where otherwise specified. For human data, beta diversity was analyzed using PERMANOVA with covariates (age, run) performed using the adonis function for the Vegan (2.6-2) package (Oksanen J, Simpson G, Blanchet F, Kindt R, Legendre P, Minchin P, O’Hara R, Solymos P et al., 2022) and alpha diversities were analysed using one-way ANOVA adjusted for age and run after performing normality tests. For differential abundance analysis, a model-free data-normalization procedure for controlling batch effects in case-control microbiome studies was applied using the q2-perc-norm plugin (Gibbons et al., 2018) (as a compatible q2-perc-norm version for release 2021.4 was not available, the release 2019.7 was used to install this plugin). q2-perc-norm plugin uses control samples to normalize participant samples to mitigate batch effects. Conversion to percentiles of controls was carried out within each run and normalized case and control samples from the different runs were combined together into the same ASVs table before statistical testing. We used the Wilcoxon rank-sum test to determine significant differences between AD participants and controls. P-values were multiple-test corrected using the Benjamini-Hochberg False Discovery Rate (FDR) procedure and a cutoff of 0.1 was applied to select the differentially abundant genera. For beta diversity analysis in rats, first distance followed by Linear mixed-effects model were used to track temporal changes (between successive time-point) in rats’ beta diversities and were calculated within QIIME2, using the q2-longitudinal plugin (Bokulich et al., 2018), by setting time and FMT as fixed effects and random intercepts (rat ID) as random effect. Alpha diversity changes were assessed by using repeated measurements ANOVA after performing normality tests (R packages: tidyverse v1.3.1, ggpubr v0.4.0, rstatix v0.7.0), and differential abundance of microbial genera using the Wilcoxon signed-rank test implementation in the ALDEx2 library (Fernandes et al., 2014). FDR correction was applied using the internal ALDEx2 implementation of the Benjamini–Hochberg post hoc procedure with a q-value of 0.1 as a cutoff. To assess taxa engraftment in rats we calculated the genera from 16S rRNA gene sequencing present in rats and respective donors and gain and loss genus events as the occurrences of a genus present (or absent) at D10 and D59 relative to the time-point prior to antibiotic treatment. Difference in means were tested by two-way ANOVA with repeated measures with donor diagnosis as between-subject factor and time from the first FMT (D10, D59) as within-subject factor. Microbiota figures were generated using ggplot2 except the stacked bar plots showing the individual composition of the microbiome (Extended Data Fig. 2F) for which GraphPad Prism was used.

### Histological preparation of ileal and colonic sections from rats

Intestinal parts were opened longitudinally along the mesenteric line, flattened, and thoroughly rinsed in PBS before being separately rolled with the mucosa inwards (Bialkowska et al., 2016). These Swiss-rolls were placed in an embedding cassette containing 10% buffered formalin for fixation overnight. Then, 24 h after fixation, samples were washed with PBS and dehydrated through a graded series of ethanol with a histokinette (Leica TP1020 Tissue Processor). Tissue samples were subsequently embedded in paraffin blocks using the Tissue-Tek^®^ Tec^TM^5 apparatus (Sakura), and 5 μm sections were cut using a microtome (Leica RM2135). Pre-cut sections were floated on pre-coated glass slides in a 40°C water bath and dried overnight at 60°C.

#### Hematoxylin and eosin (H&E)

H&E staining was performed for morphological analysis using standard histological protocols. Briefly, ileum and colon sections were deparaffinated in histolene and rehydrated in a descending series of ethanol (100%, 96%, 70%) followed by hematoxylin solution (H&E, Merck, Darmstadt, Germany) for 5 min at room temperature. After washing, slides were counterstained with 0.1% eosin for 30 s (H&E, Merck, Darmstadt, Germany), followed by dehydrating steps of an ethanol series (96%, 100%) and histolene incubation. Finally, tissue sections were coverslipped using DPX mounting medium andimaged at a 4x,10x, and 20x magnification on an Olympus BX40 light microscope. Crypt depth and villi length were analyzed at 10x magnification. Only well-oriented with longitudinally cut crypts were evaluated. Villus length was determined by measuring the distance from the base to the top of the villi. Crypt depth was measured between the crypt-villus junction and the base of the crypt using ImageJ 1.50i. 100 complete villi and 170 open crypts per rat were analyzed in a blinded fashion.

#### Alcian Blue/Periodic acid-Schiff (PAS) staining

Goblet cell staining and quantification was performed using the standard Periodic Acid-Schiff staining protocol. After deparaffinization in histolene and rehydration in a series of ethanol (2x 100%, 95%, and 70%), tissue sections were washed under running tap water. Tissue sections were placed in 3% acetic acid for 3 min, followed by incubation in alcian blue solution (pH 2.5, dissolved in 3% acetic acid) for 30 min. Sections were then rinsed in 3% acetic acid for 10 s. After washing, sections were incubated in 0.5% periodic acid for 5 min, washed, and stained with Schiff reagent (Fisher Scientific, #1713072) for 10 min. After another wash step, sections were counterstained in nuclear fast red solution (Sigma Aldrich, #N3020) for 10 min. After a final washing step, sections were dehydrated in ethanol (95%, 100%) and histolene and finally mounted using DPX mounting medium. The number of PAS-positive cells was analyzed at 20x magnification on an Olympus BX40 light microscope. The numbers of PAS-positive cells per crypt and villi were determined by analysis of 50-80 well-oriented crypts and villi per animal using ImageJ 1.50i in a blinded manner.

### Behavioral testing in rats

Behavioral testing started 10 days after donor colonization and were conducted between 9 am and 7 pm during the light phase of the light cycle with a minimum of 36 h between each behavioral test. At least one hour before each behavioral experiment, all rats were acclimated to the experimental room (illumination 150-180 lux) to reduce stress and anxiety. Cages of different groups were alternated to control for potential time of days effects. The experimenter was blinded to the FMT human donor but not to the rat group.

#### Fecal output (FO)

Rats were removed from their home cages and placed individually in a circular open field arena (diameter 90 cm). Fecal pellets were counted every 5 min, for a cumulative time of 15 min (Sampson et al., 2016).

#### Open Field (OF)

Spontaneous locomotor activity and exploratory behavior were examined over 15 min in a circular (diameter 90 cm) open-field arena. The arena was evenly illuminated by overhead white lighting (1000 lux) and constructed of white Plexiglas (diameter centre zone: 45 cm). Total distance travelled (cm), average velocity, time spent in the center zone vs. time spent at the border zone, number of entries into the center zone and border zone were quantified using specialized software with a tracking system (Ethovision XT 8.5, Noldus Information Technology, USA). The open-field arena was cleaned with 70% ethanol between each rat to prevent odor traces.

#### Elevated Plus Maze (EPM)

Anxiety-related behavior was examined using the elevated plus-maze. The maze was made of white Plexiglas, with two open (50.5 x 10.5 cm) and two enclosed arms (50.5 x 10.5 x 40 cm), extending from a central platform (10.5 x 10.5 cm). The apparatus was elevated 60 cm above the floor on a central pedestal (75 cm; Cat # ENV-560; Med Aossciates Inc). Rats were placed in the central arena, facing one of the open arms, and allowed to explore the setting for 5 min. Time spent in the open and closed arms, the number of entries into the open and closed arms, and average velocity, were analyzed using specialized software with a tracking system (Ethovision XT 8.5, Noldus Information Technology, USA).

#### Modified spontaneous location recognition test (MSLR)

To evaluate the ability of rats to discriminate similar memory representations (a hippocampal neurogenesis-dependent task), rats were tested in a pattern separation task. In this test, rats undergo two consecutive location discrimination tasks (large separation and small separation) where the inter-stimulus distance between the novel and familiar locations has been varied. The test was carried out in an open field arena (90 cm diameter) covered with bedding under dim lighting conditions as previously described (20 lux) (Hueston et al., 2018). The testing room had three proximal cues and distal standard furniture cues for spatial navigation. Objects used for testing were cylinders of approximately 20 cm height (soda cans, glass beer bottles). Objects were secured with Blu-tack^TM^ on the bottom of the open field arena to prevent movement during object exploration. Between each session, objects were carefully cleaned with 70% ethanol to prevent odor traces. Rats were first habituated to the empty open field arena for 10 minutes per day for 5 consecutive days. 24 h after the last habituation day, rats were placed in the arena for 10 minutes and allowed to explore 3 identical objects. For the small separation paradigm, 2 of the objects (A2 and A3) were separated by a 50° angle, and the third (A1) was at an equal distance between the two. 120° angles separated the 3 objects for the large separation paradigm (Figure 3B). Twenty-four hours after acquisition, animals were presented with 2 of the previously used objects, one (A4) placed in the same position as A1, and the second (A5) placed halfway between the acquisition locations of A2 and A3. Rats were allowed to explore the objects for 5 minutes. The amount of time spent by each animal exploring the object in the novel location (A5) and the familiar location (A4) was recorded, and a discrimination ratio was calculated: Novel Exploration time/(Novel + Familiar Exploration time). Object exploration was defined as a clear nose contact with the object. A digital camera recorded the testing sessions.

#### Novel Object recognition and Novel Object location test (NOR, NOL)

To evaluate recognition memory, rats were tested in the novel object recognition test in an open field arena (diameter: 90 cm). The test was performed in three phases: habituation, acquisition, and retention. Rats were first habituated to the open field arena (without any object inside) under dim light (20 lux) for 10 min. Twenty-four hours later, rats were exposed to two identical copies of the same objects, placed in the two corners of the arena, approximately 4 cm from the sidewall (Figure 3E). The rats were placed back into the same arena facing the wall on the opposite side of the objects and allowed to freely explore for 10 min. After this acquisition period, rats were placed back into the home cage. Following a 24 h interval, rats were transferred back to the arena where a novel object replaced one of the identical objects. The rats were allowed to explore for 10 min. Recorded videos were scored manually with a stopwatch. Object exploration was defined as a clear nose contact with the object. To measure recognition memory, a preference index for the novel object was calculated as the ratio of the time spent exploring the novel object over the total time spent exploring both objects: (discrimination index [DI] = novel object exploration x 100/(novel object exploration + old object exploration). To evaluate spatial memory, rats were exposed to two identical objects placed in the left and right border zone of the open field arena (approximately 4 cm) (Figure 3F). Rats were allowed to freely explore the objects for 10 min. After this acquisition period, rats were placed back in the home cage. Following a delay of 24 h, the position of one object was changed to the opposite corner of the open-field arena. Time spent exploring the object that has changed position was compared with time spent exploring the object in the old location. Analysis was carried out as described above. The location of the moved object was counter-balanced between rats.

#### Morris water maze (MWM)

To assess hippocampal-dependent spatial learning and memory, rats were tested in the MWM as previously described (Hoban et al., 2016). In brief, a black pool (180cm diameter) with opaque water maintained at 25°C ± 2°C was used. The MWM was located in the behavior room with distal cues at several locations to allow for spatial navigation. Animals were trained (acquisition day 1-4) to find a hidden platform (10cm diameter) for 1 min located in one quadrant of the pool (Figure 3C). If they found the platform within 1 min, they were allowed to stay on it for 15 s. If they did not find the platform within 1 min, they were guided by hand to the platform and allowed to stay on it for 15 s. Rats were subjected to four trials per day. The times recorded during the four trials were averaged to calculate each rat’s escape latency per day. For each trial, the starting point varied randomly. For the probe trial (day 5), the platform was removed, and the rat was placed in a novel start position in the maze, facing the tank wall. Rats were removed after 60 seconds. Time spent in each quadrant, swim speed, path length and the number of platform crossings were analyzed using Noldus EthoVision XT 10 software.

#### Forced swim test (FST)

Antidepressant-like behavior was assessed using the FST as previously described (Kozareva et al., 2019). The test consisted of a pre-swim test and a test session. Rats were individually placed in a Pryrex cylinder (Fischer Scientific: 21 cm diameter x 46 cm height, filled to 30 cm mark with 25°C water) for 15 min. 24 h after the pre-swim tests, rats were placed back into the cylinder. A video camera recorded behavior during the pre-swim and test session for subsequent analysis. The cylinder was emptied and replaced with fresh water between each animal exposure. A time-sampling technique was employed whereby the predominant behavior (immobility, swimming, and climbing) in each of the 5-second time frames was scored. Climbing was defined as upward-directed movements of the forepaws along the side of the cylinder. Swimming was assigned when the rats moved horizontally through the cylinder crossing quadrants. Immobility was defined as no additional movement of the rat to keep its head afloat. Scores are presented as cumulative for each trial. Behavior was recorded from the side as well as from above the tank.

### Immunohistochemical and histochemical analysis of rat brain tissue

Immunohistochemistry was performed for neuroblast detection and analysis of hippocampal neurogenesis using the following primary antibodies diluted in PBS containing 5% goat serum and 0.1% Triton-X-100: anti-doublecortin (DCX, rabbit: 1:5000, Abcam, #ab18723), anti-bromodeoxyuridine (BrdU, rat: 1:100, Abcam, #ab6326) and anti-neuronal nuclear antigen (NeuN, mouse: 1:200, Millipore, #MAB3771). The number and morphology of microglia in the hippocampi were assessed by staining for ionized calcium-binding adaptor molecule 1 (Iba1, rabbit: 1:1000, Wako, #01919741). Appropriate secondary antibodies were coupled to Alexa Fluor^®^ 488, 555, 635 (dilution: 1:500, Life Technologies). Fixed hippocampal tissue was coronal sectioned (40 μm) using a cryostat (Leica CM_3050_ S, UK), transferred to PBS, and blocked (3% goat serum + 0.1% Triton-X-100, diluted in PBS) for 2h at RT on a horizontal shaker. Sections were then incubated for 48 h at 4°C with primary antibodies and subsequently washed 3x in PBS for 15 min at RT. After washing, sections were incubated for 2 h at RT with fluorophore-conjugated secondary antibodies. Finally, coronal sections were rewashed 3x in PBS for 15 min at RT, mounted and coverslipped using fluorescent mounting medium (DAKO, S3023) containing diluted 4,6-diamidino-2-phenylindole DAPI (dilution 1:50000). For double-labeling of BrdU and NeuN positive cells, hippocampal sections were washed with PBS and pre-treated with 2 N HCl at 37°C for 45 min and renatured in 0.1 M sodium tetraborate (pH 8.5) before staining.

#### Cell density of new neurons and three-dimensional (3D) reconstruction of DCX+ immature neurons

The number of BrdU/NeuN and DCX positive cells in the dentate gyrus (DG) was counted in a 1:6 series of sections through the hippocampus. The area of the DG in each section was determined by measuring the DAPI-stained region of the DG. Cell quantification and area measurement were performed using the image processing software ImageJ. Cell densities were normalized to the size of DG to obtain the number of cells per mm^2^. DCX positive cells were visually classified based on dendritic morphology according to the classification schemata of (Plümpe et al., 2006). Images were obtained using an Olympus BX40 fluorescent microscope coupled to an Olympus DP72 camera at a 20x magnification. For morphometric analysis of DCX positive cells, only postmitotic (EF-type neurons) were selected. 3D dendritic reconstructions of dendritic trees were created from confocal stack images of EF-type neurons (n=56-72 neurons per group) that reach the molecular layer (ML) and project into the granular cell layer (GCL). Images were obtained using Olympus® Fuoview FV10i confocal laser scanning microscope with a 60x water immersion objective (1.3 zoom). Stacks were prepared with a z-step of 0.47 μm in the z-direction and 1024 x 1024-pixel resolution. 3D reconstructions were created using the simple neurite tracer plugin (Longair et al., 2011) on the image processing software Fiji (Schindelin et al., 2012). The 3D Sholl analysis plugin (http://fiji.sc/Sholl_Analysis) was used to analyze dendritic complexity by quantifying the number of dendrites that crossed a series of concentric circles at a 20 μm interval from the cell soma. For the analysis, 7-9 individual DCX-positive cells were reconstructed from each rat (n=8 per group). The experimenter was blinded to the treatment groups.

#### Microglia cell density and morphological characterization

For the analysis of microglia density, Iba1-positive stained hippocampal sections were imaged using an Olympus BX40 upright microscope coupled to an Olympus DP72 camera with 10x magnification. The DG was imaged with a 10x objective on 5 sections per animal (n=8). Quantification of Iba1-positive cells in the left and right DG was performed using the ImageJ plugin “Analyze particles.” The subdivisions and respective areas of the DG (hilus, GCL, MCL) were identified using DAPI staining and measured with the image processing software ImageJ. The density of Iba1-positive cells in the DG is reported as number of cells per mm^2^. For the analysis of Iba1 cell soma size, imaging was performed on an Olympus® Fuoview FV10i confocal microscope using a 40x objective. Z stacks were prepared in a 0.47 μm step in the z-direction, 1024×1024 pixel resolution. Ten to fifteen randomly selected cells were sampled per section and 5 to 6 sections were analyzed per animal (n=8 per group). The area of the soma was measured using ImageJ and expressed as μm^2^. For microglial classification, Iba1+ cells with soma area equal to or below one standard deviation from the mean of the control group were categorized as ‘small soma size’ while cells with a soma area greater than this mean were categorized as ‘large soma size’ (Kozareva et al., 2019). The experimenter was blinded to the treatment groups.

#### Thioflavin S staining

Free-floating hippocampal and cortex sections were transferred to PBS for 15 min on a horizontal shaker. After washing, sections were incubated in a series of ethanol (80%, 70%, and 50%) for 1 min each. Sections were subsequently stained with filtered thioflavin S solution (0.1% dissolved in 50% ethanol) for 25 min at RT in the dark. Afterwards, sections were washed in a 1-min series with 50%, 70%, and 80% ethanol followed by three washes with PBS. Finally, sections were counterstained with DAPI, washed in distilled water before being cover slipped using DPX mounting medium. Image acquisition was performed with an Olympus BX40 fluorescence microscope. The experimenter was blinded to the treatment groups.

### Cell culture and serum treatment

All experiments were performed using the multipotent human hippocampal progenitor/stem cell line HPC0A07/03C (HPC, ReNeuron, UK) derived from the first trimester female foetal hippocampal tissue following medical termination (in accordance with the UK and USA ethical and legal guidelines, and obtained from Advanced Bioscience Resources (Alameda CA, USA)). HPC0A07/03C cells were conditionally immortalized by introducing *c-mycER^TAM^*transgene enabling them to proliferate indefinitely in the presence of epidermal growth factor (EGF), basic fibroblast growth factor (bFGF) and 4-hydroxy-tamoxifen (4-OHT) (Pollock et al., 2006). Removal of these three factors induces spontaneous differentiation into neurons, astrocytes or oligodendrocytes (Anacker et al., 2013, 2011; Zunszain et al., 2012).

For the *in vitro* neurogenesis assay, HPCs were seeded at passage 19 at a density of 13,200 cells/well in NUNC™ 96-well plates (ThermoFisher, #167008). Cells were treated with 1% participant serum 24 hours post-seeding for the proliferation assay; and for the differentiation assay cells were treated with serum 24 hours post-seeding in proliferation medium, and one more time 3 days post-seeding in differentiation medium (Fig. 5A). See (Anacker et al., 2011) and Supplementary Table 1 for cell culture medium composition. Control conditions consisted of either proliferation or differentiation medium supplemented with 1% Penicillin-Streptomycin (10,000 U/mL ThermoFisher, 15140122). For each experiment, there were technical triplicates. The coefficient of variation for each marker was below 20%, apart from CC3 (below 30%), calculated across different plates and batches of experiment.

**Supplementary Table 1:**
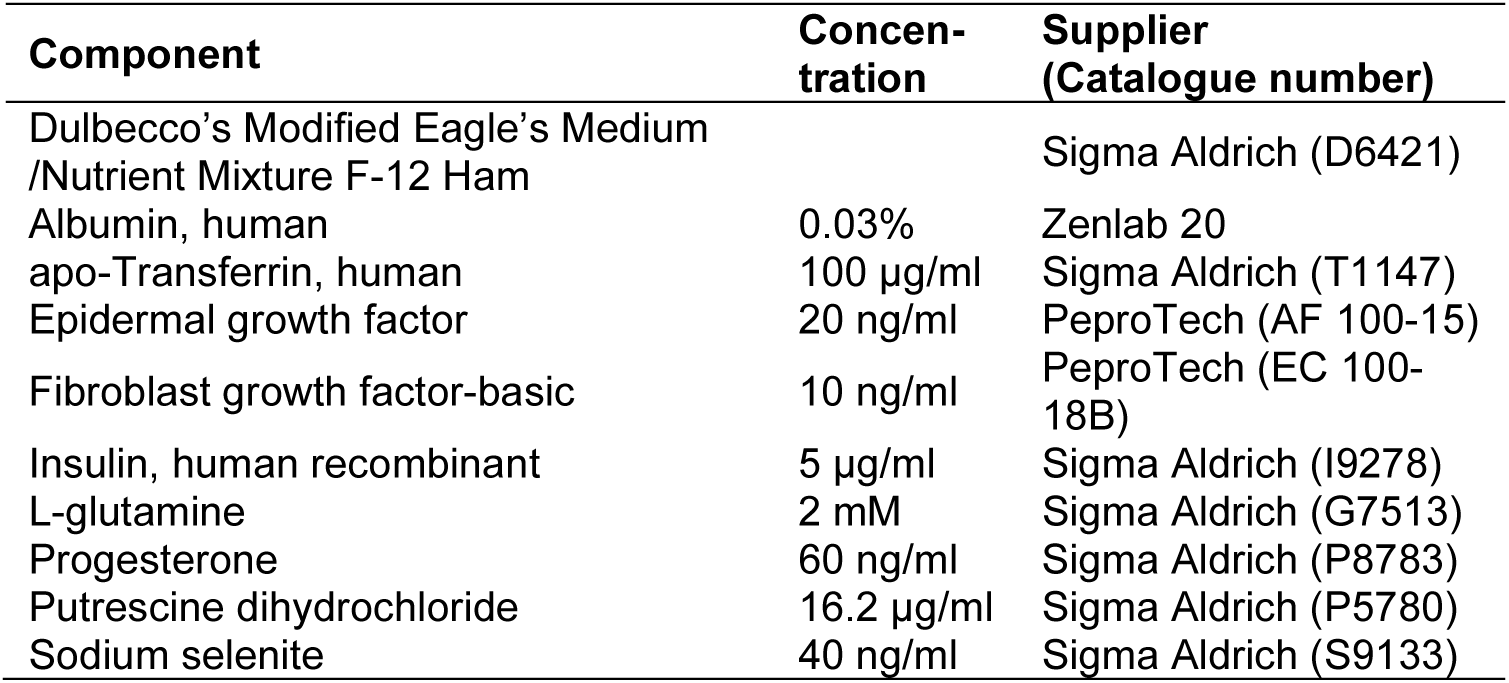
Cell culture medium components

| Component | Concentration | Supplier (Catalogue number) |
| --- | --- | --- |
| Dulbecco's Modified Eagle's Medium /Nutrient Mixture F-12 Ham |  | Sigma Aldrich (D6421) |
| Albumin, human | 0.03% | Zenlab 20 |
| apo-Transferrin, human | 100 µg/ml | Sigma Aldrich (T1147) |
| Epidermal growth factor | 20 ng/ml | PeproTech (AF 100-15) |
| Fibroblast growth factor-basic | 10 ng/ml | PeproTech (EC 100-18B) |
| Insulin, human recombinant | 5 µg/ml | Sigma Aldrich (I9278) |
| L-glutamine | 2 mM | Sigma Aldrich (G7513) |
| Progesterone | 60 ng/ml | Sigma Aldrich (P8783) |
| Putrescine dihydrochloride | 16.2 µg/ml | Sigma Aldrich (P5780) |
| Sodium selenite | 40 ng/ml | Sigma Aldrich (S9133) |

#### Immunocytochemistry

Cells were fixed in 4% paraformaldehyde (PFA) (Alfa Aesar, #43368.9M) after 48 hours of serum treatment for the proliferation and 7 days of treatment for the differentiation phase of the assay. Briefly, cells were washed once with 37°C PBS and fixed in 4% PFA at RT for 20 minutes (50 µl/well), then washed twice with PBS for storage at 4°C prior to immunocytochemistry (ICC). On the day of ICC, cells were first blocked in ‘5% normal donkey serum + 0.3% Triton X-100’ in PBS (i.e., blocking solution) at RT for 1 hour (50 µl/well). Upon removal of blocking solution, cells were incubated with diluted primary antibodies overnight at 4°C (30 µl/well). Cells were then washed with PBS (150 µl/well) twice and incubated with secondary antibodies (diluted 1:500) at RT for 2 hours (30 µl/well) covered from light. Upon removal of secondary antibodies, cells were washed with PBS (150 µl/well) twice and incubated with DAPI nuclear stain (Sigma Aldrich, #D9542) at RT for 5 mins (50 µl/well). Finally, cells were washed with PBS (150 µl/well) twice and stored at 4°C with 0.05% sodium azide in PBS (200 µl/well) prior to imaging. All primary and secondary antibody solutions were made in blocking solution (as described above). Mouse monoclonal anti-Ki67 (Cell Signaling, #9449, 1:800) was used to assess proliferation; rabbit monoclonal anti-CC3 (Cell Signaling, #9664, 1:500) to assess apoptotic cell death; rabbit polyclonal anti-DCX (Abcam, #ab18723, 1:500) for neuroblasts; and mouse monoclonal anti-MAP2 (Abcam, #ab11267, 1:500) for mature neurons. Secondary antibodies were conjugated with either Alexa 488 (Thermo Fisher Scientific, #A21202) or Alexa 555 (Thermo Fisher Scientific, #A31572) fluorophores. Nuclei were counterstained with DAPI (Sigma Aldrich, #D9542). See Supplementary Table 2 for antibody information.

**Supplementary Table 2:** Antibody information for immunocytochemistry on cell cultures

| **Antibodies** |  |  |
| --- | --- | --- |
| Mouse monoclonal anti-Ki67 | Cell Signaling Technology | Cat# 9449,<br>RRID:AB_2797703 |
| Rabbit monoclonal anti-CC3 | Cell Signaling Technology | Cat# 9664,<br>RRID:AB_2070042 |
| Rabbit polyclonal anti-DCX | Abcam | Cat# ab18723,<br>RRID:AB_732011 |
| Mouse monoclonal anti-MAP2 | Abcam | Cat# ab11267,<br>RRID:AB_297885 |
| Alexa 488-conjugated donkey anti-mouse IgG (H+L) | Thermo Fisher Scientific | Cat# A-21202,<br>RRID:AB_141607 |
| Alexa 555-conjugated donkey anti-rabbit IgG (H+L) | Thermo Fisher Scientific | Cat# A-31572,<br>RRID:AB_162543 |

#### High-content imaging

Semi-automated quantification of cellular phenotypes was performed using the CellInsight™ CX5 High Content Screening (HCS) Platform (ThermoFisher). The “Cell Health Profiling” application was used to detect the nucleus (DAPI) and to quantify neurogenic markers expressed in the nucleus (Ki67), and the cell body/dendrites (CC3, DCX and MAP2) (HCS Studio™ Cell Analysis Software, ThermoFisher). Based on the values from positive and negative staining controls, thresholds were set for average intensity within the target regions of interest (e.g., nuclear or cell body). Any cell with an average intensity greater than the threshold was deemed positive for a given neurogenic marker. Fifteen fields were scanned per well of a 96-well plate. A representative protocol used for cellular phenotyping is shown in Supplementary Table 3.

**Supplementary Table 3:**
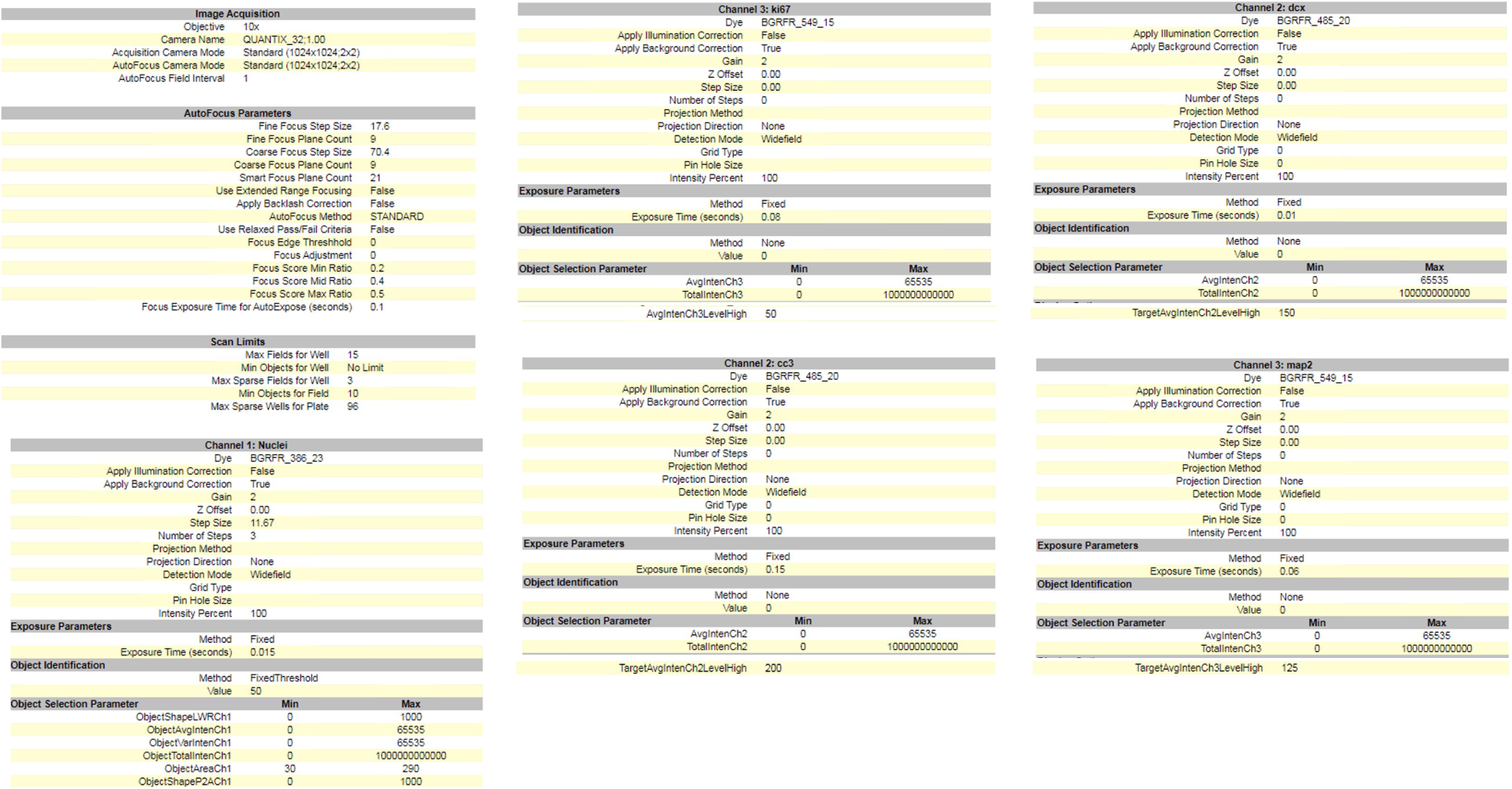
Cellular phenotyping protocol on Thermo Scientific™ CellInsight™ CX5 High Content Screening Platform: Parameters used for semi-automated quantification of DAPI (nuclear), Ki67, CC3, DCX, and MAP2 are shown.

#### Neurite outgrowth

The images for neurite outgrowth analysis were taken on day 7 of differentiation using CellInsight High Content Imaging Platform (ThermoFisher, CX7A1110/CX7B1110) and then analysed using Columbus (Perkin Elmer). For neurite outgrowth analysis, DAPI was used to visualise the nuclei of the cells, DCX marked neuroblasts in early differentiation and migration, and MAP2 marked more mature cells in the later stages of differentiation (Supplementary Figure 1). In the Columbus image analysis system, a protocol was created to optimize the detection of DCX- and MAP2-positive cells by adjusting parameters, regarding cell size, shape and contrasts (de Lucia et al., 2020).

**Supplementary Figure 1:**
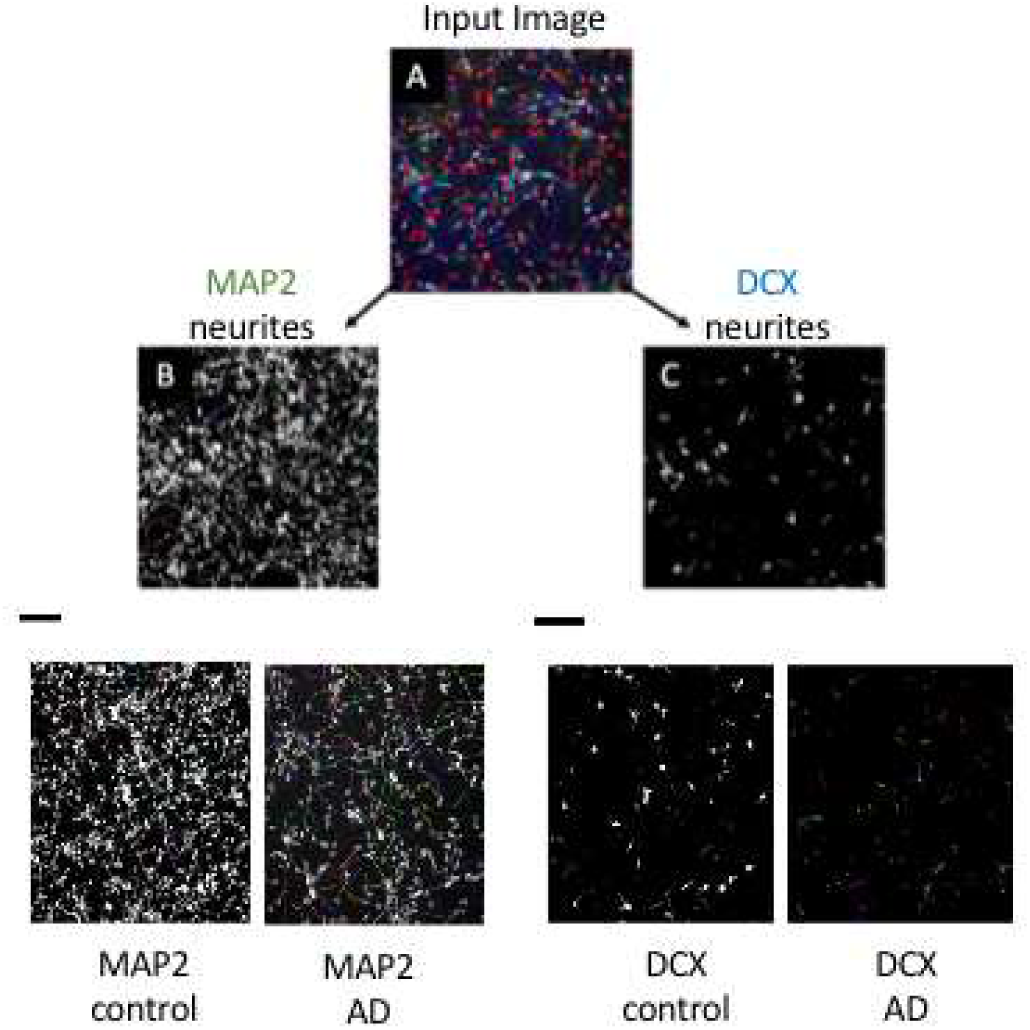
Representative example of the detection of neurites in Columbus imaging software. Images from CellInsight were analysed with the Columbus imaging software, in which settings were optimised to detect cell bodies and neurites. These settings were applied to each image (e.g. A) to detect MAP2-positive neurites (B) and DCX-positive neurites (C).

This protocol was designed using both positive and negative controls, namely images with large numbers of cells and images with very few cells, respectively. The goal of this protocol was to detect as many neurites present as accurately as possible. Picking up spurious elements and interpreting them as neurites was carefully avoided, and as such, the final protocol was conservative in picking up neurites. The detection of MAP2-positive and DCX-posive neurites from an input image is schematically shown in Supplementary Figure 1. Once the protocol was created and adjusted, the finished protocol was applied to analyse the cells in all images. This final protocol with the optimised settings can be found in Supplementary Figure 2.

**Supplementary Figure 2:**
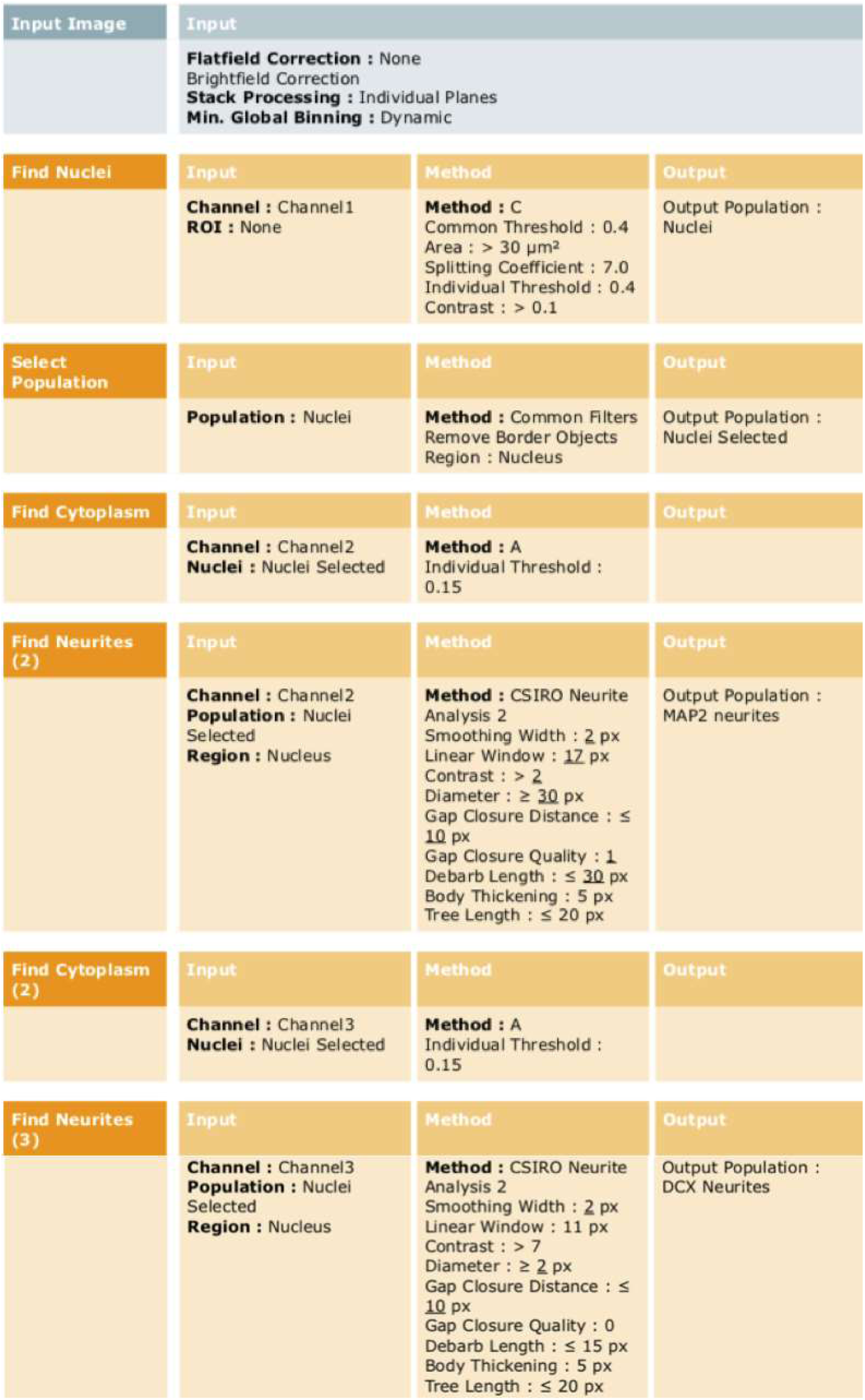
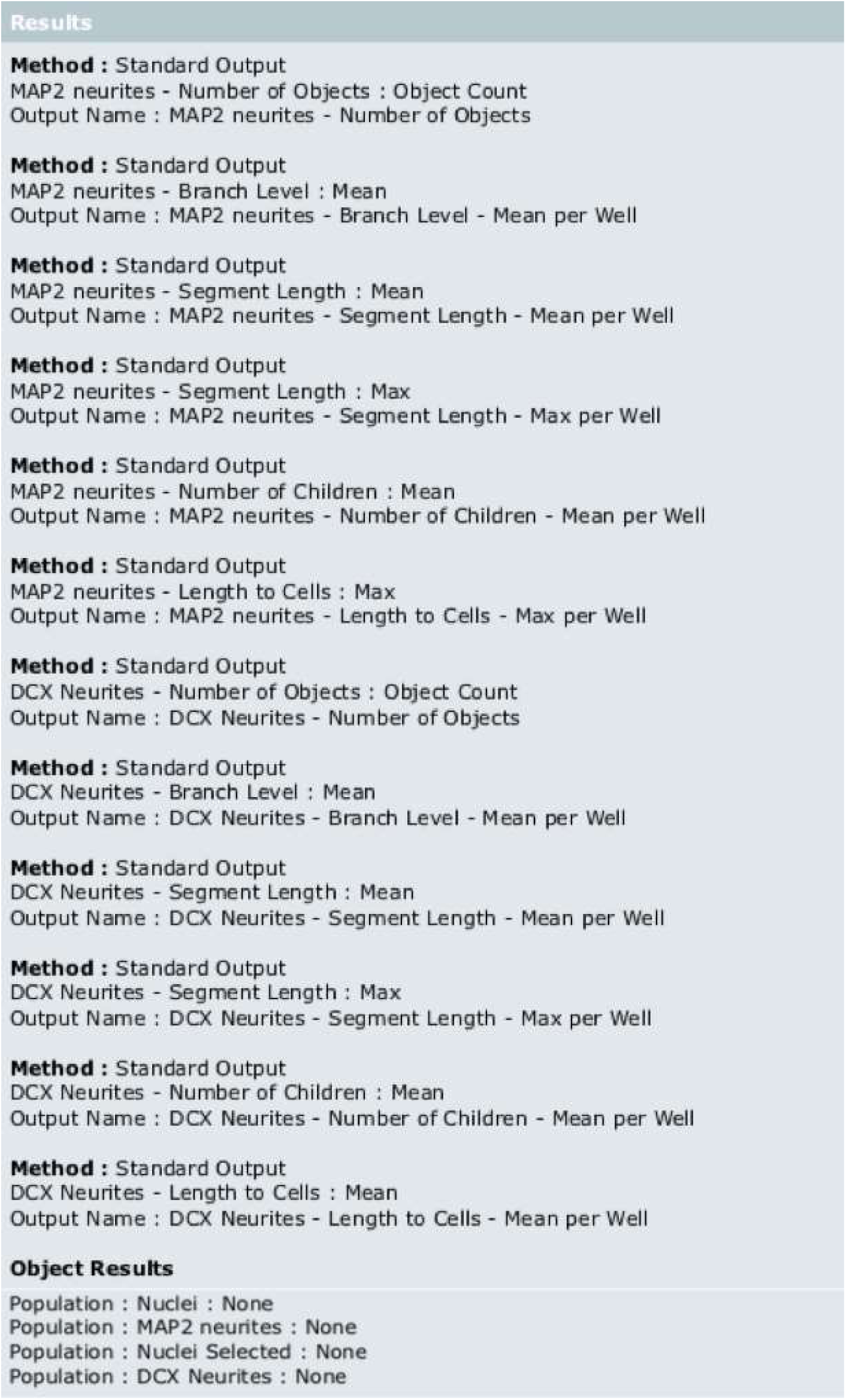
Columbus image analysis protocol

The parameters measured were ‘number of cells’, ‘total neurite length’, ‘segment length’, ‘branch level’, ‘number of children’ (number of attached neurite segments) and ‘length to cells’ (distance of the shortest path from the outer end of the neurite segment to the cell body. It is equal for the segment length for segments directly attached to the cell body) (Supplementary Figure 3).

**Supplementary Figure 3:**
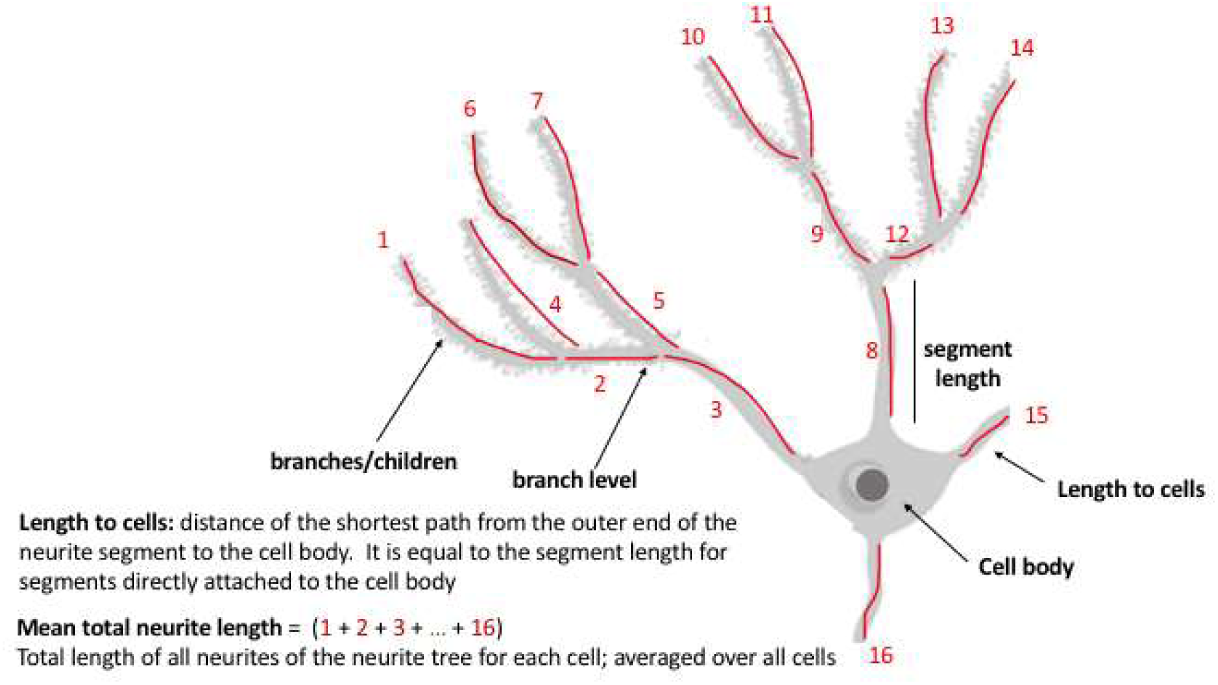
Schematic image analysis parameters for neurite outgrowth

### Rat Hippocampal and Caecal metabolomics

*Preparation of hippocampal tissue*: Frozen hippocampal tissue samples were transferred to homogenization tubes containing 300 μl ice-cold MeOH:MQW (1:2 v/v) and stainless steel beads. Samples were then placed in precooled blocks (−20°C) and homogenized for 3 x 30 s. Beat beating was followed by further disruption of the hippocampal tissue in an ultrasonic bath for 10 min. Samples were centrifuged for 10 min (12 000 g, 4°C). and 300 μl of the supernatant was collected and transferred to a new Eppendorf tube, and the residue was re-extracted with 200 μl MeOH:MQW (1:2 v/v) following the same procedure as described above. The two supernatants were combined, 200 μl chloroform: methanol (1:3,v.v) was added, and the samples subsequently vortexed. Next, 350 μl of the top (polar) layer was transferred to a new Eppendorf tube, and the samples were dried under a light flow of N2. The dried samples were resuspended in 150 μl resuspension mix eluent A1 (10mM ammonium formate, 0.1% formic acid dissolved in water) and eluent B1 (10 mM ammonium formate, 0,1% formic acid dissolved in methanol), mixed 9:1 (v/v) and transferred to a labeled spin-filter tube and filtered at 3000 g for 5 min. In a final step, samples were diluted five times using eluent A1 (10mM ammonium formate, 0.1% formic acid dissolved in water).

*Preparation of caecal content*: Caecal content was added to 4xwt/vol MQQW (MilliQ-water), immediately vortexed, and centrifuged for 10 min (16000 g, 4°C). After centrifugation, the supernatant was collected and filtered by centrifugation (5 min at 15000 g, 4°C) using spinX filters. Finally, samples were diluted five times in an eluent A1 (10 mM ammonium formate, 0.1% formic acid dissolved in water).

#### Mass spectrometry and feature annotation

Analysis of caecal and hippocampal samples was carried out by MS-Omics using a Thermo Scientific Vanquish LC coupled to Thermo Q Exactive HF MS. An electrospray ionization interface was used as an ionization source. Analysis was performed in negative and positive ionization mode. The UPLC was performed using a slightly modified version of the protocol described by (Doneanu et al., 2011). Peak areas were extracted using Compound Discoverer 3.1 (Thermo Scientific). Identification of compounds was performed at four levels; Level 1: identification by retention times (compared against in-house authentic standards), accurate mass (with an acceptable deviation of 3 ppm), and MS/MS spectra, Level 2a: identification by retention times (compared against in-house authentic standards), accurate mass (with an acceptable deviation of 3 ppm). Level 2b: identification by accurate mass (with an acceptable deviation of 3 ppm), and MS/MS spectra, Level 3: identification by accurate mass alone (with an acceptable deviation of 3 ppm).

*Bioinformatic analysis*: Differential expression analyses were limited to features (metabolites) annotated at the highest confidence levels 1 and 2a (caecal features, n=209; hippocampal features, n=149). All analyses were performed in R (version 4.1.1). Raw feature peak area values below their associated limit of detection (as reported by MsOmics) were considered missing (i.e. set to “*NA*”), and only features with a maximum of 25% missingness per condition were retained for quantification (caecal features remaining, n=208; hippocampal features remaining, n=147). Additionally, to remove features displaying high technical variance, only metabolites with <10% relative standard deviation in the pooled quality control samples were retained (caecal features remaining, n=184; hippocampal features remaining, n=123). Data were subsequently normalized with variance stabilizing normalization (VSN), using the “*vsn*” package (allowing for 10% outliers, by default). While originally developed for microarray data (Huber et al., 2002), VSN has successfully been applied in untargeted (Li et al., 2016) and simulated (Jauhiainen et al., 2014) metabolomics of comparable dataset size to the one herein, capitalizing on similar mean-variance relationships (van den Berg et al., 2006) and error models (Rocke and Durbin, 2001). The generalized log2 (glog2) transformation employed in VSN approximates the standard log2 function for values >>0. The “*limma*” package was used for differential expression analysis, with trend = TRUE and robust = TRUE in the eBayes function (Ritchie et al., 2015). Resulting feature *p*-values were adjusted for multiple comparisons with the Benjamini-Hochberg method, with a 5% false discovery rate (FDR) threshold for significance. Partial Least Squares Discriminant Analysis (PLS-DA) was conducted to asses group clustering in feature space, with permutation testing (n=1,000) to assess model significance (defined as pQ^2^ < 0.05). PLS-DA was performed using the “*mixOmics*” and “*ropls*” packages. The full list of caecal and hippocampal metabolites quantified is provided in Supplementary Table 4.

**Supplementary Table 4:** Quantified metabolites in caecal content and hippocampus.

|  |  | Caecum |  |  | Hippocampus |  |  |
| --- | --- | --- | --- | --- | --- | --- | --- |
| Metabolite | Annotation level | FC | P-value | Adjusted P-value | FC | P-value | Adjusted P-value |
| Histidine | 1 | 0.456 | 0.00431 | 0.477 | 0.0259 | 0.771 | 0.989 |
| 2-Phenylethyl acetate | 1 | 0.429 | 0.00787 | 0.477 | - | - | - |
| Aminoadipic acid | 2a | 0.272 | 0.0117 | 0.477 | 0.146 | 0.151 | 0.989 |
| Cytosine | 1 | -0.699 | 0.0118 | 0.477 | - | - | - |
| Succinic acid | 1 | 0.337 | 0.013 | 0.477 | 0.672 | 0.309 | 0.989 |
| Tetradecanedioic acid | 1 | 0.413 | 0.0186 | 0.57 | - | - | - |
| Xanthurenic acid | 1 | 0.553 | 0.0228 | 0.576 | - | - | - |
| Hydroxybutyric acid | 1 | -0.534 | 0.0255 | 0.576 | - | - | - |
| 2-Oxo-3-phenylpropanoic acid | 1 | -0.667 | 0.0314 | 0.576 | - | - | - |
| Acetylhistidine | 1 | 0.389 | 0.0363 | 0.576 | - | - | - |
| Vanillyl alcohol | 2a | 0.247 | 0.0394 | 0.576 | - | - | - |
| Kynurenic acid | 1 | 0.465 | 0.0396 | 0.576 | - | - | - |
| Galactonic acid | 2a | -0.599 | 0.0407 | 0.576 | - | - | - |
| Ferulic acid | 1 | -0.491 | 0.0504 | 0.649 | - | - | - |
| 4-Coumaric acid | 1 | -0.455 | 0.054 | 0.649 | - | - | - |
| Hexadecanedioic acid | 2a | 0.425 | 0.0565 | 0.649 | - | - | - |
| 5-Aminolevulinic acid | 1 | -0.299 | 0.0609 | 0.659 | - | - | - |
| Genistein | 2a | -0.848 | 0.0698 | 0.702 | - | - | - |
| Pipecolinic acid | 1 | 0.366 | 0.0725 | 0.702 | - | - | - |
| 3-Methyl-2-oxovaleric acid | 2a | -0.386 | 0.0822 | 0.74 | - | - | - |
| Glucose-x-Phosphate | 2a | -0.405 | 0.0866 | 0.74 | -0.605 | 0.0576 | 0.989 |
| N-Acetylornithine | 1 | 0.496 | 0.0892 | 0.74 | -0.0884 | 0.469 | 0.989 |
| N-Acetylglutamic acid | 1 | 0.238 | 0.101 | 0.74 | 0.0677 | 0.577 | 0.989 |
| Indole-3-lactic acid | 2a | -0.208 | 0.099 | 0.74 | - | - | - |
| Muricholic-acid | 1 | 0.709 | 0.104 | 0.74 | - | - | - |
| Deoxycholic acid | 1 | 0.52 | 0.105 | 0.74 | - | - | - |
| N-Acetylglutamine | 1 | 0.228 | 0.119 | 0.742 | 0.233 | 0.165 | 0.989 |
| Citric acid | 1 | -0.3 | 0.121 | 0.742 | - | - | - |
| Pentose alcohol2 | 2a | -0.238 | 0.125 | 0.742 | - | - | - |
| 7-Methylguanine | 1 | -0.256 | 0.13 | 0.742 | -0.0513 | 0.744 | 0.989 |
| Raffinose | 1 | -0.57 | 0.132 | 0.742 | - | - | - |
| Xanthine | 1 | -0.179 | 0.134 | 0.742 | -0.105 | 0.811 | 0.989 |
| Pinitol | 2a | -0.307 | 0.14 | 0.742 | - | - | - |
| Dodecanedioic acid | 1 | 0.185 | 0.14 | 0.742 | - | - | - |
| Asparagine | 1 | 0.412 | 0.151 | 0.742 | - | - | - |
| Ornithine | 1 | -0.256 | 0.152 | 0.742 | - | - | - |
| Isethionic acid | 2a | -0.521 | 0.152 | 0.742 | 0.11 | 0.566 | 0.989 |
| Indole-3-glyoxylic acid | 1 | -0.688 | 0.158 | 0.742 | - | - | - |
| Lysine | 2a | -0.19 | 0.159 | 0.742 | - | - | - |
| Acetylarginine | 1 | 0.283 | 0.163 | 0.742 | 0.149 | 0.376 | 0.989 |
| Serine | 1 | -0.127 | 0.173 | 0.742 | -0.119 | 0.212 | 0.989 |
| Cholic acid | 1 | 0.664 | 0.176 | 0.742 | - | - | - |
| Citrulline | 1 | -0.372 | 0.181 | 0.742 | -0.0757 | 0.418 | 0.989 |
| Xanthosin | 1 | -0.383 | 0.185 | 0.742 | -0.388 | 0.454 | 0.989 |
| α-Hydroxyisovaleric acid | 1 | 0.435 | 0.188 | 0.742 | - | - | - |
| Arginine | 1 | 0.703 | 0.189 | 0.742 | -0.0444 | 0.804 | 0.989 |
| Allopurinol | 1 | 0.449 | 0.19 | 0.742 | 0.0525 | 0.886 | 0.989 |
| 2-Deoxyinosine | 1 | 0.457 | 0.194 | 0.742 | - | - | - |
| Allantoin | 1 | -0.629 | 0.211 | 0.756 | - | - | - |
| 3-(3-Hydroxyphenyl)-propionate | 1 | 0.399 | 0.213 | 0.756 | - | - | - |
| Cystathionine | 2a | -0.467 | 0.213 | 0.756 | - | - | - |
| Acetylcholine | 1 | 0.312 | 0.22 | 0.756 | 0.0958 | 0.927 | 0.989 |
| Glycine | 2a | -0.156 | 0.227 | 0.756 | 0.0349 | 0.722 | 0.989 |
| 5-Hydroxytryptamine (Serotonin) | 2a | -0.402 | 0.227 | 0.756 | - | - | - |
| Pyridoxamine | 1 | 0.227 | 0.23 | 0.756 | - | - | - |
| Dihydroferulic acid | 1 | 0.317 | 0.231 | 0.756 | - | - | - |
| Galactos/glucos-amin | 1 | -0.236 | 0.234 | 0.756 | - | - | - |
| Glycylvaline | 1 | -0.449 | 0.243 | 0.756 | - | - | - |
| Ursodeoxycholic acid | 2a | 0.28 | 0.245 | 0.756 | - | - | - |
| Thymine | 1 | -0.362 | 0.25 | 0.756 | - | - | - |
| Glyceric acid | 1 | -0.14 | 0.251 | 0.756 | 0.243 | 0.234 | 0.989 |
| Pyruvic acid | 1 | -0.246 | 0.264 | 0.783 | -0.411 | 0.704 | 0.989 |
| N-α-Acetyllysine | 1 | 0.184 | 0.274 | 0.794 | - | - | - |
| Glucuronic acid | 1 | -0.457 | 0.276 | 0.794 | - | - | - |
| Threonic acid | 2a | 0.279 | 0.281 | 0.796 | -0.029 | 0.93 | 0.989 |
| 2-Hydroxyphenylacetic acid | 1 | 0.233 | 0.299 | 0.812 | - | - | - |
| Noradrenaline | 1 | 0.179 | 0.301 | 0.812 | - | - | - |
| Uridine | 1 | -0.458 | 0.302 | 0.812 | -0.0386 | 0.756 | 0.989 |
| 4-Hydroxybenzaldehyde | 1 | -0.29 | 0.305 | 0.812 | -0.144 | 0.763 | 0.989 |
| Hydroxyisocaproic acid | 1 | 0.308 | 0.329 | 0.865 | - | - | - |
| Inosine | 2a | -0.323 | 0.347 | 0.866 | - | - | - |
| Uracil | 1 | -0.094 | 0.349 | 0.866 | 0.275 | 0.332 | 0.989 |
| 3-Phenyllactic acid | 1 | 0.162 | 0.352 | 0.866 | - | - | - |
| N-Acetyltyrosine | 2a | 0.137 | 0.356 | 0.866 | -0.0137 | 0.978 | 0.989 |
| Ascorbic acid | 1 | 0.311 | 0.357 | 0.866 | -1.14 | 0.427 | 0.989 |
| Urocanic acid | 1 | -0.287 | 0.358 | 0.866 | -0.0173 | 0.969 | 0.989 |
| Shikimic acid | 1 | 0.243 | 0.363 | 0.866 | - | - | - |
| Hexose dimer1 | 2a | -0.149 | 0.367 | 0.866 | - | - | - |
| Pentose alcohol1 | 1 | -0.123 | 0.386 | 0.867 | - | - | - |
| Propionylcarnitine (C3) | 1 | 0.512 | 0.388 | 0.867 | -0.119 | 0.549 | 0.989 |
| N-Acetylneuraminic acid | 1 | 0.398 | 0.388 | 0.867 | 0.0375 | 0.794 | 0.989 |
| 2-Methylbutyrylglycine | 1 | 0.149 | 0.389 | 0.867 | - | - | - |
| Tryptamine | 2a | -0.246 | 0.399 | 0.867 | - | - | - |
| Uric acid | 1 | -0.395 | 0.404 | 0.867 | - | - | - |
| Betaine | 1 | 0.24 | 0.407 | 0.867 | - | - | - |
| 5-Methyluridine | 2a | -0.199 | 0.416 | 0.867 | - | - | - |
| Hyodeoxycholic acid | 1 | 0.423 | 0.417 | 0.867 | - | - | - |
| Methanesulfonic acid | 1 | -0.217 | 0.419 | 0.867 | - | - | - |
| Thiamine (B1) | 1 | 0.104 | 0.424 | 0.867 | - | - | - |
| Citraconic acid | 1 | 0.121 | 0.426 | 0.867 | - | - | - |
| Hexose1 | 1 | -0.256 | 0.436 | 0.867 | -0.0872 | 0.312 | 0.989 |
| Traumatic Acid | 1 | 0.129 | 0.44 | 0.867 | - | - | - |
| L-DOPA | 2a | -0.19 | 0.442 | 0.867 | - | - | - |
| Aldopentose2 | 2a | 0.0913 | 0.443 | 0.867 | - | - | - |
| N-Acetylmannosamine | 1 | 0.0961 | 0.471 | 0.913 | - | - | - |
| Hexanoylglycine | 1 | 0.129 | 0.484 | 0.919 | - | - | - |
| α-Ketoglutaric acid | 1 | -0.196 | 0.488 | 0.919 | - | - | - |
| Serine phosphate | 2a | 0.101 | 0.496 | 0.919 | 0.179 | 0.281 | 0.989 |
| Carnosine | 1 | -0.16 | 0.499 | 0.919 | -0.158 | 0.352 | 0.989 |
| Azelaic acid | 1 | -0.0783 | 0.504 | 0.919 | - | - | - |
| Pantothenic acid | 1 | 0.118 | 0.504 | 0.919 | - | - | - |
| N-Formylmethionine | 1 | 0.0922 | 0.513 | 0.919 | - | - | - |
| 4-Guanidinobutyric acid | 2a | 0.111 | 0.514 | 0.919 | - | - | - |
| Tryptophan | 1 | -0.134 | 0.528 | 0.933 | 0.0558 | 0.557 | 0.989 |
| 4-Hydroxy-2,5-dimethyl-3(2H)-furanone (Furaneol) | 2a | -0.179 | 0.565 | 0.954 | - | - | - |
| Acetylsalicylic acid | 2a | 0.12 | 0.568 | 0.954 | - | - | - |
| 3-Methyladipic acid | 1 | 0.153 | 0.569 | 0.954 | - | - | - |
| Homoarginine | 2a | -0.106 | 0.574 | 0.954 | -0.0136 | 0.916 | 0.989 |
| Thymidine | 1 | 0.164 | 0.582 | 0.954 | - | - | - |
| 3-Methylbutanoic acid | 2a | 0.0826 | 0.588 | 0.954 | - | - | - |
| N-Acetylphenylalanine | 1 | 0.161 | 0.595 | 0.954 | - | - | - |
| 5-Methylcytosine | 1 | -0.119 | 0.617 | 0.954 | - | - | - |
| Choline | 1 | -0.143 | 0.618 | 0.954 | -0.0821 | 0.622 | 0.989 |
| Aldopentose1 | 2a | -0.0529 | 0.621 | 0.954 | - | - | - |
| δ-Gluconolactone | 1 | -0.209 | 0.625 | 0.954 | - | - | - |
| Glutamic acid | 1 | 0.0373 | 0.627 | 0.954 | -0.00723 | 0.885 | 0.989 |
| Creatine | 1 | 0.27 | 0.627 | 0.954 | -0.0351 | 0.411 | 0.989 |
| Benzaldehyde | 2a | 0.0531 | 0.638 | 0.954 | - | - | - |
| Valine | 1 | 0.0489 | 0.652 | 0.954 | - | - | - |
| Alanine | 1 | -0.0341 | 0.655 | 0.954 | -0.0244 | 0.948 | 0.989 |
| Hippuric acid | 1 | 0.279 | 0.661 | 0.954 | - | - | - |
| α-Aminobutyric acid | 1 | 0.034 | 0.666 | 0.954 | - | - | - |
| Sorbitol/Manitol | 2a | -0.126 | 0.669 | 0.954 | - | - | - |
| Indole-3-carboxylate | 1 | -0.149 | 0.685 | 0.954 | - | - | - |
| Leucylalanine | 1 | -0.0895 | 0.688 | 0.954 | - | - | - |
| Homocitrulline | 2a | -0.0545 | 0.69 | 0.954 | -0.344 | 0.0411 | 0.989 |
| Isocitric acid | 1 | -0.0697 | 0.69 | 0.954 | 0.359 | 0.301 | 0.989 |
| Phenylacetylglutamine | 2a | 0.0636 | 0.692 | 0.954 | - | - | - |
| 3-Methylxanthine | 1 | 0.148 | 0.693 | 0.954 | - | - | - |
| Kynuerine | 2a | 0.0581 | 0.697 | 0.954 | - | - | - |
| Phosphoenolpyruvate | 2a | -0.0241 | 0.698 | 0.954 | - | - | - |
| 5-Methoxytryptophan | 1 | 0.153 | 0.706 | 0.954 | - | - | - |
| Proline | 2a | 0.0384 | 0.709 | 0.954 | 0.0451 | 0.761 | 0.989 |
| 3-Methoxytyramine | 1 | -0.0214 | 0.981 | 0.989 | - | - | - |
| Carnitine | 1 | 0.146 | 0.712 | 0.954 | -0.00918 | 0.887 | 0.989 |
| Leucine | 1 | 0.0362 | 0.716 | 0.954 | -0.0158 | 0.885 | 0.989 |
| N-Acetyltryptophan | 2a | 0.0512 | 0.718 | 0.954 | - | - | - |
| Pyroglutamic acid | 1 | 0.0575 | 0.719 | 0.954 | 0.182 | 0.611 | 0.989 |
| Threonine | 1 | 0.0302 | 0.722 | 0.954 | 0.05 | 0.493 | 0.989 |
| N-Acetylalanine | 1 | -0.0404 | 0.723 | 0.954 | - | - | - |
| Trigonelline (N'-methylnicotinate) | 1 | 0.0887 | 0.729 | 0.954 | - | - | - |
| 3-Methylhistidine | 1 | 0.0916 | 0.731 | 0.954 | -0.442 | 0.172 | 0.989 |
| Methionine sulfoxide | 2a | -0.0408 | 0.739 | 0.954 | -0.302 | 0.403 | 0.989 |
| Methionine | 1 | -0.0288 | 0.744 | 0.954 | -0.0137 | 0.916 | 0.989 |
| Suberic acid | 1 | 0.0596 | 0.754 | 0.954 | - | - | - |
| Daidzein | 1 | -0.0867 | 0.758 | 0.954 | - | - | - |
| Malic acid | 1 | -0.096 | 0.761 | 0.954 | 0.239 | 0.598 | 0.989 |
| 4-Vinylguaicol | 1 | 0.0578 | 0.762 | 0.954 | - | - | - |
| Sebacic acid | 1 | 0.0335 | 0.77 | 0.957 | - | - | - |
| 3-(2-Hydroxyethyl)indole | 2a | 0.0407 | 0.796 | 0.977 | - | - | - |
| Vancomycin | 1 | -0.0251 | 0.801 | 0.977 | - | - | - |
| Dopamine | 2a | -0.0543 | 0.808 | 0.977 | - | - | - |
| Nicotinic acid | 1 | 0.0256 | 0.81 | 0.977 | - | - | - |
| 3-Ureidopropionate | 2a | -0.0835 | 0.818 | 0.977 | -0.117 | 0.331 | 0.989 |
| Diaminopimelic acid | 2a | 0.0583 | 0.822 | 0.977 | 0.107 | 0.492 | 0.989 |
| 4-Methyl-2-oxovaleric acid (ketoleucine) | 2a | -0.0375 | 0.898 | 0.987 | - | - | - |
| N-Acetylserine | 1 | -0.0849 | 0.826 | 0.977 | - | - | - |
| 2-Ethylhydracrylic acid | 2a | 0.0331 | 0.828 | 0.977 | -0.362 | 0.215 | 0.989 |
| IMP | 2a | 0.024 | 0.843 | 0.987 | - | - | - |
| N8-Acetylspermidine | 1 | -0.0303 | 0.853 | 0.987 | - | - | - |
| 2'Deoxyuridine | 1 | 0.0507 | 0.866 | 0.987 | - | - | - |
| Isoleucine | 1 | 0.0197 | 0.868 | 0.987 | 0.147 | 0.466 | 0.989 |
| Aspartic acid | 1 | 0.0197 | 0.872 | 0.987 | 0.041 | 0.908 | 0.989 |
| Lactic acid | 1 | 0.0609 | 0.884 | 0.987 | -0.0122 | 0.702 | 0.989 |
| Hexose dimer2 | 1 | 0.0378 | 0.885 | 0.987 | - | - | - |
| Gentisic acid | 1 | -0.0385 | 0.895 | 0.987 | - | - | - |
| Ethylmalonic acid | 1 | 0.0209 | 0.899 | 0.987 | - | - | - |
| Salicylic acid | 1 | 0.0207 | 0.901 | 0.987 | - | - | - |
| Cystine | 2a | 0.0138 | 0.902 | 0.987 | -0.0579 | 0.874 | 0.989 |
| Taurine | 1 | 0.046 | 0.91 | 0.987 | -0.117 | 0.034 | 0.989 |
| Glycylleucine | 1 | 0.0229 | 0.912 | 0.987 | - | - | - |
| Hypoxanthine | 1 | 0.0143 | 0.917 | 0.987 | -0.147 | 0.66 | 0.989 |
| Argininosuccinic acid | 2a | 0.0138 | 0.942 | 0.989 | -0.213 | 0.164 | 0.989 |
| Pimelic acid | 1 | -0.0114 | 0.944 | 0.989 | - | - | - |
| Phenylalanine | 1 | 0.00662 | 0.95 | 0.989 | 0.00452 | 0.963 | 0.989 |
| Theanine | 1 | 0.0118 | 0.952 | 0.989 | - | - | - |
| Prostaglandine E2 | 2a | -0.0108 | 0.959 | 0.989 | - | - | - |
| Adipic acid | 1 | 0.0107 | 0.96 | 0.989 | - | - | - |
| Quinic acid | 1 | -0.0215 | 0.966 | 0.989 | - | - | - |
| N-Acetylaspartic acid | 1 | -0.0113 | 0.969 | 0.989 | 0.107 | 0.685 | 0.989 |
| Tyrosine | 1 | 0.00353 | 0.973 | 0.989 | 0.0854 | 0.385 | 0.989 |
| Glycolic acid | 1 | -<br>0.00419 | 0.984 | 0.989 | - | - | - |
| 5-Hydroxyindol-3-acetic acid | 1 | 0.00997 | 0.989 | 0.989 | - | - | - |
| Acetylmuramic acid | 2a | -<br>0.00267 | 0.989 | 0.989 | - | - | - |
| 4-Guanidinobutyric acid | 1 | - | - | - | -0.244 | 0.0339 | 0.989 |
| 2-Deoxycytidine | 1 | - | - | - | -0.249 | 0.0538 | 0.989 |
| Glyceraldehyde-3-phosphate | 2a | - | - | - | -0.424 | 0.0639 | 0.989 |
| Morphine 6-β-glucopyranosiduronide | 2a | - | - | - | -0.392 | 0.114 | 0.989 |
| O-Phosphorylethanolamine | 1 | - | - | - | -0.173 | 0.122 | 0.989 |
| Allantoin | 2a | - | - | - | -0.12 | 0.129 | 0.989 |
| Pentose alcohol | 2a | - | - | - | -0.164 | 0.129 | 0.989 |
| Ribulose-5-phosphate | 1 | - | - | - | -0.272 | 0.154 | 0.989 |
| 3-Methyladipic acid | 2a | - | - | - | -0.328 | 0.174 | 0.989 |
| Ciproflaxacin | 2a | - | - | - | -0.383 | 0.179 | 0.989 |
| N-α-Acetyllysine | 2a | - | - | - | 0.216 | 0.185 | 0.989 |
| 5-Hydroxytryptamine (Serotonin) | 1 | - | - | - | 0.368 | 0.214 | 0.989 |
| Fumaric acid | 1 | - | - | - | 0.135 | 0.233 | 0.989 |
| 2-Deoxyuridine | 2a | - | - | - | -0.0898 | 0.234 | 0.989 |
| Dulcitol | 2a | - | - | - | -0.105 | 0.24 | 0.989 |
| Guanosine | 1 | - | - | - | 1.23 | 0.244 | 0.989 |
| N-Iso-Valerylglycine | 2a | - | - | - | -0.358 | 0.282 | 0.989 |
| Pyridoxal 5'-phosphate | 2a | - | - | - | 0.21 | 0.313 | 0.989 |
| Adipic acid | 2a | - | - | - | -0.218 | 0.317 | 0.989 |
| GMP | 1 | - | - | - | 0.975 | 0.329 | 0.989 |
| Nicotine amide | 1 | - | - | - | -0.0478 | 0.336 | 0.989 |
| Quinic acid | 2a | - | - | - | 0.406 | 0.357 | 0.989 |
| Fructose | 2a | - | - | - | -0.0888 | 0.401 | 0.989 |
| UMP | 2a | - | - | - | 1.14 | 0.422 | 0.989 |
| Asparagine | 2a | - | - | - | -0.0944 | 0.427 | 0.989 |
| CMP | 1 | - | - | - | 0.278 | 0.432 | 0.989 |
| Glutathione | 1 | - | - | - | -0.419 | 0.446 | 0.989 |
| 2-Hydroxybutyric acid | 2a | - | - | - | 0.0779 | 0.475 | 0.989 |
| AMP | 1 | - | - | - | 1.29 | 0.53 | 0.989 |
| Ethylmalonic acid/Glutaric acid | 2a | - | - | - | -0.115 | 0.531 | 0.989 |
| α-aminobutyric acid | 1 | - | - | - | 0.0382 | 0.547 | 0.989 |
| cis-Aconitic acid | 2a | - | - | - | 0.246 | 0.549 | 0.989 |
| α-Ketoglutaric acid | 2a | - | - | - | 0.361 | 0.562 | 0.989 |
| Glyoxylic acid | 2a | - | - | - | 0.252 | 0.58 | 0.989 |
| Uric acid | 2a | - | - | - | 0.0983 | 0.58 | 0.989 |
| Cysteine | 2a | - | - | - | 0.432 | 0.683 | 0.989 |
| Ornithine | 2a | - | - | - | -0.0499 | 0.61 | 0.989 |
| Galactonic acid | 1 | - | - | - | 0.153 | 0.632 | 0.989 |
| CDP-choline | 1 | - | - | - | -0.0798 | 0.639 | 0.989 |
| 2-Deoxyinosine | 2a | - | - | - | 0.149 | 0.751 | 0.989 |
| Hexose2 | 2a | - | - | - | 0.395 | 0.685 | 0.989 |
| Choline phosphate (PCHO) | 1 | - | - | - | -0.157 | 0.77 | 0.989 |
| Lysine | 1 | - | - | - | -0.0368 | 0.807 | 0.989 |
| Butyrylcarnitine (C4) | 1 | - | - | - | -0.0438 | 0.81 | 0.989 |
| γ-Aminobutyric acid (GABA) | 1 | - | - | - | -0.0329 | 0.821 | 0.989 |
| Anserine | 1 | - | - | - | -0.0299 | 0.835 | 0.989 |
| Pentose2 | 2a | - | - | - | -0.232 | 0.837 | 0.989 |
| Adenosine | 1 | - | - | - | -0.044 | 0.851 | 0.989 |
| Pentose1 | 2a | - | - | - | 0.055 | 0.862 | 0.989 |
| Adenine | 1 | - | - | - | 0.0217 | 0.866 | 0.989 |
| CDP-ethanolamine | 2a | - | - | - | -0.0186 | 0.87 | 0.989 |
| Phosphoenolpyruvate | 1 | - | - | - | 0.126 | 0.874 | 0.989 |
| Acetylcarnitine (C2) | 1 | - | - | - | -0.0152 | 0.886 | 0.989 |
| Methanesulfonic acid | 2a | - | - | - | -0.0185 | 0.886 | 0.989 |
| Glutamine | 1 | - | - | - | -<br>0.00793 | 0.917 | 0.989 |
| Creatinine | 1 | - | - | - | -<br>0.00639 | 0.922 | 0.989 |
| Inosine | 1 | - | - | - | 0.0316 | 0.922 | 0.989 |
| Cytidine | 1 | - | - | - | -0.0172 | 0.937 | 0.989 |
| Trimethylamine N-Oxide | 2a | - | - | - | -<br>0.00634 | 0.967 | 0.989 |
| Hexomedimer1 | 2a | - | - | - | 0.00627 | 0.981 | 0.989 |
| N1-Methyl-Adenosine | 1 | - | - | - | -0.0013 | 0.99 | 0.99 |
| (-) Not quantified<br>Abbreviation: FC – Fold change (generalised logarithm, base 2)<br>NB: the same metabolite can be annotated at different confidence levels in the two datasets. |  |  |  |  |  |  |  |

### Quantitative Real-time PCR for immune profile in rat colon

Isolation of total RNA was performed using the GenElute kit (Sigma) as described by the manufacturer. Briefly, rat colon was homogenized in lysis buffer using 1 mm glass beads (Thistle Scientific) in a MagnaLyser system (Roche). The samples were filtered through a binding column, and an equal volume of 70% ethanol was added to the filtrate. After purification through columns and washing, mRNA was eluted into elution solution. Total RNA yield and purity were assessed with the Nanodrop System (Thermo Scientific) and isolated RNA was stored in RNase-free water at −80 °C. Quantitative real-time reverse transcription PCR (qRT-PCR) was carried out with the QuantiNova SYBR Green RT-PCR kit (Qiagen, Manchester USA) using a LightCycler® 480 Instrument II (Roche) according to the manufacturer’s instruction. Each reaction consisted of 1μl of sample, 5 μl of 2x Quanti Nova SYBR Green RT-PCR Master Mix, 0.1 μl of QN SYBR Green RT-Mix, 0.5 μl (0.5 μM) of both forward (F) and reverse (R) primers (detailed in Supplementary Table 5) and 3.4 μl of RNase free water. Relative quantification was based on the internal reference gene β-Actin to determine virtual mRNA levels of the target gene. Reactions were run in technical triplicates for each biological sample, and mean ct values for each reaction were taken into account for data analysis. A melting curve was obtained for the amplicon products to determine their melting temperatures to ascertain primer specificity.

**Supplementary Table 5:**
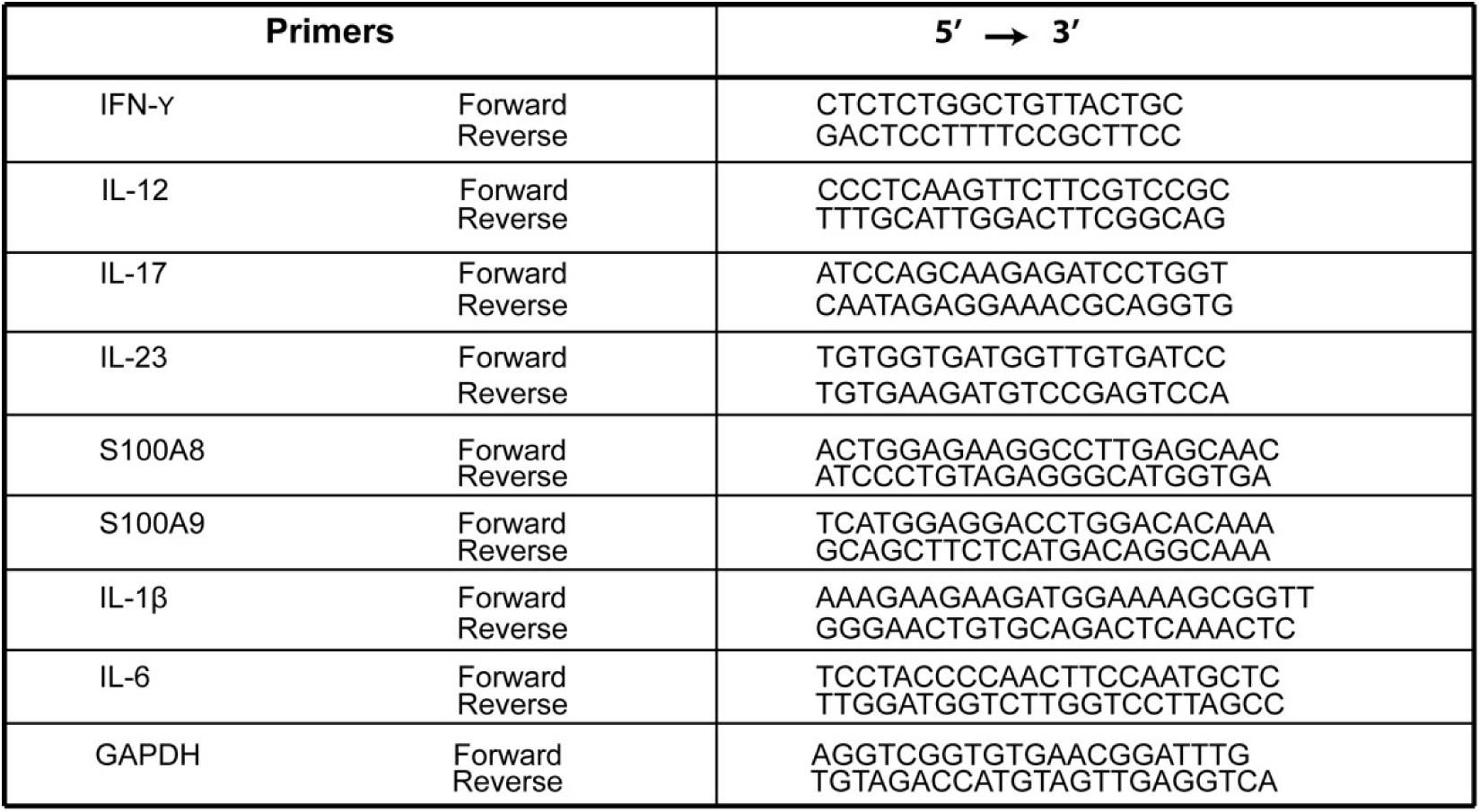
Primers used for RT-qPCR on rat colon

### Cytokine assay in rat plasma

Tail blood (10 days after FMT) and trunk blood (at the end of the study) was collected in tubes containing heparin and immediately centrifuged at 1,000 x g for 15 min at 4°C. The resulting supernatant (plasma) was collected, aliquoted, and stored at −80°C. Plasma levels of IL-6, IL-4, IL-10 IFN-γ, TNF-α, Il-1β were assayed in duplicate using high sensitivity commercially available electrochemiluminescence MULTI-SPOT^®^ Meso Scale Discovery kits (MSD, Rockville, MD, 75USA) as per manufacturer’s instructions. Reading and analysis were conducted using the MESO QuickPlex SQ 120, Sector Imager 2400. Concentration is expressed in pg/ml.

### Statistics

Data are depicted as mean with the standard error of the mean (SEM). Normal distribution was determined by the Shapiro-Wilk test. For normally distributed data, single comparisons were tested using independent Student’s t-test. Non-parametric data were examined using the Mann-Whitney U-Test. Experiments with more than two groups were subjected to a one-way ANOVA if normally distributed and compared using Bonferroni’s *post hoc* test. Multiple comparisons of nonparametric data were made by a Kruskal-Wallis analysis followed by Dunn’s *post hoc* test. Experiments with two between factors (treatment, donors) were analyzed using a two-way ANOVA to determine genotype and treatment effect or interaction between both factors. Experiments with one “between-subjects” factor (treatment group) and one “within-subjects” factor (trial) were analyzed using a two-way mixed ANOVA. Pearson or Spearman’s rank correlation were performed when data were normally or not normally distributed, respectively. Statistical analysis was performed using GraphPad Prism 8.0 or in R (v.3.6.3) as described above. Statistical significance was set at p ≤ 0.05. Statistically significant differences are indicated in the figures by *p ≤ 0.05, **p ≤ 0.01, and ***p ≤ 0.001.

## Notes

### Competing Interest Statement

The authors have declared no competing interest.

## REFERENCES

2016 Alzheimer’s disease facts and figures., 2016.. Alzheimers. Dement. 12, 459–509. https://doi.org/10.1016/j.jalz.2016.03.001

Adewuyi, E.O., O’Brien, E.K., Nyholt, D.R., Porter, T., Laws, S.M., 2022. A large-scale genome-wide cross-trait analysis reveals shared genetic architecture between Alzheimer’s disease and gastrointestinal tract disorders. Commun. Biol. 5, 691. https://doi.org/10.1038/s42003-022-03607-2

Ally, B.A., Hussey, E.P., Ko, P.C., Molitor, R.J., 2013. Pattern separation and pattern completion in Alzheimer’s disease: evidence of rapid forgetting in amnestic mild cognitive impairment. Hippocampus 23, 1246–1258. https://doi.org/10.1002/hipo.22162

Asti, A., Gioglio, L., 2014. Can a bacterial endotoxin be a key factor in the kinetics of amyloid fibril formation? J. Alzheimers. Dis. 39, 169–79. https://doi.org/10.3233/JAD-131394

Bacher, M., Deuster, O., Aljabari, B., Egensperger, R., Neff, F., Jessen, F., Popp, J., Noelker, C., Reese, J.P., Al-Abed, Y., Dodel, R., 2010. The role of macrophage migration inhibitory factor in Alzheimer’s disease. Mol. Med. 16, 116–121. https://doi.org/10.2119/molmed.2009.00123

Bekinschtein, P., Kent, B.A., Oomen, C.A., Clemenson, G.D., Gage, F.H., Saksida, L.M., Bussey, T.J., 2013. BDNF in the dentate gyrus is required for consolidation of “pattern-separated” memories. Cell Rep. 5, 759–768. https://doi.org/10.1016/j.celrep.2013.09.027

Bellenguez, C., Küçükali, F., Jansen, I.E., Kleineidam, L., Moreno-Grau, S., Amin, N., Naj, A.C., Campos-Martin, R., Grenier-Boley, B., Andrade, V., Holmans, P.A., Boland, A., Damotte, V., van der Lee, S.J., Costa, M.R., Kuulasmaa, T., Yang, Q., de Rojas, I., Bis, J.C., Yaqub, A., Prokic, I., Chapuis, J., Ahmad, S., Giedraitis, V., Aarsland, D., Garcia-Gonzalez, P., Abdelnour, C., Alarcón-Martín, E., Alcolea, D., Alegret, M., Alvarez, I., Álvarez, V., Armstrong, N.J., Tsolaki, A., Antúnez, C., Appollonio, I., Arcaro, M., Archetti, S., Pastor, A.A., Arosio, B., Athanasiu, L., Bailly, H., Banaj, N., Baquero, M., Barral, S., Beiser, A., Pastor, A.B., Below, J.E., Benchek, P., Benussi, L., Berr, C., Besse, C., Bessi, V., Binetti, G., Bizarro, A., Blesa, R., Boada, M., Boerwinkle, E., Borroni, B., Boschi, S., Bossù, P., Bråthen, G., Bressler, J., Bresner, C., Brodaty, H., Brookes, K.J., Brusco, L.I., Buiza-Rueda, D., Bûrger, K., Burholt, V., Bush, W.S., Calero, M., Cantwell, L.B., Chene, G., Chung, J., Cuccaro, M.L., Carracedo, Á., Cecchetti, R., Cervera-Carles, L., Charbonnier, C., Chen, H.-H., Chillotti, C., Ciccone, S., Claassen, J.A.H.R., Clark, C., Conti, E., Corma-Gómez, A., Costantini, E., Custodero, C., Daian, D., Dalmasso, M.C., Daniele, A., Dardiotis, E., Dartigues, J.-F., de Deyn, P.P., de Paiva Lopes, K., de Witte, L.D., Debette, S., Deckert, J., Del Ser, T., Denning, N., DeStefano, A., Dichgans, M., Diehl-Schmid, J., Diez-Fairen, M., Rossi, P.D., Djurovic, S., Duron, E., Düzel, E., Dufouil, C., Eiriksdottir, G., Engelborghs, S., Escott-Price, V., Espinosa, A., Ewers, M., Faber, K.M., Fabrizio, T., Nielsen, S.F., Fardo, D.W., Farotti, L., Fenoglio, C., Fernández-Fuertes, M., Ferrari, R., Ferreira, C.B., Ferri, E., Fin, B., Fischer, P., Fladby, T., Fließbach, K., Fongang, B., Fornage, M., Fortea, J., Foroud, T.M., Fostinelli, S., Fox, N.C., Franco-Macías, E., Bullido, M.J., Frank-García, A., Froelich, L., Fulton-Howard, B., Galimberti, D., García-Alberca, J.M., García-González, P., Garcia-Madrona, S., Garcia-Ribas, G., Ghidoni, R., Giegling, I., Giorgio, G., Goate, A.M., Goldhardt, O., Gomez-Fonseca, D., González-Pérez, A., Graff, C., Grande, G., Green, E., Grimmer, T., Grünblatt, E., Grunin, M., Gudnason, V., Guetta-Baranes, T., Haapasalo, A., Hadjigeorgiou, G., Haines, J.L., Hamilton-Nelson, K.L., Hampel, H., Hanon, O., Hardy, J., Hartmann, A.M., Hausner, L., Harwood, J., Heilmann-Heimbach, S., Helisalmi, S., Heneka, M.T., Hernández, I., Herrmann, M.J., Hoffmann, P., Holmes, C., Holstege, H., Vilas, R.H., Hulsman, M., Humphrey, J., Biessels, G.J., Jian, X., Johansson, C., Jun, G.R., Kastumata, Y., Kauwe, J., Kehoe, P.G., Kilander, L., Ståhlbom, A.K., Kivipelto, M., Koivisto, A., Kornhuber, J., Kosmidis, M.H., Kukull, W.A., Kuksa, P.P., Kunkle, B.W., Kuzma, A.B., Lage, C., Laukka, E.J., Launer, L., Lauria, A., Lee, C.-Y., Lehtisalo, J., Lerch, O., Lleó, A., Longstreth, W.J., Lopez, O., de Munain, A.L., Love, S., Löwemark, M., Luckcuck, L., Lunetta, K.L., Ma, Y., Macías, J., MacLeod, C.A., Maier, W., Mangialasche, F., Spallazzi, M., Marquié, M., Marshall, R., Martin, E.R., Montes, A.M., Rodríguez, C.M., Masullo, C., Mayeux, R., Mead, S., Mecocci, P., Medina, M., Meggy, A., Mehrabian, S., Mendoza, S., Menéndez-González, M., Mir, P., Moebus, S., Mol, M., Molina-Porcel, L., Montrreal, L., Morelli, L., Moreno, F., Morgan, K., Mosley, T., Nöthen, M.M., Muchnik, C., Mukherjee, S., Nacmias, B., Ngandu, T., Nicolas, G., Nordestgaard, B.G., Olaso, R., Orellana, A., Orsini, M., Ortega, G., Padovani, A., Paolo, C., Papenberg, G., Parnetti, L., Pasquier, F., Pastor, P., Peloso, G., Pérez-Cordón, A., Pérez-Tur, J., Pericard, P., Peters, O., Pijnenburg, Y.A.L., Pineda, J.A., Piñol-Ripoll, G., Pisanu, C., Polak, T., Popp, J., Posthuma, D., Priller, J., Puerta, R., Quenez, O., Quintela, I., Thomassen, J.Q., Rábano, A., Rainero, I., Rajabli, F., Ramakers, I., Real, L.M., Reinders, M.J.T., Reitz, C., Reyes-Dumeyer, D., Ridge, P., Riedel-Heller, S., Riederer, P., Roberto, N., Rodriguez-Rodriguez, E., Rongve, A., Allende, I.R., Rosende-Roca, M., Royo, J.L., Rubino, E., Rujescu, D., Sáez, M.E., Sakka, P., Saltvedt, I., Sanabria, Á., Sánchez-Arjona, M.B., Sanchez-Garcia, F., Juan, P.S., Sánchez-Valle, R., Sando, S.B., Sarnowski, C., Satizabal, C.L., Scamosci, M., Scarmeas, N., Scarpini, E., Scheltens, P., Scherbaum, N., Scherer, M., Schmid, M., Schneider, A., Schott, J.M., Selbæk, G., Seripa, D., Serrano, M., Sha, J., Shadrin, A.A., Skrobot, O., Slifer, S., Snijders, G.J.L., Soininen, H., Solfrizzi, V., Solomon, A., Song, Y., Sorbi, S., Sotolongo-Grau, O., Spalletta, G., Spottke, A., Squassina, A., Stordal, E., Tartan, J.P., Tárraga, L., Tesí, N., Thalamuthu, A., Thomas, T., Tosto, G., Traykov, L., Tremolizzo, L., Tybjærg-Hansen, A., Uitterlinden, A., Ullgren, A., Ulstein, I., Valero, S., Valladares, O., Broeckhoven, C. Van, Vance, J., Vardarajan, B.N., van der Lugt, A., Dongen, J. Van, van Rooij, J., van Swieten, J., Vandenberghe, R., Verhey, F., Vidal, J.-S., Vogelgsang, J., Vyhnalek, M., Wagner, M., Wallon, D., Wang, L.-S., Wang, R., Weinhold, L., Wiltfang, J., Windle, G., Woods, B., Yannakoulia, M., Zare, H., Zhao, Y., Zhang, X., Zhu, C., Zulaica, M., Farrer, L.A., Psaty, B.M., Ghanbari, M., Raj, T., Sachdev, P., Mather, K., Jessen, F., Ikram, M.A., de Mendonça, A., Hort, J., Tsolaki, M., Pericak-Vance, M.A., Amouyel, P., Williams, J., Frikke-Schmidt, R., Clarimon, J., Deleuze, J.-F., Rossi, G., Seshadri, S., Andreassen, O.A., Ingelsson, M., Hiltunen, M., Sleegers, K., Schellenberg, G.D., van Duijn, C.M., Sims, R., van der Flier, W.M., Ruiz, A., Ramirez, A., Lambert, J.-C., 2022. New insights into the genetic etiology of Alzheimer’s disease and related dementias. Nat. Genet. 54, 412–436. https://doi.org/10.1038/s41588-022-01024-z

Benjamin, J.S., Pilarowski, G.O., Carosso, G.A., Zhang, L., Huso, D.L., Goff, L.A., Vernon, H.J., Hansen, K.D., Bjornsson, H.T., 2017. A ketogenic diet rescues hippocampal memory defects in a mouse model of Kabuki syndrome. Proc. Natl. Acad. Sci. U. S. A. 114, 125–130. https://doi.org/10.1073/pnas.1611431114

Berron, D., Cardenas-Blanco, A., Bittner, D., Metzger, C.D., Spottke, A., Heneka, M.T., Fliessbach, K., Schneider, A., Teipel, S.J., Wagner, M., Speck, O., Jessen, F., Düzel, E., 2019. Higher CSF Tau Levels Are Related to Hippocampal Hyperactivity and Object Mnemonic Discrimination in Older Adults. J. Neurosci. Off. J. Soc. Neurosci. 39, 8788–8797. https://doi.org/10.1523/JNEUROSCI.1279-19.2019

Bertram, L., Tanzi, R.E., 2008. Thirty years of Alzheimer’s disease genetics: the implications of systematic meta-analyses. Nat. Rev. Neurosci. 9, 768–778. https://doi.org/10.1038/nrn2494

Bettcher, B.M., Tansey, M.G., Dorothée, G., Heneka, M.T., 2021. Peripheral and central immune system crosstalk in Alzheimer disease - a research prospectus. Nat. Rev. Neurol. 17, 689–701. https://doi.org/10.1038/s41582-021-00549-x

Bjarnason, I., 2017. The Use of Fecal Calprotectin in Inflammatory Bowel Disease. Gastroenterol. Hepatol. (N. Y). 13, 53–56.

Blennow, K., de Leon, M.J., Zetterberg, H., 2006. Alzheimer’s disease. Lancet (London, England) 368, 387–403. https://doi.org/10.1016/S0140-6736(06)69113-7

Braak, H., Braak, E., Bohl, J., 1993. Staging of Alzheimer-related cortical destruction. Eur. Neurol. 33, 403–408. https://doi.org/10.1159/000116984

Brännström, K., Islam, T., Sandblad, L., Olofsson, A., 2017. The role of histidines in amyloid β fibril assembly. FEBS Lett. 591, 1167–1175. https://doi.org/10.1002/1873-3468.12616

Brosseron, F., Krauthausen, M., Kummer, M., Heneka, M.T., 2014. Body Fluid Cytokine Levels in Mild Cognitive Impairment and Alzheimer???s Disease: a Comparative Overview. Mol. Neurobiol. 50, 534–544. https://doi.org/10.1007/s12035-014-8657-1

Cattaneo, A., Cattane, N., Galluzzi, S., Provasi, S., Lopizzo, N., Festari, C., Ferrari, C., Guerra, U.P., Paghera, B., Muscio, C., Bianchetti, A., Volta, G.D., Turla, M., Cotelli, M.S., Gennuso, M., Prelle, A., Zanetti, O., Lussignoli, G., Mirabile, D., Bellandi, D., Gentile, S., Belotti, G., Villani, D., Harach, T., Bolmont, T., Padovani, A., Boccardi, M., Frisoni, G.B., 2017. Association of brain amyloidosis with pro-inflammatory gut bacterial taxa and peripheral inflammation markers in cognitively impaired elderly. Neurobiol. Aging 49, 60–68. https://doi.org/10.1016/j.neurobiolaging.2016.08.019

Chen, Yijing, Fang, L., Chen, S., Zhou, H., Fan, Y., Lin, L., Li, J., Xu, J., Chen, Yuewen, Ma, Y., Chen, Yu, 2020. Gut Microbiome Alterations Precede Cerebral Amyloidosis and Microglial Pathology in a Mouse Model of Alzheimer’s Disease. Biomed Res. Int. 2020, 8456596. https://doi.org/10.1155/2020/8456596

Choi, S.H., Tanzi, R.E., 2019. Is Alzheimer’s Disease a Neurogenesis Disorder? Cell Stem Cell 25, 7–8. https://doi.org/10.1016/j.stem.2019.06.001

Choo, J.M., Leong, L.E.X., Rogers, G.B., 2015. Sample storage conditions significantly influence faecal microbiome profiles. Sci. Rep. 5, 16350. https://doi.org/10.1038/srep16350

Clelland, C.D., Choi, M., Romberg, C., Clemenson, G.D.J., Fragniere, A., Tyers, P., Jessberger, S., Saksida, L.M., Barker, R.A., Gage, F.H., Bussey, T.J., 2009. A functional role for adult hippocampal neurogenesis in spatial pattern separation. Science 325, 210–213. https://doi.org/10.1126/science.1173215

Connell, E., Le Gall, G., Pontifex, M.G., Sami, S., Cryan, J.F., Clarke, G., Müller, M., Vauzour, D., 2022. Microbial-derived metabolites as a risk factor of age-related cognitive decline and dementia. Mol. Neurodegener. 17, 43. https://doi.org/10.1186/s13024-022-00548-6

Corso, G., Cristofano, A., Sapere, N., la Marca, G., Angiolillo, A., Vitale, M., Fratangelo, R., Lombardi, T., Porcile, C., Intrieri, M., Di Costanzo, A., 2017. Serum Amino Acid Profiles in Normal Subjects and in Patients with or at Risk of Alzheimer Dementia. Dement. Geriatr. Cogn. Dis. Extra 7, 143–159. https://doi.org/10.1159/000466688

Craig-Schapiro, R., Kuhn, M., Xiong, C., Pickering, E.H., Liu, J., Misko, T.P., Perrin, R.J., Bales, K.R., Soares, H., Fagan, A.M., Holtzman, D.M., 2011. Multiplexed immunoassay panel identifies novel CSF biomarkers for Alzheimer’s disease diagnosis and prognosis. PLoS One 6, e18850. https://doi.org/10.1371/journal.pone.0018850

Cruz-Pereira, J.S., Rea, K., Nolan, Y.M., O’Leary, O.F., Dinan, T.G., Cryan, J.F., 2020. Depression’s Unholy Trinity: Dysregulated Stress, Immunity, and the Microbiome. Annu. Rev. Psychol. 71, 49–78. https://doi.org/10.1146/annurev-psych-122216-011613

D’Angelo, F., Felley, C., Frossard, J.L., 2017. Calprotectin in Daily Practice: Where Do We Stand in 2017? Digestion 95, 293–301. https://doi.org/10.1159/000476062

de Lucia, C., Murphy, T., Maruszak, A., Wright, P., Powell, T.R., Hartopp, N., de Jong, S., O’Sullivan, M.J., Breen, G., Price, J., Lovestone, S., Thuret, S., 2021. Serum from Older Adults Increases Apoptosis and Molecular Aging Markers in Human Hippocampal Progenitor Cells. Aging Dis. 12, 2151–2172. https://doi.org/10.14336/AD.2021.0409

de Lucia, C., Murphy, T., Steves, C.J., Dobson, R.J.B., Proitsi, P., Thuret, S., 2020. Lifestyle mediates the role of nutrient-sensing pathways in cognitive aging: cellular and epidemiological evidence. Commun. Biol. 3, 157. https://doi.org/10.1038/s42003-020-0844-1

Dekkers, K.F., Sayols-Baixeras, S., Baldanzi, G., Nowak, C., Hammar, U., Nguyen, D., Varotsis, G., Brunkwall, L., Nielsen, N., Eklund, A.C., Bak Holm, J., Nielsen, H.B., Ottosson, F., Lin, Y.-T., Ahmad, S., Lind, L., Sundström, J., Engström, G., Smith, J.G., Ärnlöv, J., Orho-Melander, M., Fall, T., 2022. An online atlas of human plasma metabolite signatures of gut microbiome composition. Nat. Commun. 13, 5370. https://doi.org/10.1038/s41467-022-33050-0

Deng, W., Aimone, J.B., Gage, F.H., 2010. New neurons and new memories: how does adult hippocampal neurogenesis affect learning and memory? Nat. Rev. Neurosci. 11, 339–350. https://doi.org/10.1038/nrn2822

Drapeau, E., Mayo, W., Aurousseau, C., Le Moal, M., Piazza, P.-V., Abrous, D.N., 2003. Spatial memory performances of aged rats in the water maze predict levels of hippocampal neurogenesis. Proc. Natl. Acad. Sci. U. S. A. 100, 14385–14390. https://doi.org/10.1073/pnas.2334169100

Du Preez, A., Lefèvre-Arbogast, S., Houghton, V., de Lucia, C., Low, D.Y., Helmer, C., Féart, C., Delcourt, C., Proust-Lima, C., Pallàs, M., Ruigrok, S.R., Altendorfer, B., González-Domínguez, R., Sánchez-Pla, A., Urpi-Sardà, M., Andres-Lacueva, C., Aigner, L., Lucassen, P.J., Korosi, A., Manach, C., Samieri, C., Thuret, S., 2022. The serum metabolome mediates the concert of diet, exercise, and neurogenesis, determining the risk for cognitive decline and dementia. Alzheimers. Dement. 18, 654–675. https://doi.org/10.1002/alz.12428

Ekonomou, A., Savva, G.M., Brayne, C., Forster, G., Francis, P.T., Johnson, M., Perry, E.K., Attems, J., Somani, A., Minger, S.L., Ballard, C.G., 2015. Stage-specific changes in neurogenic and glial markers in Alzheimer’s disease. Biol. Psychiatry 77, 711–719. https://doi.org/10.1016/j.biopsych.2014.05.021

Ferreiro, E., Lanzillo, M., Canhoto, D., Carvalho da Silva, A.M., Mota, S.I., Dias, I.S., Ferreira, I.L., Fontes, A.R., Mastrella, G., Pinheiro, P., Valero, J., Rego, A.C., 2020. Chronic hyperglycemia impairs hippocampal neurogenesis and memory in an Alzheimer’s disease mouse model. Neurobiol. Aging 92, 98–113. https://doi.org/10.1016/j.neurobiolaging.2020.04.003

Fujii, Y., Nguyen, T.T.T., Fujimura, Y., Kameya, N., Nakamura, S., Arakawa, K., Morita, H., 2019. Fecal metabolite of a gnotobiotic mouse transplanted with gut microbiota from a patient with Alzheimer’s disease. Biosci. Biotechnol. Biochem. 83, 2144–2152. https://doi.org/10.1080/09168451.2019.1644149

Gallart-Palau, X., Serra, A., Lee, B.S.T., Guo, X., Sze, S.K., 2017. Brain ureido degenerative protein modifications are associated with neuroinflammation and proteinopathy in Alzheimer’s disease with cerebrovascular disease. J. Neuroinflammation 14, 175. https://doi.org/10.1186/s12974-017-0946-y

Gao, J., Zhou, N., Wu, Y., Lu, M., Wang, Q., Xia, C., Zhou, M., Xu, Y., 2021. Urinary metabolomic changes and microbiotic alterations in presenilin1/2 conditional double knockout mice. J. Transl. Med. 19, 351. https://doi.org/10.1186/s12967-021-03032-9

García-Corzo, L., Calatayud-Baselga, I., Casares-Crespo, L., Mora-Martínez, C., Julián Escribano-Saiz, J., Hortigüela, R., Asenjo-Martínez, A., Jordán-Pla, A., Ercoli, S., Flames, N., López-Alonso, V., Vilar, M., Mira, H., 2022. The transcription factor LEF1 interacts with NFIX and switches isoforms during adult hippocampal neural stem cell quiescence. Front. cell Dev. Biol. 10, 912319. https://doi.org/10.3389/fcell.2022.912319

Garthe, A., Kempermann, G., 2013. An old test for new neurons: refining the Morris water maze to study the functional relevance of adult hippocampal neurogenesis. Front. Neurosci. 7, 63. https://doi.org/10.3389/fnins.2013.00063

Gebara, E., Udry, F., Sultan, S., Toni, N., 2015. Taurine increases hippocampal neurogenesis in aging mice. Stem Cell Res. 14, 369–379. https://doi.org/10.1016/j.scr.2015.04.001

Gonçalves, J.T., Schafer, S.T., Gage, F.H., 2016. Adult Neurogenesis in the Hippocampus: From Stem Cells to Behavior. Cell 167, 897–914. https://doi.org/10.1016/j.cell.2016.10.021

Gong, C.-X., Liu, F., Iqbal, K., 2018. Multifactorial Hypothesis and Multi-Targets for Alzheimer’s Disease. J. Alzheimers. Dis. 64, S107–S117. https://doi.org/10.3233/JAD-179921

González-Domínguez, R., García-Barrera, T., Vitorica, J., Gómez-Ariza, J.L., 2015. High throughput multiorgan metabolomics in the APP/PS1 mouse model of Alzheimer’s disease. Electrophoresis 36, 2237–2249. https://doi.org/10.1002/elps.201400544

Goodman, T., Trouche, S., Massou, I., Verret, L., Zerwas, M., Roullet, P., Rampon, C., 2010. Young hippocampal neurons are critical for recent and remote spatial memory in adult mice. Neuroscience 171, 769–778. https://doi.org/10.1016/j.neuroscience.2010.09.047

Goyal, D., Ali, S.A., Singh, R.K., 2021. Emerging role of gut microbiota in modulation of neuroinflammation and neurodegeneration with emphasis on Alzheimer’s disease. Prog. Neuropsychopharmacol. Biol. Psychiatry 106, 110112. https://doi.org/10.1016/j.pnpbp.2020.110112

Guo, M., Peng, J., Huang, X., Xiao, L., Huang, F., Zuo, Z., 2021. Gut Microbiome Features of Chinese Patients Newly Diagnosed with Alzheimer’s Disease or Mild Cognitive Impairment. J. Alzheimers. Dis. 80, 299–310. https://doi.org/10.3233/JAD-201040

Han, Y., Quan, X., Chuang, Y., Liang, Q., Li, Y., Yuan, Z., Bian, Y., Wei, L., Wang, J., Zhao, Y., 2022. A multi-omics analysis for the prediction of neurocognitive disorders risk among the elderly in Macao. Clin. Transl. Med. 12, e909. https://doi.org/10.1002/ctm2.909

Haran, J.P., Bhattarai, S.K., Foley, S.E., Dutta, P., Ward, D. V, Bucci, V., McCormick, B.A., 2019. Alzheimer’s Disease Microbiome Is Associated with Dysregulation of the Anti-Inflammatory P-Glycoprotein Pathway. MBio 10. https://doi.org/10.1128/mBio.00632-19

He, Y., Wu, W., Zheng, H.-M., Li, P., McDonald, D., Sheng, H.-F., Chen, M.-X., Chen, Z.-H., Ji, G.-Y., Zheng, Z.-D.-X., Mujagond, P., Chen, X.-J., Rong, Z.-H., Chen, P., Lyu, L.-Y., Wang, X., Wu, C.-B., Yu, N., Xu, Y.-J., Yin, J., Raes, J., Knight, R., Ma, W.-J., Zhou, H.-W., 2018. Regional variation limits applications of healthy gut microbiome reference ranges and disease models. Nat. Med. 24, 1532–1535. https://doi.org/10.1038/s41591-018-0164-x

Heine, V.M., Maslam, S., Joëls, M., Lucassen, P.J., 2004. Prominent decline of newborn cell proliferation, differentiation, and apoptosis in the aging dentate gyrus, in absence of an age-related hypothalamus-pituitary-adrenal axis activation. Neurobiol. Aging 25, 361–375. https://doi.org/10.1016/S0197-4580(03)00090-3

Heneka, M.T., Carson, M.J., Khoury, J. El, Landreth, G.E., Brosseron, F., Feinstein, D.L., Jacobs, A.H., Wyss-Coray, T., Vitorica, J., Ransohoff, R.M., Herrup, K., Frautschy, S.A., Finsen, B., Brown, G.C., Verkhratsky, A., Yamanaka, K., Koistinaho, J., Latz, E., Halle, A., Petzold, G.C., Town, T., Morgan, D., Shinohara, M.L., Perry, V.H., Holmes, C., Bazan, N.G., Brooks, D.J., Hunot, S., Joseph, B., Deigendesch, N., Garaschuk, O., Boddeke, E., Dinarello, C.A., Breitner, J.C., Cole, G.M., Golenbock, D.T., Kummer, M.P., 2015. Neuroinflammation in Alzheimer’s disease. Lancet Neurol. https://doi.org/10.1016/S1474-4422(15)70016-5

Heneka, M.T., Kummer, M.P., Stutz, A., Delekate, A., Schwartz, S., Vieira-Saecker, A., Griep, A., Axt, D., Remus, A., Tzeng, T.-C., Gelpi, E., Halle, A., Korte, M., Latz, E., Golenbock, D.T., 2013. NLRP3 is activated in Alzheimer’s disease and contributes to pathology in APP/PS1 mice. Nature 493, 674–678. https://doi.org/10.1038/nature11729

Hernández-Benítez, R., Ramos-Mandujano, G., Pasantes-Morales, H., 2012. Taurine stimulates proliferation and promotes neurogenesis of mouse adult cultured neural stem/progenitor cells. Stem Cell Res. 9, 24–34. https://doi.org/10.1016/j.scr.2012.02.004

Holden, H.M., Gilbert, P.E., 2012. Less efficient pattern separation may contribute to age-related spatial memory deficits. Front. Aging Neurosci. 4, 9. https://doi.org/10.3389/fnagi.2012.00009

Honarpisheh, P., Reynolds, C.R., Blasco Conesa, M.P., Moruno Manchon, J.F., Putluri, N., Bhattacharjee, M.B., Urayama, A., McCullough, L.D., Ganesh, B.P., 2020. Dysregulated Gut Homeostasis Observed Prior to the Accumulation of the Brain Amyloid-β in Tg2576 Mice. Int. J. Mol. Sci. 21. https://doi.org/10.3390/ijms21051711

Hunsberger, H.C., Greenwood, B.P., Tolstikov, V., Narain, N.R., Kiebish, M.A., Denny, C.A., 2020. Divergence in the metabolome between natural aging and Alzheimer’s disease. Sci. Rep. 10, 12171. https://doi.org/10.1038/s41598-020-68739-z

Iqbal, K., Alonso, A. del C., Chen, S., Chohan, M.O., El-Akkad, E., Gong, C.-X., Khatoon, S., Li, B., Liu, F., Rahman, A., Tanimukai, H., Grundke-Iqbal, I., 2005. Tau pathology in Alzheimer disease and other tauopathies. Biochim. Biophys. Acta 1739, 198–210. https://doi.org/10.1016/j.bbadis.2004.09.008

Ising, C., Venegas, C., Zhang, S., Scheiblich, H., Schmidt, S. V, Vieira-Saecker, A., Schwartz, S., Albasset, S., McManus, R.M., Tejera, D., Griep, A., Santarelli, F., Brosseron, F., Opitz, S., Stunden, J., Merten, M., Kayed, R., Golenbock, D.T., Blum, D., Latz, E., Buée, L., Heneka, M.T., 2019. NLRP3 inflammasome activation drives tau pathology. Nature 575, 669–673. https://doi.org/10.1038/s41586-019-1769-z

Jansen, I.E., Savage, J.E., Watanabe, K., Bryois, J., Williams, D.M., Steinberg, S., Sealock, J., Karlsson, I.K., Hägg, S., Athanasiu, L., Voyle, N., Proitsi, P., Witoelar, A., Stringer, S., Aarsland, D., Almdahl, I.S., Andersen, F., Bergh, S., Bettella, F., Bjornsson, S., Brækhus, A., Bråthen, G., de Leeuw, C., Desikan, R.S., Djurovic, S., Dumitrescu, L., Fladby, T., Hohman, T.J., Jonsson, P. V, Kiddle, S.J., Rongve, A., Saltvedt, I., Sando, S.B., Selbæk, G., Shoai, M., Skene, N.G., Snaedal, J., Stordal, E., Ulstein, I.D., Wang, Y., White, L.R., Hardy, J., Hjerling-Leffler, J., Sullivan, P.F., van der Flier, W.M., Dobson, R., Davis, L.K., Stefansson, H., Stefansson, K., Pedersen, N.L., Ripke, S., Andreassen, O.A., Posthuma, D., 2019. Genome-wide meta-analysis identifies new loci and functional pathways influencing Alzheimer’s disease risk. Nat. Genet. 51, 404–413. https://doi.org/10.1038/s41588-018-0311-9

Jessberger, S., Clark, R.E., Broadbent, N.J., Clemenson, G.D.J., Consiglio, A., Lie, D.C., Squire, L.R., Gage, F.H., 2009. Dentate gyrus-specific knockdown of adult neurogenesis impairs spatial and object recognition memory in adult rats. Learn. Mem. 16, 147–154. https://doi.org/10.1101/lm.1172609

Karamujić-Čomić, H., Ahmad, S., Radjabzadeh, D., Bonnechere, B., Kaddurah-Daouk, R.F., Kraaij, R., Ikram, M.A., Amin, N., van Duijn, C.M., 2020. Clostridium shows a higher abundance in less neurovascular and neurodegenerative changes: A microbiome-wide association study. Alzheimer’s & Dement. 16, e044743. https://doi.org/https://doi.org/10.1002/alz.044743

Kim, N., Jeon, S.H., Ju, I.G., Gee, M.S., Do, J., Oh, M.S., Lee, J.K., 2021. Transplantation of gut microbiota derived from Alzheimer’s disease mouse model impairs memory function and neurogenesis in C57BL/6 mice. Brain. Behav. Immun. 98, 357–365. https://doi.org/10.1016/j.bbi.2021.09.002

Kivipelto, M., Mangialasche, F., Ngandu, T., 2018. Lifestyle interventions to prevent cognitive impairment, dementia and Alzheimer disease. Nat. Rev. Neurol. 14, 653–666. https://doi.org/10.1038/s41582-018-0070-3

Kowalski, K., Mulak, A., 2022. Small intestinal bacterial overgrowth in Alzheimer’s disease. J. Neural Transm. 129, 75–83. https://doi.org/10.1007/s00702-021-02440-x

Lambert, J.C., Ibrahim-Verbaas, C.A., Harold, D., Naj, A.C., Sims, R., Bellenguez, C., DeStafano, A.L., Bis, J.C., Beecham, G.W., Grenier-Boley, B., Russo, G., Thorton-Wells, T.A., Jones, N., Smith, A. V, Chouraki, V., Thomas, C., Ikram, M.A., Zelenika, D., Vardarajan, B.N., Kamatani, Y., Lin, C.F., Gerrish, A., Schmidt, H., Kunkle, B., Dunstan, M.L., Ruiz, A., Bihoreau, M.T., Choi, S.H., Reitz, C., Pasquier, F., Cruchaga, C., Craig, D., Amin, N., Berr, C., Lopez, O.L., De Jager, P.L., Deramecourt, V., Johnston, J.A., Evans, D., Lovestone, S., Letenneur, L., Morón, F.J., Rubinsztein, D.C., Eiriksdottir, G., Sleegers, K., Goate, A.M., Fiévet, N., Huentelman, M.W., Gill, M., Brown, K., Kamboh, M.I., Keller, L., Barberger-Gateau, P., McGuiness, B., Larson, E.B., Green, R., Myers, A.J., Dufouil, C., Todd, S., Wallon, D., Love, S., Rogaeva, E., Gallacher, J., St George-Hyslop, P., Clarimon, J., Lleo, A., Bayer, A., Tsuang, D.W., Yu, L., Tsolaki, M., Bossù, P., Spalletta, G., Proitsi, P., Collinge, J., Sorbi, S., Sanchez-Garcia, F., Fox, N.C., Hardy, J., Deniz Naranjo, M.C., Bosco, P., Clarke, R., Brayne, C., Galimberti, D., Mancuso, M., Matthews, F., Moebus, S., Mecocci, P., Del Zompo, M., Maier, W., Hampel, H., Pilotto, A., Bullido, M., Panza, F., Caffarra, P., Nacmias, B., Gilbert, J.R., Mayhaus, M., Lannefelt, L., Hakonarson, H., Pichler, S., Carrasquillo, M.M., Ingelsson, M., Beekly, D., Alvarez, V., Zou, F., Valladares, O., Younkin, S.G., Coto, E., Hamilton-Nelson, K.L., Gu, W., Razquin, C., Pastor, P., Mateo, I., Owen, M.J., Faber, K.M., Jonsson, P. V, Combarros, O., O’Donovan, M.C., Cantwell, L.B., Soininen, H., Blacker, D., Mead, S., Mosley, T.H.J., Bennett, D.A., Harris, T.B., Fratiglioni, L., Holmes, C., de Bruijn, R.F., Passmore, P., Montine, T.J., Bettens, K., Rotter, J.I., Brice, A., Morgan, K., Foroud, T.M., Kukull, W.A., Hannequin, D., Powell, J.F., Nalls, M.A., Ritchie, K., Lunetta, K.L., Kauwe, J.S., Boerwinkle, E., Riemenschneider, M., Boada, M., Hiltuenen, M., Martin, E.R., Schmidt, R., Rujescu, D., Wang, L.S., Dartigues, J.F., Mayeux, R., Tzourio, C., Hofman, A., Nöthen, M.M., Graff, C., Psaty, B.M., Jones, L., Haines, J.L., Holmans, P.A., Lathrop, M., Pericak-Vance, M.A., Launer, L.J., Farrer, L.A., van Duijn, C.M., Van Broeckhoven, C., Moskvina, V., Seshadri, S., Williams, J., Schellenberg, G.D., Amouyel, P., 2013. Meta-analysis of 74,046 individuals identifies 11 new susceptibility loci for Alzheimer’s disease. Nat. Genet. 45, 1452–1458. https://doi.org/10.1038/ng.2802

Leal, S.L., Ferguson, L.A., Harrison, T.M., Jagust, W.J., 2019. Development of a mnemonic discrimination task using naturalistic stimuli with applications to aging and preclinical Alzheimer’s disease. Learn. Mem. 26, 219–228. https://doi.org/10.1101/lm.048967.118

Leblhuber, F., Geisler, S., Steiner, K., Fuchs, D., Schütz, B., 2015. Elevated fecal calprotectin in patients with Alzheimer’s dementia indicates leaky gut. J. Neural Transm. 122, 1319–1322. https://doi.org/10.1007/s00702-015-1381-9

Lee, K.S., Chung, J.H., Lee, K.H., Shin, M.-J., Oh, B.H., Hong, C.H., 2008. Bioplex analysis of plasma cytokines in Alzheimer’s disease and mild cognitive impairment. Immunol. Lett. 121, 105–109. https://doi.org/10.1016/j.imlet.2008.09.004

Li, B., He, Y., Ma, J., Huang, P., Du, J., Cao, L., Wang, Y., Xiao, Q., Tang, H., Chen, S., 2019. Mild cognitive impairment has similar alterations as Alzheimer’s disease in gut microbiota. Alzheimers. Dement. 15, 1357–1366. https://doi.org/10.1016/j.jalz.2019.07.002

Li, Y., Ning, L., Yin, Y., Wang, R., Zhang, Z., Hao, L., Wang, B., Zhao, X., Yang, X., Yin, L., Wu, S., Guo, D., Zhang, C., 2020. Age-related shifts in gut microbiota contribute to cognitive decline in aged rats. Aging (Albany. NY). 12, 7801–7817. https://doi.org/10.18632/aging.103093

Licht, T., Keshet, E., 2015. The vascular niche in adult neurogenesis. Mech. Dev. 138 Pt 1, 56–62. https://doi.org/10.1016/j.mod.2015.06.001

Ling, Z., Zhu, M., Liu, X., Shao, L., Cheng, Y., Yan, X., Jiang, R., Wu, S., 2020a. Fecal Fungal Dysbiosis in Chinese Patients With Alzheimer’s Disease. Front. cell Dev. Biol. 8, 631460. https://doi.org/10.3389/fcell.2020.631460

Ling, Z., Zhu, M., Yan, X., Cheng, Y., Shao, L., Liu, X., Jiang, R., Wu, S., 2020b. Structural and Functional Dysbiosis of Fecal Microbiota in Chinese Patients With Alzheimer’s Disease. Front. cell Dev. Biol. 8, 634069. https://doi.org/10.3389/fcell.2020.634069

Liu, G., Yu, Q., Tan, B., Ke, X., Zhang, C., Li, H., Zhang, T., Lu, Y., 2022. Gut dysbiosis impairs hippocampal plasticity and behaviors by remodeling serum metabolome. Gut Microbes 14, 2104089. https://doi.org/10.1080/19490976.2022.2104089

Liu, P., Wu, L., Peng, G., Han, Y., Tang, R., Ge, J., Zhang, L., Jia, L., Yue, S., Zhou, K., Li, L., Luo, B., Wang, B., 2019. Altered microbiomes distinguish Alzheimer’s disease from amnestic mild cognitive impairment and health in a Chinese cohort. Brain. Behav. Immun. 80, 633–643. https://doi.org/10.1016/j.bbi.2019.05.008

Liu, S., Gao, J., Zhu, M., Liu, K., Zhang, H.-L., 2020. Gut Microbiota and Dysbiosis in Alzheimer’s Disease: Implications for Pathogenesis and Treatment. Mol. Neurobiol. 57, 5026–5043. https://doi.org/10.1007/s12035-020-02073-3

Long, J.M., Holtzman, D.M., 2019. Alzheimer Disease: An Update on Pathobiology and Treatment Strategies. Cell 179, 312–339. https://doi.org/10.1016/j.cell.2019.09.001

Marizzoni, M., Cattaneo, A., Mirabelli, P., Festari, C., Lopizzo, N., Nicolosi, V., Mombelli, E., Mazzelli, M., Luongo, D., Naviglio, D., Coppola, L., Salvatore, M., Frisoni, G.B., 2020a. Short-Chain Fatty Acids and Lipopolysaccharide as Mediators Between Gut Dysbiosis and Amyloid Pathology in Alzheimer’s Disease. J. Alzheimers. Dis. 78, 683–697. https://doi.org/10.3233/JAD-200306

Marizzoni, M., Gurry, T., Provasi, S., Greub, G., Lopizzo, N., Ribaldi, F., Festari, C., Mazzelli, M., Mombelli, E., Salvatore, M., Mirabelli, P., Franzese, M., Soricelli, A., Frisoni, G.B., Cattaneo, A., 2020b. Comparison of Bioinformatics Pipelines and Operating Systems for the Analyses of 16S rRNA Gene Amplicon Sequences in Human Fecal Samples. Front. Microbiol. 11. https://doi.org/10.3389/fmicb.2020.01262

Mazurkiewicz-Kwilecki, I.M., Nsonwah, S., 1989. Changes in the regional brain histamine and histidine levels in postmortem brains of Alzheimer patients. Can. J. Physiol. Pharmacol. 67, 75–78. https://doi.org/10.1139/y89-013

Minter, M.R., Zhang, C., Leone, V., Ringus, D.L., Zhang, X., Oyler-Castrillo, P., Musch, M.W., Liao, F., Ward, J.F., Holtzman, D.M., Chang, E.B., Tanzi, R.E., Sisodia, S.S., 2016. Antibiotic-induced perturbations in gut microbial diversity influences neuro-inflammation and amyloidosis in a murine model of Alzheimer’s disease. Sci. Rep. 6, 30028. https://doi.org/10.1038/srep30028

Mishra, R., Phan, T., Kumar, P., Morrissey, Z., Gupta, M., Hollands, C., Shetti, A., Lopez, K.L., Maienschein-Cline, M., Suh, H., Hen, R., Lazarov, O., 2022. Augmenting neurogenesis rescues memory impairments in Alzheimer’s disease by restoring the memory-storing neurons. J. Exp. Med. 219. https://doi.org/10.1084/jem.20220391

Möhle, L., Mattei, D., Heimesaat, M.M., Bereswill, S., Fischer, A., Alutis, M., French, T., Hambardzumyan, D., Matzinger, P., Dunay, I.R., Wolf, S.A., 2016. Ly6C(hi) Monocytes Provide a Link between Antibiotic-Induced Changes in Gut Microbiota and Adult Hippocampal Neurogenesis. Cell Rep. 15, 1945–1956. https://doi.org/10.1016/j.celrep.2016.04.074

Moreno-Jimenez, E.P., Flor-Garcia, M., Terreros-Roncal, J., Rabano, A., Cafini, F., Pallas-Bazarra, N., Avila, J., Llorens-Martin, M., 2019. Adult hippocampal neurogenesis is abundant in neurologically healthy subjects and drops sharply in patients with Alzheimer’s disease. Nat. Med. 25, 554–560. https://doi.org/10.1038/s41591-019-0375-9

Nielsen, J.E., Maltesen, R.G., Havelund, J.F., Færgeman, N.J., Gotfredsen, C.H., Vestergård, K., Kristensen, S.R., Pedersen, S., 2021. Characterising Alzheimer’s disease through integrative NMR- and LC-MS-based metabolomics. Metab. open 12, 100125. https://doi.org/10.1016/j.metop.2021.100125

Nomura, K., Ishikawa, D., Okahara, K., Ito, S., Haga, K., Takahashi, M., Arakawa, A., Shibuya, T., Osada, T., Kuwahara-Arai, K., Kirikae, T., Nagahara, A., 2021. Bacteroidetes Species Are Correlated with Disease Activity in Ulcerative Colitis. J. Clin. Med. 10. https://doi.org/10.3390/jcm10081749

Ogbonnaya, E.S., Clarke, G., Shanahan, F., Dinan, T.G., Cryan, J.F., O’Leary, O.F., 2015. Adult Hippocampal Neurogenesis Is Regulated by the Microbiome. Biol. Psychiatry. https://doi.org/10.1016/j.biopsych.2014.12.023

Palmas, V., Pisanu, S., Madau, V., Casula, E., Deledda, A., Cusano, R., Uva, P., Loviselli, A., Velluzzi, F., Manzin, A., 2022. Gut Microbiota Markers and Dietary Habits Associated with Extreme Longevity in Healthy Sardinian Centenarians. Nutrients 14. https://doi.org/10.3390/nu14122436

Parizkova, M., Lerch, O., Andel, R., Kalinova, J., Markova, H., Vyhnalek, M., Hort, J., Laczó, J., 2020. Spatial Pattern Separation in Early Alzheimer’s Disease. J. Alzheimers. Dis. 76, 121–138. https://doi.org/10.3233/JAD-200093

Pellegrini, C., Daniele, S., Antonioli, L., Benvenuti, L., D’Antongiovanni, V., Piccarducci, R., Pietrobono, D., Citi, V., Piragine, E., Flori, L., Ippolito, C., Segnani, C., Palazon-Riquelme, P., Lopez-Castejon, G., Martelli, A., Colucci, R., Bernardini, N., Trincavelli, M.L., Calderone, V., Martini, C., Blandizzi, C., Fornai, M., 2020. Prodromal Intestinal Events in Alzheimer’s Disease (AD): Colonic Dysmotility and Inflammation Are Associated with Enteric AD-Related Protein Deposition. Int. J. Mol. Sci. 21. https://doi.org/10.3390/ijms21103523

Pluvinage, J. V, Wyss-Coray, T., 2020. Author Correction: Systemic factors as mediators of brain homeostasis, ageing and neurodegeneration. Nat. Rev. Neurosci. 21, 298. https://doi.org/10.1038/s41583-020-0293-3

Popp, J., Bacher, M., Kölsch, H., Noelker, C., Deuster, O., Dodel, R., Jessen, F., 2009. Macrophage migration inhibitory factor in mild cognitive impairment and Alzheimer’s disease. J. Psychiatr. Res. 43, 749–753. https://doi.org/10.1016/j.jpsychires.2008.10.006

Rei, D., Saha, S., Haddad, M., Haider Rubio, A., Perlaza, B.L., Berard, M., Ungeheuer, M.-N., Sokol, H., Lledo, P.-M., 2022. Age-associated gut microbiota impairs hippocampus-dependent memory in a vagus-dependent manner. JCI insight. https://doi.org/10.1172/jci.insight.147700

Ren, T., Gao, Y., Qiu, Y., Jiang, S., Zhang, Q., Zhang, J., Wang, Limin, Zhang, Y., Wang, Lijuan, Nie, K., 2020. Gut Microbiota Altered in Mild Cognitive Impairment Compared With Normal Cognition in Sporadic Parkinson’s Disease. Front. Neurol. 11, 137. https://doi.org/10.3389/fneur.2020.00137

Robinson, M., Lee, B.Y., Hane, F.T., 2017. Recent Progress in Alzheimer’s Disease Research, Part 2: Genetics and Epidemiology. J. Alzheimers. Dis. 57, 317–330. https://doi.org/10.3233/JAD-161149

Rodríguez-Iglesias, N., Sierra, A., Valero, J., 2019. Rewiring of Memory Circuits: Connecting Adult Newborn Neurons With the Help of Microglia. Front. cell Dev. Biol. 7, 24. https://doi.org/10.3389/fcell.2019.00024

Rosenzweig, S., Wojtowicz, J.M., 2011. Analyzing dendritic growth in a population of immature neurons in the adult dentate gyrus using laminar quantification of disjointed dendrites. Front. Neurosci. 5, 34. https://doi.org/10.3389/fnins.2011.00034

Ryan, S.M., O’Keeffe, G.W., O’Connor, C., Keeshan, K., Nolan, Y.M., 2013. Negative regulation of TLX by IL-1β correlates with an inhibition of adult hippocampal neural precursor cell proliferation. Brain. Behav. Immun. 33, 7–13. https://doi.org/10.1016/j.bbi.2013.03.005

Saad, Y., Segal, D., Ayali, A., 2015. Enhanced Neurite Outgrowth and Branching Precede Increased Amyloid-β-Induced Neuronal Apoptosis in a Novel Alzheimer’s Disease Model. J. Alzheimer’s Dis. 43, 993–1006. https://doi.org/10.3233/JAD-140009

Sahay, A., Scobie, K.N., Hill, A.S., O’Carroll, C.M., Kheirbek, M.A., Burghardt, N.S., Fenton, A.A., Dranovsky, A., Hen, R., 2011. Increasing adult hippocampal neurogenesis is sufficient to improve pattern separation. Nature 472, 466–470. https://doi.org/10.1038/nature09817

Saresella, M., La Rosa, F., Piancone, F., Zoppis, M., Marventano, I., Calabrese, E., Rainone, V., Nemni, R., Mancuso, R., Clerici, M., 2016. The NLRP3 and NLRP1 inflammasomes are activated in Alzheimer’s disease. Mol. Neurodegener. 11, 23. https://doi.org/10.1186/s13024-016-0088-1

Sawin, E.A., De Wolfe, T.J., Aktas, B., Stroup, B.M., Murali, S.G., Steele, J.L., Ney, D.M., 2015. Glycomacropeptide is a prebiotic that reduces Desulfovibrio bacteria, increases cecal short-chain fatty acids, and is anti-inflammatory in mice. Am. J. Physiol. Gastrointest. Liver Physiol. 309, G590–601. https://doi.org/10.1152/ajpgi.00211.2015

Selkoe, D.J., Hardy, J., 2016. The amyloid hypothesis of Alzheimer’s disease at 25 years. EMBO Mol. Med. 8, 595–608. https://doi.org/10.15252/emmm.201606210

Shivaraj, M.C., Marcy, G., Low, G., Ryu, J.R., Zhao, X., Rosales, F.J., Goh, E.L.K., 2012. Taurine induces proliferation of neural stem cells and synapse development in the developing mouse brain. PLoS One 7, e42935. https://doi.org/10.1371/journal.pone.0042935

Sinha, N., Berg, C.N., Tustison, N.J., Shaw, A., Hill, D., Yassa, M.A., Gluck, M.A., 2018. APOE ε4 status in healthy older African Americans is associated with deficits in pattern separation and hippocampal hyperactivation. Neurobiol. Aging 69, 221–229. https://doi.org/10.1016/j.neurobiolaging.2018.05.023

Slingerland, A.E., Schwabkey, Z., Wiesnoski, D.H., Jenq, R.R., 2017. Clinical Evidence for the Microbiome in Inflammatory Diseases. Front. Immunol. 8, 400. https://doi.org/10.3389/fimmu.2017.00400

Smith, D.G., Ciccotosto, G.D., Tew, D.J., Perez, K., Curtain, C.C., Boas, J.F., Masters, C.L., Cappai, R., Barnham, K.J., 2010. Histidine 14 modulates membrane binding and neurotoxicity of the Alzheimer’s disease amyloid-beta peptide. J. Alzheimers. Dis. 19, 1387–1400. https://doi.org/10.3233/JAD-2010-1334

Smith, L.K., White 3rd, C.W., Villeda, S.A., 2018. The systemic environment: at the interface of aging and adult neurogenesis. Cell Tissue Res. 371, 105–113. https://doi.org/10.1007/s00441-017-2715-8

Song, J., Yang, L., Nan, D., He, Q., Wan, Y., Guo, H., 2018. Histidine Alleviates Impairments Induced by Chronic Cerebral Hypoperfusion in Mice. Front. Physiol. 9, 662. https://doi.org/10.3389/fphys.2018.00662

Squire, L.R., 1992. Memory and the hippocampus: a synthesis from findings with rats, monkeys, and humans. Psychol. Rev. 99, 195–231. https://doi.org/10.1037/0033-295x.99.2.195

Stadlbauer, V., Engertsberger, L., Komarova, I., Feldbacher, N., Leber, B., Pichler, G., Fink, N., Scarpatetti, M., Schippinger, W., Schmidt, R., Horvath, A., 2020. Dysbiosis, gut barrier dysfunction and inflammation in dementia: a pilot study. BMC Geriatr. 20, 248. https://doi.org/10.1186/s12877-020-01644-2

Stojanov, S., Berlec, A., Štrukelj, B., 2020. The Influence of Probiotics on the Firmicutes/Bacteroidetes Ratio in the Treatment of Obesity and Inflammatory Bowel disease. Microorganisms 8. https://doi.org/10.3390/microorganisms8111715

Sun, J., Xu, J., Ling, Y., Wang, F., Gong, T., Yang, C., Ye, S., Ye, K., Wei, D., Song, Z., Chen, D., Liu, J., 2019. Fecal microbiota transplantation alleviated Alzheimer’s disease-like pathogenesis in APP/PS1 transgenic mice. Transl. Psychiatry 9, 189. https://doi.org/10.1038/s41398-019-0525-3

Tachikawa, M., Hosoya, K.-I., 2011. Transport characteristics of guanidino compounds at the blood-brain barrier and blood-cerebrospinal fluid barrier: relevance to neural disorders. Fluids Barriers CNS 8, 13. https://doi.org/10.1186/2045-8118-8-13

Tanzi, R.E., Bertram, L., 2005. Twenty years of the Alzheimer’s disease amyloid hypothesis: a genetic perspective. Cell 120, 545–555. https://doi.org/10.1016/j.cell.2005.02.008

Tobin, M.K., Musaraca, K., Disouky, A., Shetti, A., Bheri, A., Honer, W.G., Kim, N., Dawe, R.J., Bennett, D.A., Arfanakis, K., Lazarov, O., 2019. Human Hippocampal Neurogenesis Persists in Aged Adults and Alzheimer’s Disease Patients. Cell Stem Cell 24, 974–982.e3. https://doi.org/10.1016/j.stem.2019.05.003

Varesi, A., Pierella, E., Romeo, M., Piccini, G.B., Alfano, C., Bjørklund, G., Oppong, A., Ricevuti, G., Esposito, C., Chirumbolo, S., Pascale, A., 2022. The Potential Role of Gut Microbiota in Alzheimer’s Disease: From Diagnosis to Treatment. Nutrients 14. https://doi.org/10.3390/nu14030668

Veitch, D.P., Weiner, M.W., Aisen, P.S., Beckett, L.A., Cairns, N.J., Green, R.C., Harvey, D., Jack, C.R.J., Jagust, W., Morris, J.C., Petersen, R.C., Saykin, A.J., Shaw, L.M., Toga, A.W., Trojanowski, J.Q., 2019. Understanding disease progression and improving Alzheimer’s disease clinical trials: Recent highlights from the Alzheimer’s Disease Neuroimaging Initiative. Alzheimers. Dement. 15, 106–152. https://doi.org/10.1016/j.jalz.2018.08.005

Verhaar, B.J.H., Hendriksen, H.M.A., de Leeuw, F.A., Doorduijn, A.S., van Leeuwenstijn, M., Teunissen, C.E., Barkhof, F., Scheltens, P., Kraaij, R., van Duijn, C.M., Nieuwdorp, M., Muller, M., van der Flier, W.M., 2021. Gut Microbiota Composition Is Related to AD Pathology. Front. Immunol. 12, 794519. https://doi.org/10.3389/fimmu.2021.794519

Villeda, S.A., Luo, J., Mosher, K.I., Zou, B., Britschgi, M., Bieri, G., Stan, T.M., Fainberg, N., Ding, Z., Eggel, A., Lucin, K.M., Czirr, E., Park, J.-S., CouillardDesprés, S., Aigner, L., Li, G., Peskind, E.R., Kaye, J.A., Quinn, J.F., Galasko, D.R., Xie, X.S., Rando, T.A., Wyss-Coray, T., Couillard-Despres, S., Aigner, L., Li, G., Peskind, E.R., Kaye, J.A., Quinn, J.F., Galasko, D.R., Xie, X.S., Rando, T.A., Wyss-Coray, T., 2011. The ageing systemic milieu negatively regulates neurogenesis and cognitive function. Nature 477, 90–94. https://doi.org/10.1038/nature10357

Villeda, S.A., Plambeck, K.E., Middeldorp, J., Castellano, J.M., Mosher, K.I., Luo, J., Smith, L.K., Bieri, G., Lin, K., Berdnik, D., Wabl, R., Udeochu, J., Wheatley, E.G., Zou, B., Simmons, D.A., Xie, X.S., Longo, F.M., Wyss-Coray, T., 2014. Young blood reverses age-related impairments in cognitive function and synaptic plasticity in mice. Nat. Med. 20, 659–663. https://doi.org/10.1038/nm.3569

Vogt, N.M., Kerby, R.L., Dill-McFarland, K.A., Harding, S.J., Merluzzi, A.P., Johnson, S.C., Carlsson, C.M., Asthana, S., Zetterberg, H., Blennow, K., Bendlin, B.B., Rey, F.E., 2017. Gut microbiome alterations in Alzheimer’s disease. Sci. Rep. 7, 13537. https://doi.org/10.1038/s41598-017-13601-y

Walgrave, H., Balusu, S., Snoeck, S., Vanden Eynden, E., Craessaerts, K., Thrupp, N., Wolfs, L., Horré, K., Fourne, Y., Ronisz, A., Silajdžić, E., Penning, A., Tosoni, G., Callaerts-Vegh, Z., D’Hooge, R., Thal, D.R., Zetterberg, H., Thuret, S., Fiers, M., Frigerio, C.S., De Strooper, B., Salta, E., 2021. Restoring miR-132 expression rescues adult hippocampal neurogenesis and memory deficits in Alzheimer’s disease. Cell Stem Cell 28, 1805–1821.e8. https://doi.org/10.1016/j.stem.2021.05.001

Wang, G., Zhou, Y., Huang, F.-J., Tang, H.-D., Xu, X.-H., Liu, J.-J., Wang, Y., Deng, Y.-L., Ren, R.-J., Xu, W., Ma, J.-F., Zhang, Y.-N., Zhao, A.-H., Chen, S.-D., Jia, W., 2014. Plasma metabolite profiles of Alzheimer’s disease and mild cognitive impairment. J. Proteome Res. 13, 2649–2658. https://doi.org/10.1021/pr5000895

Wang, J.-H., Guo, L., Wang, S., Yu, N.-W., Guo, F.-Q., 2022. The potential pharmacological mechanisms of β-hydroxybutyrate for improving cognitive functions. Curr. Opin. Pharmacol. 62, 15–22. https://doi.org/10.1016/j.coph.2021.10.005

Webb, C.E., Foster, C.M., Horn, M.M., Kennedy, K.M., Rodrigue, K.M., 2020. Beta-amyloid burden predicts poorer mnemonic discrimination in cognitively normal older adults. Neuroimage 221, 117199. https://doi.org/10.1016/j.neuroimage.2020.117199

Whiley, L., Chappell, K.E., D’Hondt, E., Lewis, M.R., Jiménez, B., Snowden, S.G., Soininen, H., Kłoszewska, I., Mecocci, P., Tsolaki, M., Vellas, B., Swann, J.R., Hye, A., Lovestone, S., Legido-Quigley, C., Holmes, E., 2021. Metabolic phenotyping reveals a reduction in the bioavailability of serotonin and kynurenine pathway metabolites in both the urine and serum of individuals living with Alzheimer’s disease. Alzheimers. Res. Ther. 13, 20. https://doi.org/10.1186/s13195-020-00741-z

Wu, G., Zhou, J., Yang, M., Xu, C., Pang, H., Qin, X., Lin, S., Yang, J., Hu, J., 2022. The Regulatory Effects of Taurine on Neurogenesis and Apoptosis of Neural Stem Cells in the Hippocampus of Rats. Adv. Exp. Med. Biol. 1370, 351–367. https://doi.org/10.1007/978-3-030-93337-1_34

Xi, J., Ding, D., Zhu, H., Wang, R., Su, F., Wu, W., Xiao, Z., Liang, X., Zhao, Q., Hong, Z., Fu, H., Xiao, Q., 2021. Disturbed microbial ecology in Alzheimer’s disease: evidence from the gut microbiota and fecal metabolome. BMC Microbiol. 21, 226. https://doi.org/10.1186/s12866-021-02286-z

Xie, K., Qin, Q., Long, Z., Yang, Y., Peng, C., Xi, C., Li, L., Wu, Z., Daria, V., Zhao, Y., Wang, F., Wang, M., 2021. High-Throughput Metabolomics for Discovering Potential Biomarkers and Identifying Metabolic Mechanisms in Aging and Alzheimer’s Disease. Front. cell Dev. Biol. 9, 602887. https://doi.org/10.3389/fcell.2021.602887

Yassa, M.A., Stark, C.E.L., 2011. Pattern separation in the hippocampus. Trends Neurosci. 34, 515–525. https://doi.org/10.1016/j.tins.2011.06.006

Zhan, X., Stamova, B., Jin, L.W., Decarli, C., Phinney, B., Sharp, F.R., 2016. Gram-negative bacterial molecules associate with Alzheimer disease pathology. Neurology 87, 2324–2332. https://doi.org/10.1212/WNL.0000000000003391

Zhan, X., Stamova, B., Sharp, F.R., 2018. Lipopolysaccharide Associates with Amyloid Plaques, Neurons and Oligodendrocytes in Alzheimer’s Disease Brain: A Review. Front. Aging Neurosci..

Zhao, Y., Chen, H., Iqbal, J., Liu, X., Zhang, H., Xiao, S., Jin, N., Yao, F., Shen, L., 2021. Targeted metabolomics study of early pathological features in hippocampus of triple transgenic Alzheimer’s disease male mice. J. Neurosci. Res. 99, 927–946. https://doi.org/10.1002/jnr.24750

Zhao, Y., Jaber, V., Lukiw, W.J., 2017. Secretory Products of the Human GI Tract Microbiome and Their Potential Impact on Alzheimer’s Disease (AD): Detection of Lipopolysaccharide (LPS) in AD Hippocampus. Front. Cell. Infect. Microbiol. 7, 318. https://doi.org/10.3389/fcimb.2017.00318

Zhuang, Z.-Q., Shen, L.-L., Li, W.-W., Fu, X., Zeng, F., Gui, L., Lu, Y., Cai, M., Zhu, C., Tan, Y.-L., Zheng, P., Li, H.-Y., Zhu, J., Zhou, H.-D., Bu, X.-L., Wang, Y.-J., 2018. Gut Microbiota is Altered in Patients with Alzheimer’s Disease. J. Alzheimers. Dis. 63, 1337–1346. https://doi.org/10.3233/JAD-180176

## References

Anacker, C., Cattaneo, A., Musaelyan, K., Zunszain, P.A., Horowitz, M., Molteni, R., Luoni, A., Calabrese, F., Tansey, K., Gennarelli, M., Thuret, S., Price, J., Uher, R., Riva, M.A., Pariante, C.M., 2013. Role for the kinase SGK1 in stress, depression, and glucocorticoid effects on hippocampal neurogenesis. Proc. Natl. Acad. Sci. U. S. A. 110, 8708–8713. https://doi.org/10.1073/pnas.1300886110

Anacker, C., Zunszain, P.A., Cattaneo, A., Carvalho, L.A., Garabedian, M.J., Thuret, S., Price, J., Pariante, C.M., 2011. Antidepressants increase human hippocampal neurogenesis by activating the glucocorticoid receptor. Mol. Psychiatry 16, 738–750. https://doi.org/10.1038/mp.2011.26

Bialkowska, A.B., Ghaleb, A.M., Nandan, M.O., Yang, V.W., 2016. Improved Swiss-rolling Technique for Intestinal Tissue Preparation for Immunohistochemical and Immunofluorescent Analyses. J. Vis. Exp. https://doi.org/10.3791/54161

Bokulich, N.A., Dillon, M.R., Zhang, Y., Rideout, J.R., Bolyen, E., Li, H., Albert, P.S., Caporaso, J.G., 2018. q2-longitudinal: Longitudinal and Paired-Sample Analyses of Microbiome Data. mSystems 3. https://doi.org/10.1128/mSystems.00219-18

Bolyen, E., Rideout, J.R., Dillon, M.R., Bokulich, N.A., Abnet, C.C., Al-Ghalith, G.A., Alexander, H., Alm, E.J., Arumugam, M., Asnicar, F., Bai, Y., Bisanz, J.E., Bittinger, K., Brejnrod, A., Brislawn, C.J., Brown, C.T., Callahan, B.J., Caraballo-Rodriguez, A.M., Chase, J., Cope, E.K., Da Silva, R., Diener, C., Dorrestein, P.C., Douglas, G.M., Durall, D.M., Duvallet, C., Edwardson, C.F., Ernst, M., Estaki, M., Fouquier, J., Gauglitz, J.M., Gibbons, S.M., Gibson, D.L., Gonzalez, A., Gorlick, K., Guo, J., Hillmann, B., Holmes, S., Holste, H., Huttenhower, C., Huttley, G.A., Janssen, S., Jarmusch, A.K., Jiang, L., Kaehler, B.D., Kang, K. Bin, Keefe, C.R., Keim, P., Kelley, S.T., Knights, D., Koester, I., Kosciolek, T., Kreps, J., Langille, M.G.I., Lee, J., Ley, R., Liu, Y.-X., Loftfield, E., Lozupone, C., Maher, M., Marotz, C., Martin, B.D., McDonald, D., McIver, L.J., Melnik, A. V, Metcalf, J.L., Morgan, S.C., Morton, J.T., Naimey, A.T., Navas-Molina, J.A., Nothias, L.F., Orchanian, S.B., Pearson, T., Peoples, S.L., Petras, D., Preuss, M.L., Pruesse, E., Rasmussen, L.B., Rivers, A., Robeson, M.S. 2nd, Rosenthal, P., Segata, N., Shaffer, M., Shiffer, A., Sinha, R., Song, S.J., Spear, J.R., Swafford, A.D., Thompson, L.R., Torres, P.J., Trinh, P., Tripathi, A., Turnbaugh, P.J., Ul-Hasan, S., van der Hooft, J.J.J., Vargas, F., Vazquez-Baeza, Y., Vogtmann, E., von Hippel, M., Walters, W., Wan, Y., Wang, M., Warren, J., Weber, K.C., Williamson, C.H.D., Willis, A.D., Xu, Z.Z., Zaneveld, J.R., Zhang, Y., Zhu, Q., Knight, R., Caporaso, J.G., 2019. Author Correction: Reproducible, interactive, scalable and extensible microbiome data science using QIIME 2. Nat. Biotechnol. https://doi.org/10.1038/s41587-019-0252-6

Bruce-Keller, A.J., Salbaum, J.M., Luo, M., Blanchard, E. 4th, Taylor, C.M., Welsh, D.A., Berthoud, H.-R., 2015. Obese-type gut microbiota induce neurobehavioral changes in the absence of obesity. Biol. Psychiatry 77, 607–615. https://doi.org/10.1016/j.biopsych.2014.07.012

Doneanu, C.E., Chen, W., Mazzeo, J.R., 2011. UPLC/MS Monitoring of Water-Soluble Vitamin Bs in Cell Culture Media in Minutes. Water Appl. Note 720004042en.

Fernandes, A.D., Reid, J.N., Macklaim, J.M., McMurrough, T.A., Edgell, D.R., Gloor, G.B., 2014. Unifying the analysis of high-throughput sequencing datasets: characterizing RNA-seq, 16S rRNA gene sequencing and selective growth experiments by compositional data analysis. Microbiome 2, 15. https://doi.org/10.1186/2049-2618-2-15

Folstein, M.F., Folstein, S.E., McHugh, P.R., 1975. Mini-Mental State: A practice method for grading the cognitive state of patients for the clinician. J Psychiatr Res 12, 189–198.

Gheorghe, C.E., Ritz, N.L., Martin, J.A., Wardill, H.R., Cryan, J.F., Clarke, G., 2021. Investigating causality with fecal microbiota transplantation in rodents: applications, recommendations and pitfalls. Gut Microbes 13, 1941711. https://doi.org/10.1080/19490976.2021.1941711

Gibbons, S.M., Duvallet, C., Alm, E.J., 2018. Correcting for batch effects in case-control microbiome studies. PLoS Comput. Biol. 14, e1006102. https://doi.org/10.1371/journal.pcbi.1006102

Hoban, A.E., Moloney, R.D., Golubeva, A. V, McVey Neufeld, K.A., O’Sullivan, O., Patterson, E., Stanton, C., Dinan, T.G., Clarke, G., Cryan, J.F., 2016. Behavioural and neurochemical consequences of chronic gut microbiota depletion during adulthood in the rat. Neuroscience 339, 463–477. https://doi.org/10.1016/j.neuroscience.2016.10.003

Huber, W., von Heydebreck, A., Sültmann, H., Poustka, A., Vingron, M., 2002. Variance stabilization applied to microarray data calibration and to the quantification of differential expression. Bioinformatics 18 Suppl 1, S96–104. https://doi.org/10.1093/bioinformatics/18.suppl_1.s96

Hueston, C.M., O’Leary, J.D., Hoban, A.E., Kozareva, D.A., Pawley, L.C., O’Leary, O.F., Cryan, J.F., Nolan, Y.M., 2018. Chronic interleukin-1beta in the dorsal hippocampus impairs behavioural pattern separation. Brain. Behav. Immun. 74, 252–264. https://doi.org/10.1016/j.bbi.2018.09.015

Jauhiainen, A., Madhu, B., Narita, Masako, Narita, Masashi, Griffiths, J., Tavaré, S., 2014. Normalization of metabolomics data with applications to correlation maps. Bioinformatics 30, 2155–2161. https://doi.org/10.1093/bioinformatics/btu175

Kozareva, D.A., Hueston, C.M., O’Leime, C.S., Crotty, S., Dockery, P., Cryan, J.F., Nolan, Y.M., 2019. Absence of the neurogenesis-dependent nuclear receptor TLX induces inflammation in the hippocampus. J. Neuroimmunol. 331, 87–96. https://doi.org/10.1016/j.jneuroim.2017.08.008

Li, B., Tang, J., Yang, Q., Cui, X., Li, S., Chen, S., Cao, Q., Xue, W., Chen, N., Zhu, F., 2016. Performance Evaluation and Online Realization of Data-driven Normalization Methods Used in LC/MS based Untargeted Metabolomics Analysis. Sci. Rep. 6, 38881. https://doi.org/10.1038/srep38881

Longair, M.H., Baker, D.A., Armstrong, J.D., 2011. Simple Neurite Tracer: open source software for reconstruction, visualization and analysis of neuronal processes. Bioinformatics 27, 2453–2454. https://doi.org/10.1093/bioinformatics/btr390

McKhann, G., Knopman, D.S., Chertkow, H., Hymann, B., Jack, C.R., Kawas, C., Klunk, W., Koroshetz, W., Manly, J., Mayeux, R., Mohs, R., Morris, J., Rossor, M., Scheltens, P., Carrillo, M., Weintrub, S., Phelphs, C., 2011. The diagnosis of dementia due to Alzheimer’s disease: Recommendations from the National Institute on Aging-Alzheimer’s Association workgroups on diagnostic guidelines for Alzheimer’s disease. Alzheimers Dement. 7, 263–269.

Oksanen J, Simpson G, Blanchet F, Kindt R, Legendre P, Minchin P, O’Hara R, Solymos P, S.M., Szoecs E, Wagner H, Barbour M, Bedward M, Bolker B, Borcard D, Carvalho G, Chirico M, D.C.M., Durand S, Evangelista H, FitzJohn R, Friendly M, Furneaux B, Hannigan G, Hill M, Lahti L, M.D., Ouellette M, Ribeiro Cunha E, Smith T, Stier A, Ter Braak C, W.J., 2022. vegan: Community Ecology Package. R package version 2.6-2.

Plümpe, T., Ehninger, D., Steiner, B., Klempin, F., Jessberger, S., Brandt, M., Römer, B., Rodriguez, G.R., Kronenberg, G., Kempermann, G., 2006. Variability of doublecortin-associated dendrite maturation in adult hippocampal neurogenesis is independent of the regulation of precursor cell proliferation. BMC Neurosci. 7, 77. https://doi.org/10.1186/1471-2202-7-77

Pollock, K., Stroemer, P., Patel, S., Stevanato, L., Hope, A., Miljan, E., Dong, Z., Hodges, H., Price, J., Sinden, J.D., 2006. A conditionally immortal clonal stem cell line from human cortical neuroepithelium for the treatment of ischemic stroke. Exp. Neurol. 199, 143–155. https://doi.org/10.1016/j.expneurol.2005.12.011

Quast, C., Pruesse, E., Yilmaz, P., Gerken, J., Schweer, T., Yarza, P., Peplies, J., Glöckner, F.O., 2013. The SILVA ribosomal RNA gene database project: Improved data processing and web-based tools. Nucleic Acids Res. 41(Databas, D590–6. https://doi.org/10.1093/nar/gks1219

Ritchie, M.E., Phipson, B., Wu, D., Hu, Y., Law, C.W., Shi, W., Smyth, G.K., 2015. limma powers differential expression analyses for RNA-sequencing and microarray studies. Nucleic Acids Res. 43, e47. https://doi.org/10.1093/nar/gkv007

Rocke, D.M., Durbin, B., 2001. A model for measurement error for gene expression arrays. J. Comput. Biol. a J. Comput. Mol. cell Biol. 8, 557–569. https://doi.org/10.1089/106652701753307485

Sampson, T.R., Debelius, J.W., Thron, T., Janssen, S., Shastri, G.G., Ilhan, Z.E., Challis, C., Schretter, C.E., Rocha, S., Gradinaru, V., Chesselet, M.F., Keshavarzian, A., Shannon, K.M., Krajmalnik-Brown, R., Wittung-Stafshede, P., Knight, R., Mazmanian, S.K., 2016. Gut Microbiota Regulate Motor Deficits and Neuroinflammation in a Model of Parkinson’s Disease. Cell 167, 1469–1480.e12. https://doi.org/10.1016/j.cell.2016.11.018

Schindelin, J., Arganda-Carreras, I., Frise, E., Kaynig, V., Longair, M., Pietzsch, T., Preibisch, S., Rueden, C., Saalfeld, S., Schmid, B., Tinevez, J.-Y., White, D.J., Hartenstein, V., Eliceiri, K., Tomancak, P., Cardona, A., 2012. Fiji: an open-source platform for biological-image analysis. Nat. Methods 9, 676–682. https://doi.org/10.1038/nmeth.2019

Staley, C., Kaiser, T., Beura, L.K., Hamilton, M.J., Weingarden, A.R., Bobr, A., Kang, J., Masopust, D., Sadowsky, M.J., Khoruts, A., 2017. Stable engraftment of human microbiota into mice with a single oral gavage following antibiotic conditioning. Microbiome 5, 87. https://doi.org/10.1186/s40168-017-0306-2

van den Berg, R.A., Hoefsloot, H.C.J., Westerhuis, J.A., Smilde, A.K., van der Werf, M.J., 2006. Centering, scaling, and transformations: improving the biological information content of metabolomics data. BMC Genomics 7, 142. https://doi.org/10.1186/1471-2164-7-142

Zunszain, P.A., Anacker, C., Cattaneo, A., Choudhury, S., Musaelyan, K., Myint, A.M., Thuret, S., Price, J., Pariante, C.M., 2012. Interleukin-1β: a new regulator of the kynurenine pathway affecting human hippocampal neurogenesis. Neuropsychopharmacol. Off. Publ. Am. Coll. Neuropsychopharmacol. 37, 939–949. https://doi.org/10.1038/npp.2011.277

